# The minor antennae of photosystem II contribute to qH-energy dissipation in *Arabidopsis*

**DOI:** 10.1101/2025.02.11.637767

**Authors:** Pierrick Bru, Aurélie Crepin, Yolande Provot, Zeno Guardini, Roberto Bassi, Luca Dall’Osto, Alizée Malnoë

**Affiliations:** Department of Plant Physiology, Umeå Plant Science Centre (UPSC), Umeå University, Umeå, Sweden; Department of Biology, Indiana University, Bloomington, IN, USA; Aix-Marseille University, CEA, CNRS, BIAM, LGBP Team, Marseille, France; Department of Biotechnology, University of Verona, Verona, Italy; Anton Dohrn Experimental Marine Station, Naples, Italy

**Author notes:** The author responsible for distribution of materials integral to the findings presented in this article in accordance with the policy described in the Instructions for Authors (https://academic.oup.com/plcell/pages/General-Instructions) is: Alizée Malnoë.

**Keywords:** photosynthesis, energy dissipation, nonphotochemical quenching qH, *Arabidopsis thaliana*, light-harvesting complexes, CRISPR–Cas9, abiotic stress

## Abstract

Photosynthesis is a biological process that converts light energy into chemical energy. Excessive light can damage the photosynthetic machinery, so plants have evolved photoprotective mechanisms such as non-photochemical quenching (NPQ). Among the NPQ mechanisms, qH is a form of sustained quenching, dependent on LIPOCALIN IN THE PLASTID (LCNP) and repressed by SUPPRESSOR OF QUENCHING 1 (SOQ1), protecting against abiotic stress. Recently, we showed in *Arabidopsis thaliana* that qH can occur in the major light-harvesting complexes (Lhcb1, Lhcb2, Lhcb3) but independently of any specific major antenna. Interestingly, in mutants with little or no accumulation of major antennae (*koLHCII, lhcb1, cpsrp43*), qH can still be induced. Here, we show that the minor antennae can be quenched by qH and remain quenched once isolated. To investigate the role of minor antennae in qH, we combined the *soq1* mutant, which displays high qH, with mutations in each minor antenna type (Lhcb4, Lhcb5, or Lhcb6), or with a mutant lacking all minor antennae. None are strictly required for qH to occur. Still, the absence of Lhcb6 decreases qH induction likely due to an indirect effect from the slower electron transport rate and/or a different macro-organization of photosynthetic complexes in the thylakoids. Overall, this work demonstrates that the minor antennae are a secondary target for qH and could serve as an additional safety valve for photoprotective energy dissipation during prolonged stress.

## Introduction

Plants rely on light energy to drive the photosynthetic machinery that synthesizes organic compounds through the Calvin-Benson cycle. However, excessive light can cause damage by generating reactive oxygen species (ROS), which, if unmanaged, can be lethal to the plant (Khorobrykh et al. 2020, Foyer and Hanke 2022). Plants have evolved strategies to mitigate light-induced damage, such as avoiding light absorption through leaf or chloroplast movement (Dwyer and Hangarter 2022) and dissipating excess light as heat in a process known as non-photochemical quenching (NPQ)(Horton et al. 1996). NPQ comprises several mechanisms, each identified by its specific molecular players and relaxation kinetics in the dark, labeled “q” (for quenching of chlorophyll fluorescence) followed by a letter denoting the mechanism (e.g., qE, qZ, qH)(Li et al. 2009, Bassi and Dall’Osto 2021). In *Arabidopsis thaliana*, we investigated a sustained NPQ mechanism called qH, which forms and relaxes over hours and protects plants from severe photooxidative stress associated with critically adverse conditions such as drought or cold and high light (Malnoë 2018). qH requires the lumenal protein LIPOCALIN IN THE PLASTID (LCNP), is repressed by the thylakoid membrane protein SUPPRESSOR OF QUENCHING 1 (SOQ1) and is relaxed by the stromal protein RELAXATION OF QH1 (ROQH1) (Brooks et al. 2013, Malnoë et al. 2017, Amstutz et al. 2020). Upon light absorption by chlorophyll, excitation energy can be used for photochemistry, dissipated as heat (NPQ), or reemitted as fluorescence (Müller et al. 2001). An inverse relationship between amplitudes of NPQ and chlorophyll fluorescence, observable when photochemistry is saturated by a strong light pulse, allows NPQ assessment (Brooks and Niyogi 2011). Because biomass and seed yield are linked with light energy capture and usage efficiency, understanding NPQ to improve these properties in specific conditions (e.g. shade to light transition) can help improve crop production (Zhu et al. 2004, Kromdijk et al. 2016, Wang et al. 2020, De Souza et al. 2022, Long et al. 2022, Khaipho-Burch et al. 2023).

Photosystem II (PSII) is composed of a dimeric reaction center surrounded by light-harvesting complexes (LHCs). LHCs include major antennae (Lhcb1, 2, 3) and minor antennae (Lhcb4/CP29, Lhcb5/CP26, Lhcb6/CP24) associated with the PSII core. Major antennae can form homo- or hetero-trimeric complexes associated to the PSII core directly in a strongly bound manner (S-LHCII), or moderately (M-LHCII) associated through Lhcb4 and 6, or loosely (L-LHCII) associated through other trimers, while minor antennae are present in a monomeric form connected to the PSII core (Ballottari et al. 2012, Su et al. 2017). LHC proteins mediate the efficient transfer of absorbed light energy to photosystems, ensuring balanced energy excitation between PSI and PSII. They are also crucial for photoprotection, helping to minimize photooxidative damage (Iwai et al. 2024). For qE, the monomeric antennae contribute to the early phase of quenching followed by quenching in the trimeric antennae (Dall’Osto et al. 2017). In *Arabidopsis thaliana*, Lhcb1, 2, and 4 are the protein products of five, three, and three genes respectively, whereas Lhcb3, 5, and 6 are each encoded by one gene. Although these isoforms exhibit slight variations in pigment composition, they all bind chlorophyll *b*, which is absent in the core complexes (Caffarri et al. 2014). While the different antennae collectively contribute to light harvesting, different subunits also participate in specific functions in regulating photosynthesis (Caffarri et al. 2004, Crepin and Caffarri 2018). As the land plants-specific Lhcb3 and Lhcb6 interact and promote M-LHCII binding (Caffarri et al. 2009, Su et al. 2017), the regulation of their expression can participate in adjusting PSII antenna size under excess light conditions (Ballottari et al. 2007, Albanese et al. 2016); specific phosphorylation of Lhcb2 mediates the binding of trimeric LHCII to PSI core, balancing excitation of both photosystems (Crepin and Caffarri 2015, Longoni et al. 2015, Pan et al. 2018, Cutolo et al. 2023a); Lhcb5 is involved in NPQ reactions likely through zeaxanthin binding (Dall’Osto et al. 2005), while a shorter isoform of Lhcb4, Lhcb4.3 (also referred to as Lhcb8) is upregulated in excess light and possibly participates in photoacclimation (Albanese et al. 2019, Opatikova et al. 2023). Antennae are imported and inserted into the thylakoid membranes via the signal recognition particle (SRP) pathway (Figure 1A), where the chaperones cpSRP43 and cpSRP54 play an important role (Ziehe et al. 2018); the absence of either protein leads to a decreased antennae accumulation (Hutin et al. 2002, Jeong et al. 2017).

**Figure 1.**
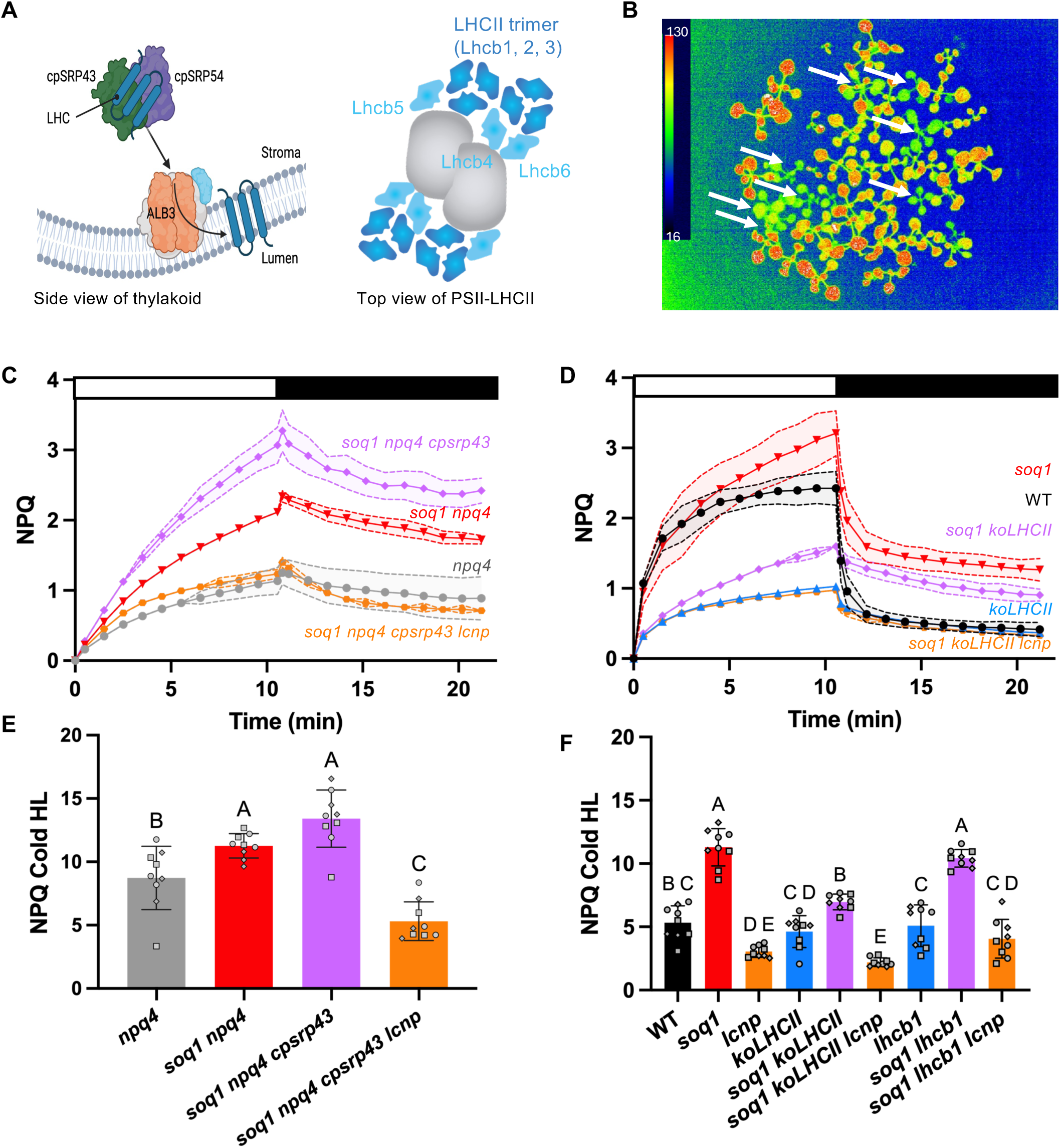
The *soq1* mutation results in a high NPQ phenotype in mutants with fewer or no major antennae. (A) Schematic model of the SRP pathway of antenna import and insertion in the thylakoids, involving the cpSRP43 protein, adapted from (Ziehe et al. 2018). The full PSII complexes comprising a dimeric core, all minor antennae and four LHCII trimers is also displayed. (B) F2 *soq1 npq4* x E40 minimum fluorescence measurement. Chlorophyll fluorescence is shown in false color: blue corresponds to lower fluorescence and red to higher fluorescence. Seedlings pointed by a white arrow with a low F_0_ have been harvested for DNA extraction. (C, D) NPQ kinetics of *soq1 npq4 cpsrp43 or soq1 koLHCII* (purple diamond), *koLHCII* (blue up-pointing triangle), *soq1 npq4 cpsrp43 lcnp* or *soq1 koLHCII lcnp* (orange hexagon), *soq1* or *soq1 npq4* (red down-pointing triangle), WT or *npq4* (black or gray circle). NPQ was induced for 10 min at 1,200 µmol photons m^-2^ s^-1^ (white bar) and then relaxed in the dark for 10 min (black bar). (E, F) NPQ measurements after a 6 h cold treatment followed by 8 h of cold and high light (Cold HL) for the same mutants shown in panels C and D, respectively. All measurements were taken from 5 (E,F) or 20 (C, D) min dark-adapted detached leaves from 6-week-old plants. Data are presented as mean ± SD (n = 3 detached leaves from independent individuals). The results display a representative experiment, or all data, from three independent biological replicates, denoted by different symbols in (E, F). Statistical analyses using ANOVA and Tukey’s multiple comparison test with a 95% confidence interval, reveals a significant increase in NPQ level for *soq1 npq4 cpsrp43, soq1 koLHCII* and *soq1 lhcb1* compared to *soq1 npq4 cpsrp43 lcnp, soq1 koLHCII lcnp* and *soq1 lhcb1 lcnp*, respectively (*p* < 0.05).

Previously, we reported that chlorophyll *b* is required for qH indicating its location in the peripheral antennae (Malnoë et al. 2017) and we evidenced that qH can occur in the LHCII trimer (Bru et al. 2022). Unexpectedly, we noticed that mutants with less LHCII in a *soq1* mutant background displayed enhanced qH compared to *soq1* (*e.g., soq1 lhcb1* (Bru et al. 2022), *soq1 npq4 cpsrp43* (*this study*)). This prompted us to further examine the role of the PSII antennae for qH. Here, we generated a *soq1 koLHCII* line devoid of trimeric antennae and found that qH can still be induced. Through isolation of photosynthetic complexes and chlorophyll fluorescence measurements, together with CRISPR-Cas9-mediated genome editing of *LCNP*, we demonstrate that qH can occur in the minor antennae. This finding is significant as it provides stable, natively quenched monomeric antenna directly isolated from plants without artificial treatment. To further investigate the role of minor antennae in qH, we generated the *soq1 NoMinor (NoM)*, *soq1 lhcb4*, *soq1 lhcb5*, and *soq1 lhcb6* mutants. We found that qH is independent of Lhcb4, Lhcb5, and Lhcb6; however, the absence of Lhcb6 appears to slightly decrease qH induction, likely due to an indirect effect on electron transport and/or a different antennae organization (de Bianchi et al. 2008, Guardini et al. 2022a). The minor antennae, therefore, contribute to qH under specific conditions and modulate the full extent of qH but are not strictly required for qH. Of note, our results show that qH in the minor antennae provides photoprotection.

## Results

### A new mutant allele of cpSRP43 results in higher NPQ in the soq1 background

The mutant named “E40” was isolated from an ethyl methanesulfonate (EMS)-mutagenized *Arabidopsis soq1 npq4* M2 population described in (Bru et al. 2020) and displayed lower chlorophyll accumulation, lower maximum, and minimum, chlorophyll fluorescence F_m_, and F_o_, respectively and higher NPQ compared to the parental line (Figure 1B,C, S1A). To identify the causative mutation for the phenotype, we used a mapping-by-sequencing approach. The mutant E40 was backcrossed to the parental line *soq1 npq4* and individuals that exhibited a low F_0_ from the F2 progeny (which represent one-quarter of the individuals with genotype *soq1 npq4* and homozygous for the new mutation) were selected. Genomic DNA was extracted from the selected F2 seedlings and subjected to whole genome sequencing. The sequencing reads were mapped onto the *Arabidopsis* reference genome and single nucleotide polymorphisms were identified. The position and frequency of single nucleotide polymorphisms were plotted and an increase in the allelic frequency of mutations approaching 100% was observed in the region between 15 and 20 Mb on chromosome 2, identifying this region as the one containing the causative mutation (Figure S2A). Only one mutated gene encoding for a protein targeted to the chloroplast was mutated in that region corresponding to *cpSRP43*. Nucleotide transitions G to A in position 873 resulted in an early stop cpSRP43-Trp291*. As other *cpsrp43* alleles have been previously described in *Arabidopsis*, the *chaos* mutant and *cpsrp43-2* (Amin et al. 1999, Klimyuk et al. 1999, Anderson et al. 2021), we named this new allele *cpsrp43-3* (hereafter referred to as *cpsrp43* for clarity). We examined by immunoblot analysis cpSRP43 protein accumulation using an antibody raised against the recombinant protein derived from *Arabidopsis thaliana* and found no accumulation of cpSRP43 in the mutant E40 (*soq1 npq4 cpsrp43*) (Figure S1B). In wild type (WT), we observed two bands, one band at the apparent size of 43 kDa for an expected size of 35.1 kDa and a higher band (at 80 kDa) which could be a dimer of cpSRP43 as this band is also lacking in the mutant. This new *cpsrp43* mutation is, therefore, a knock-out allele and results in a lower level of trimeric antennae compared to WT (Figure S2B) as previously found (Amin et al. 1999).

### qH can occur in mutants with fewer or no major antennae accumulation

The higher NPQ phenotype of *soq1 npq4 cpsrp43* (compared to *soq1 npq4*) indicates that qH may be enhanced although fewer major antennae are present, reminiscent of what we observed in the *soq1 lhcb1* line (Bru et al. 2022). Indeed, using the CRISPR-Cas9 system, we generated a *soq1 npq4 cpsrp43 lcnp* line that displayed a low level of NPQ similar to the original *npq4* mutant (Figure 1, S1A) so the enhanced NPQ is qH (i.e., LCNP-dependent). As trimeric antennae level is lower in the *cpsrp43* mutant and given that we previously identified the trimeric LHCII as a target of qH (Bru et al. 2022), we tested whether qH could occur without any of the major antennae. We combined the *koLHCII* line lacking all the major antennae (Guardini et al. 2022b) to the *soq1* mutation to generate *soq1 koLHCII* and also generated a *soq1 koLHCII lcnp* line (Figure S1C, D). Intriguingly, the *soq1 koLHCII* exhibited additional NPQ compared to *koLHCII*, in an LCNP-dependent manner (Figure 1D). All throughout, we use the term biological replicate to refer to one batch of several plant individuals grown in the same conditions at the same time; at minimum we acquire data from three independent individuals of a given genotype in three independent biological replicates.

While qH can be induced upon 10 minutes of high-light exposure in the *soq1* background, we use the combination of stress conditions of cold (4°C) and high light (1,600 µmol photons m^-2^ s^-1^) to induce qH in WT. We exposed detached leaves to 6h cold followed by an 8h cold and high-light treatment. Similarly to the short NPQ kinetic induction, the mutants *soq1 npq4 cpsrp43*, *soq1 lhcb1* and *soq1 koLHCII*, display additional LCNP-dependent NPQ compared to control lines (Figure 1E, F, S1E, F) (note that the NPQ Cold HL of *soq1 koLHCII* is higher than *koLHCII* but lower than *soq1* – Figure 1F). Therefore, qH can be fully or partially activated with fewer or no major antennae accumulation. These results suggest that the major antennae are not essential for qH activation and that qH can occur elsewhere in the antenna pool.

### Minor antennae can be quenched by qH when major antennae are absent or less accumulated

To understand the origin of qH in the mutants depleted in antennae, we treated plants with cold and high light conditions, isolated, solubilized thylakoid and separated photosynthetic complexes by clear native-PAGE followed by chlorophyll fluorescence measurement (Figure 2A). We then quantified the fluorescence of the LHCII trimer band and Lhcb monomers for each mutant relative to its control lacking LCNP (qH off). In the mutants retaining LHCII trimers (*soq1 lhcb1* and *soq1 npq4 cpsrp43*) we observed a ∼25% decrease in fluorescence yield (Figure 2B) consistent with quenched LHCII by qH described in (Bru et al. 2022). In addition, here we observed a ∼25% decrease in fluorescence in the Lhcb monomer band of all three mutants (Figure 2C).

**Figure 2.**
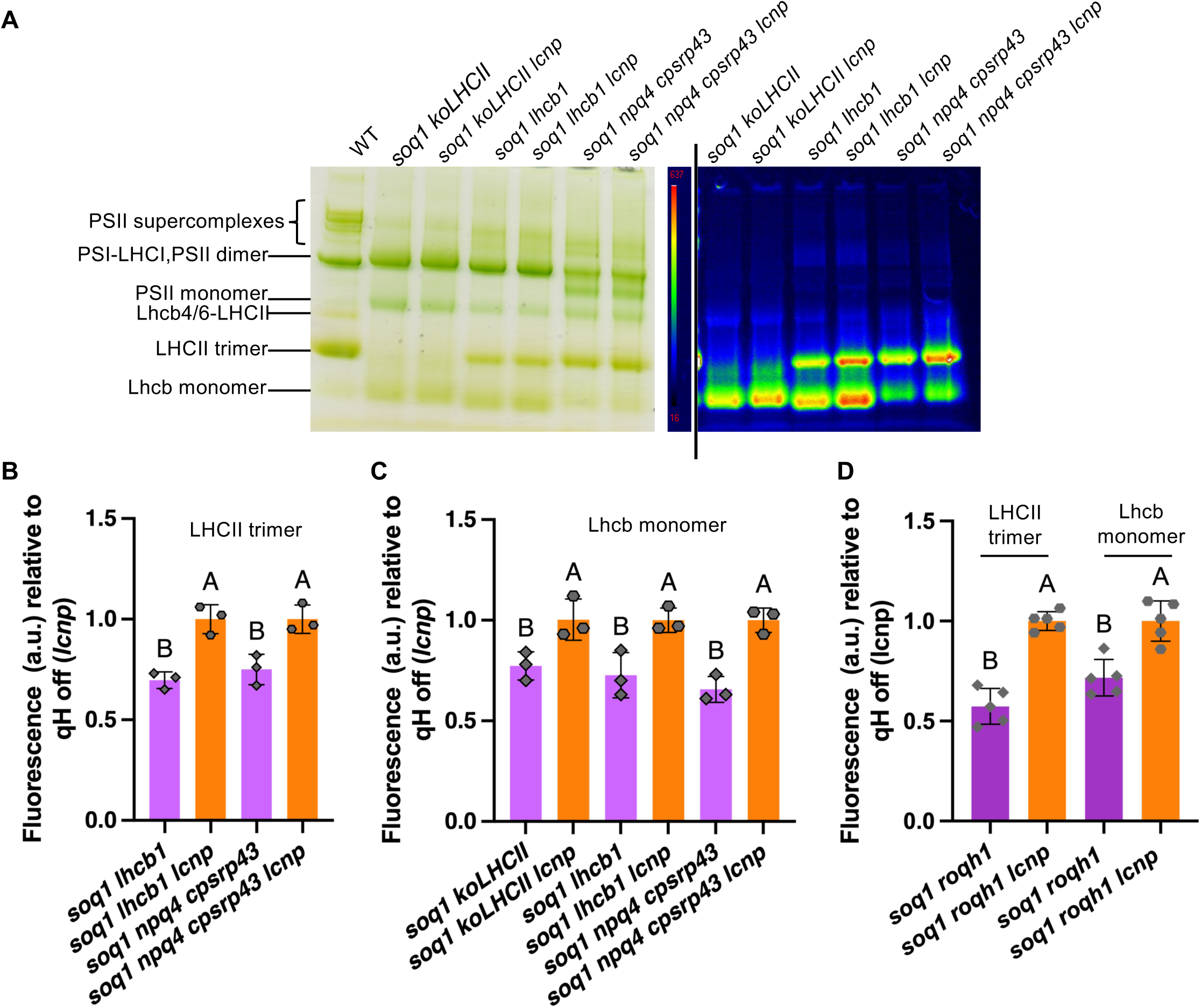
qH can be induced in the minor antennae. (A) from left to right: thylakoids corresponding to 10 µg of chlorophyll were loaded per lane for WT, *soq1 koLHCII, soq1 koLHCII lcnp, soq1 lhcb1, soq1 lhcb1 lcnp, soq1 npq4 cpsrp43* and *soq1 npq4 cpsrp43 lcnp.* Thylakoid obtained from plants exposed to 6h cold treatment followed by 8h cold and HL were solubilized to a final concentration of 0.5 mg Chl mL^-1^ with 1% α-DM, pigment-protein complexes were separated on 4-16% clear native PAGE, with the chlorophyll fluorescence of the same gel shown. This is one representative experiment from n=3 biological replicates. The additional band in *soq1 npq4 cpsrp43*, between PSII monomer and PSII dimer, likely represents the co-migration of PSI core with PSII lacking its antenna (Bujaldon et al. 2020). (B and C) Quantification of chlorophyll fluorescence for the trimers (B) and monomers (C) band, relative to corresponding mutant with qH off (*lcnp*), normalized by the intensity of the corresponding green band from the clear native gel (n=3 biological replicates). (D) Chlorophyll fluorescence quantification of trimers and monomers isolated by FPLC from the non-stressed *soq1 roqh1* mutant, normalized by the maximum optical density in the Qy peak for each sample (n=5 biological replicates). Statistical analyses were made using ANOVA with Tukey’s multiple comparison test and a 95% confidence interval, revealing a significant decrease in chlorophyll fluorescence yield in the trimeric or monomeric antennae of *soq1 lhcb1, soq1 npq4 cpsrp43*, *soq1 koLHCII* and *soq1 roqh1* compared to *soq1 lhcb1 lcnp, soq1 npq4 cpsrp43 lcnp*, *soq1 koLHCII lcnp* and *soq1 roqh1 lcnp* respectively (*p* < 0.05).

We qualitatively analyzed the trimeric and monomeric bands by SDS-PAGE and found a similar antennae composition between the qH-active samples and qH-inactive ones (*lcnp* mutant background) so the difference in chlorophyll fluorescence is not due to a different antennae composition (Figure S3). In the *soq1 lhcb1* and *soq1 lhcb1 lcnp* monomeric band we observed some major antennae (band marked as Lhcb1+2, in this case containing Lhcb2) possibly stemming from monomerization of the trimer antennae during the solubilization process. However, this is not the case for *soq1 npq4 cpsrp43* or *soq1 koLHCII* (as this mutant lacks the major antenna), so the decrease of chlorophyll fluorescence in the Lhcb monomer band does not originate from LHCII monomerization, implying that the minor antennae can be quenched by qH. If we plot the chlorophyll fluorescence normalized to the intensity levels of the corresponding band, we consistently observed a decreased fluorescence in a LCNP-dependent manner, indicative of quenching by qH. We noted however that the fluorescence yield of the trimeric fraction(s) from *soq1 npq4 cpsrp43* (*lcnp*) is overall lower than that of *soq1 lhcb1* (*lcnp*) and in contrast that the fluorescence of the monomeric fraction(s) from *soq1 npq4 cpsrp43* (*lcnp*) is overall higher than that of *soq1 lhcb1* (*lcnp*) and *soq1 koLHCII* (*lcnp*) (Figure S3C). These varying properties are likely due to a different Lhcb subunits and/or isoforms composition, as they do not possess the same chlorophyll content (Caffarri et al. 2014, Su et al. 2017) of the trimer and monomer fractions between the antennae mutants.

To verify if the quenching of monomers occurs specifically upon depletion of the major LHCII or if monomeric antennae can also be quenched when all Lhcb are present, we measured the fluorescence of the purified complexes from the non-stressed *soq1 roqh1* mutant, which is constitutively qH-quenched (Amstutz et al. 2020). Both trimeric and monomeric antennae isolated from this mutant displayed lower fluorescence compared to the *soq1 roqh1 lcnp* control (Figure 2D), suggesting that the minor antennae, unquenched in the presence of a wild-type level of trimers after a 6h cold and high light treatment (Bru et al. 2022), can be quenched if qH is activated in a stronger and more lasting manner.

### qH does not rely on a specific minor Lhcb

Now that we established that qH can occur in the minor antennae, we tested their requirement for qH. To that end, we generated a knock-out of *SOQ1* using CRISPR-Cas9 in the *No Minor* (*NoM*) mutant background, as well as in the *lhcb4* mutant and in the *lhcb6* mutant background. The *soq1 lhcb5* was obtained by crossing *soq1-1* with *lhcb5*. We confirmed by SDS-PAGE, followed by immunoblot analysis of thylakoid extract, that the expected Lhcb(s) and SOQ1 did not accumulate (Figure S4A). The absence of Lhcb4 results in the degradation of Lhcb6 (de Bianchi et al. 2011). A small amount of Lhcb6, therefore, was still detectable in the *NoM* and *lhcb4* context as well as in the T-DNA *lhcb6* mutant line (affecting *LHCB6* 5’UTR) as described (Ilikova et al. 2021). To ascertain the role of Lhcb6 in qH, we generated a “true” knock-out of *LHCB6* using CRISPR-Cas9 in the *soq1-1* background. In this new line named *soq1 lhcb6-2*, we can observe no residual Lhcb6 and that all the other antennae accumulate (Figure S4B).

We then proceeded to characterize their NPQ phenotype. Similar F_m_ between the *lhcb* and *soq1 lhcb* mutants enabled the comparison of their NPQ kinetics and level after a cold and high-light treatment (Figure S5). The *soq1 NoM*, *soq1 lhcb4*, *soq1 lhcb5* and *soq1 lhcb6* displayed a higher NPQ than their respective control (Figure 3, S6). The full extent of qH after a cold and high-light treatment was not reached in the *soq1 lhcb6* mutant (Figure 3E), which prompted us to investigate the quenching levels in isolated antennae. Quenching indeed took place in LHCII trimers (Figure 4), indicating that Lhcb6 is not strictly required for qH. These results show that qH can occur in the absence of the minor antennae and does not rely on a specific minor Lhcb. We noted that the NPQ Cold HL level of the *lhcb5* mutant was low (similar to *lcnp*) and this may be due to less qZ induction (Dall’Osto et al. 2005) as qH is otherwise properly induced in *soq1 lhcb5*.

**Figure 3.**
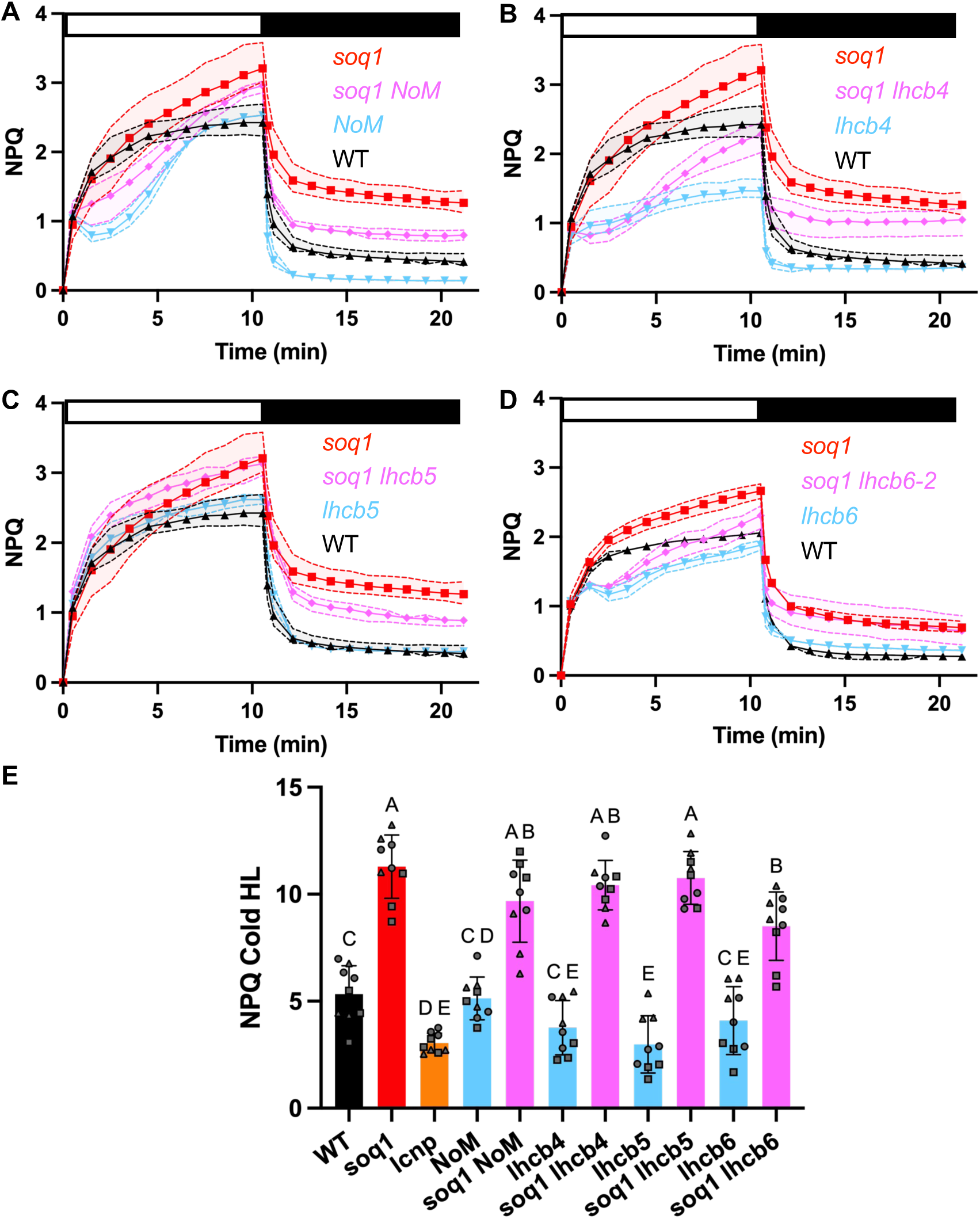
The *soq1* mutation causes a high NPQ phenotype in minor antennae mutants. (A, B, C, D) NPQ kinetics of WT (black up-pointing triangle), *soq1* (red square), *lhcb4, 5* or *6* and *NoM* (blue down-pointing triangle), *soq1 lhcb4, 5* or *6* and *soq1 NoM* (pink diamond). NPQ was induced for 10 min at 1,200 µmol photons m^-2^ s^-1^ (white bar) and relaxed in the dark for an additional 10 min (black bar). For D, similar results were obtained with *soq1 lhcb6 #3, 7* (Figure S6C). (E) Bar graph representing NPQ after 6 h cold treatment followed by 8 h cold and high light (Cold HL). *soq1 lhcb6 #10* was used here, similar results were obtained with *soq1 lhcb6-2* (Figure S6D). All measurements were taken from 5 (E) or 20 (A,B,C,D) min dark-adapted detached leaves from 6-week-old plants. Data are presented as mean ± SD (n = 3 detached leaves from independent individuals). The results display a representative experiment, or all data, from three independent biological replicates, denoted by different symbols in (E). Statistical analyses were made using ANOVA with Tukey’s multiple comparison test and a 95% confidence interval, revealing a significant increase in NPQ levels for *soq1 NoM*, *soq1 lhcb4*, *soq1 lhcb5* and *soq1 lhcb6* compared to *NoM, lhcb4, lhcb5* and *lhcb6*, respectively (*p* < 0.05). Additionally, there was a significant decrease in NPQ level for *soq1 lhcb6* compared to *soq1* (*p* <0.05).

**Figure 4.**
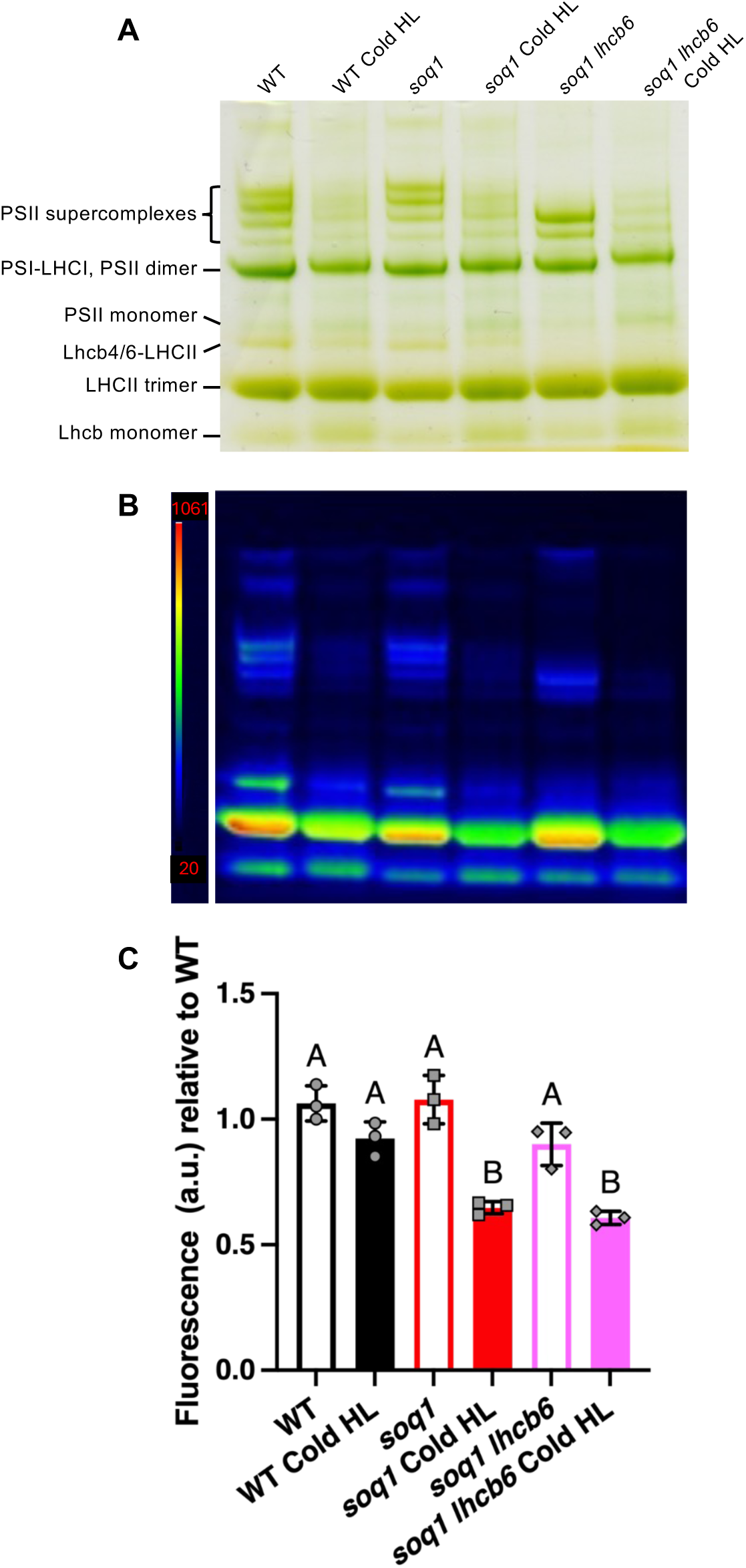
Isolated LHCII trimers from *soq1 lhcb6* are quenched upon stress treatment. (A, B) Thylakoid (10 µg chlorophyll) from WT, *soq1* and *soq1 lhcb6* plants, exposed to a 6h cold followed by 8h cold and high light treatment, were solubilized to a final concentration of 0.5 mg Chl mL^-1^ and 1% α-DM, then separated on 4-16% clear native PAGE. Gel image (A) and chlorophyll fluorescence image (B). (C) Chlorophyll fluorescence quantification of the trimeric LHCII band, relative to the average of three WT dark-adapted replicates, normalized by the corresponding green band from the gel. Data are presented as mean ± SD (n=3 biological replicates i.e. independent thylakoid preparations from independently grown and treated plants).

### qH in the minor antennae is photoprotective

To test the photoprotective role of qH quenching in the minor antennae, we exposed detached leaves from the *koLHCII* mutant in both the *soq1* (qH-active in the minor antennae) and *soq1 lcnp* (qH-inactive) backgrounds, to a cold and high light conditions for up to 45 hours. We monitored the visual phenotype as well as the leaf chlorophyll content over time. It was observed that the leaves of *soq1 koLHCII lcnp* fully bleached compared to those of *koLHCII* or *soq1 koLHCII* (Figure 5, S7). Additionally, *soq1 koLHCII* retained its green color longer than *koLHCII.* These results clearly indicate that qH in the minor antennae provides photoprotection by preventing chlorophyll damage and degradation.

**Figure 5.**
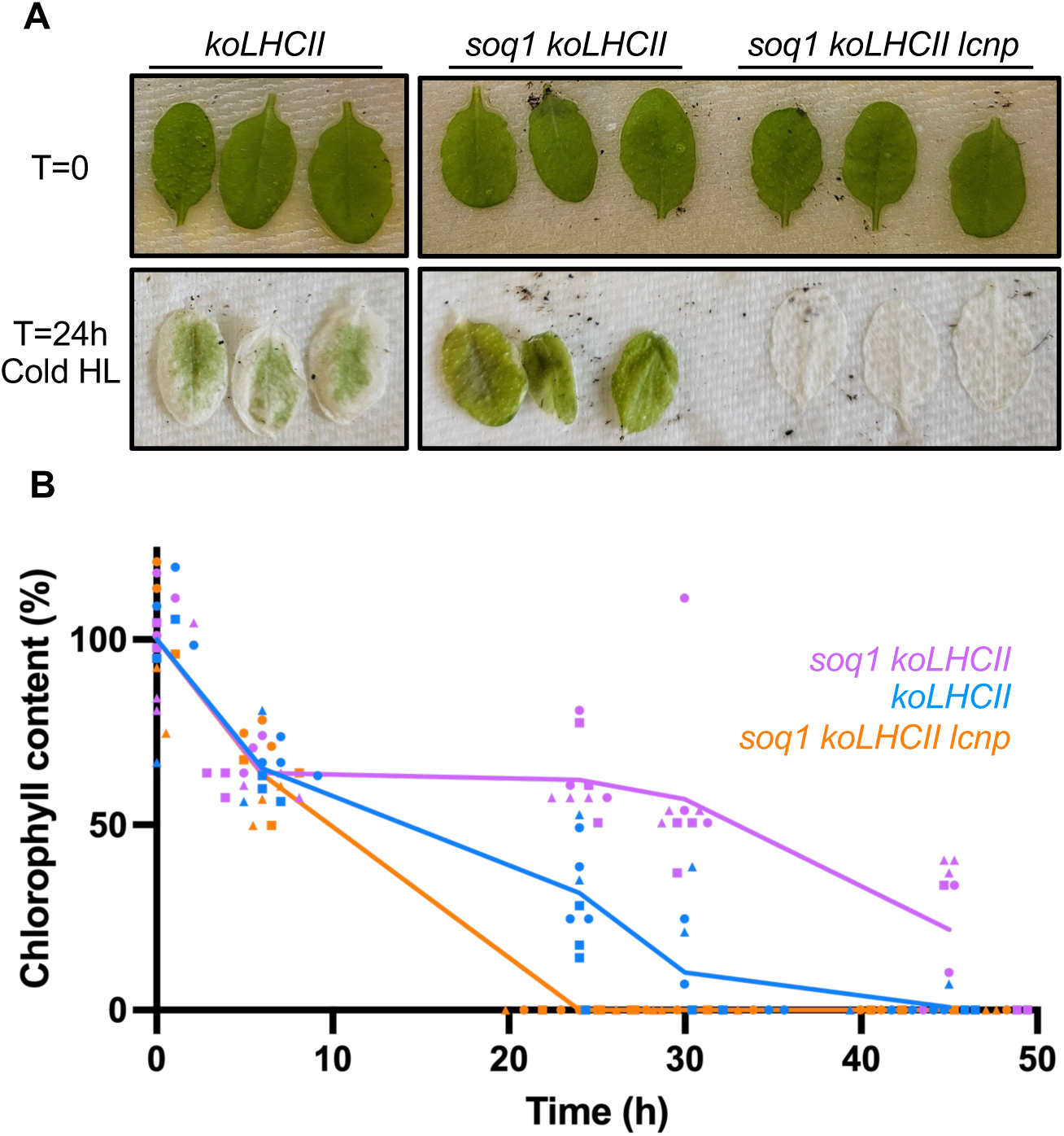
qH in the minor antennae provides photoprotection. (A) Representative image of detached leaves from *koLHCII, soq1 koLHCII, soq1 koLHCII lcnp* before (top) and after a 24h cold and high light (Cold HL) stress (bottom). (B) Leaf chlorophyll content was monitored throughout the cold and high light exposure for 45 h. Data are presented as mean (n = 9, three detached leaves from independent individuals from three independent biological replicates, denoted by different symbols). For each plant line, values are normalized on the average chlorophyll content at the start of the experiment.

## Discussion

Our findings reveal that qH can occur in the minor antennae and that this quenching is photoprotective. In the same line to our previous work on isolated quenched LHCII, the successful isolation, here, of natively quenched minor antennae by qH paves the way for revealing its molecular origin.

### The minor antennae are a secondary site for qH

The enhanced NPQ previously observed in the *soq1 lhcb1* and *soq1 npq4 cpsrp43* can now be explained by qH occurring in both the trimeric and monomeric antennae (Figure 2). The antennae organization and accumulation in these mutants may result in better access for LCNP to induce qH at both sites. Considering that we did not observe quenching in the monomeric Lhcb of *soq1* plants (Bru et al. 2022), but we found it in the non-stressed, constitutively quenched *soq1 roqh1* (Figure 2D), qH may initially target trimeric antennae (primary site) before extending to monomeric antennae (secondary site) under severe, prolonged stress. Sequential activation of quenching in trimeric and monomeric antennae has been reported before (Holzwarth et al. 2009, Dall’Osto et al. 2017). In these cases, interestingly, both types of antennae were subjected to different mechanisms, that could be attributed to different NPQ components or qE sub-components. Here, it is not known whether the origin of the quenching is similar in the major and minor antennae, however both are dependent on the LCNP protein. Structural and ultrafast spectroscopy studies will provide future insights into potential differences in the qH mechanism(s) of major and minor antennae. The minor antenna quenching could serve as an additional safeguard during prolonged stress, complementing the protective role of major antenna quenching (Figure 6).

**Figure 6.**
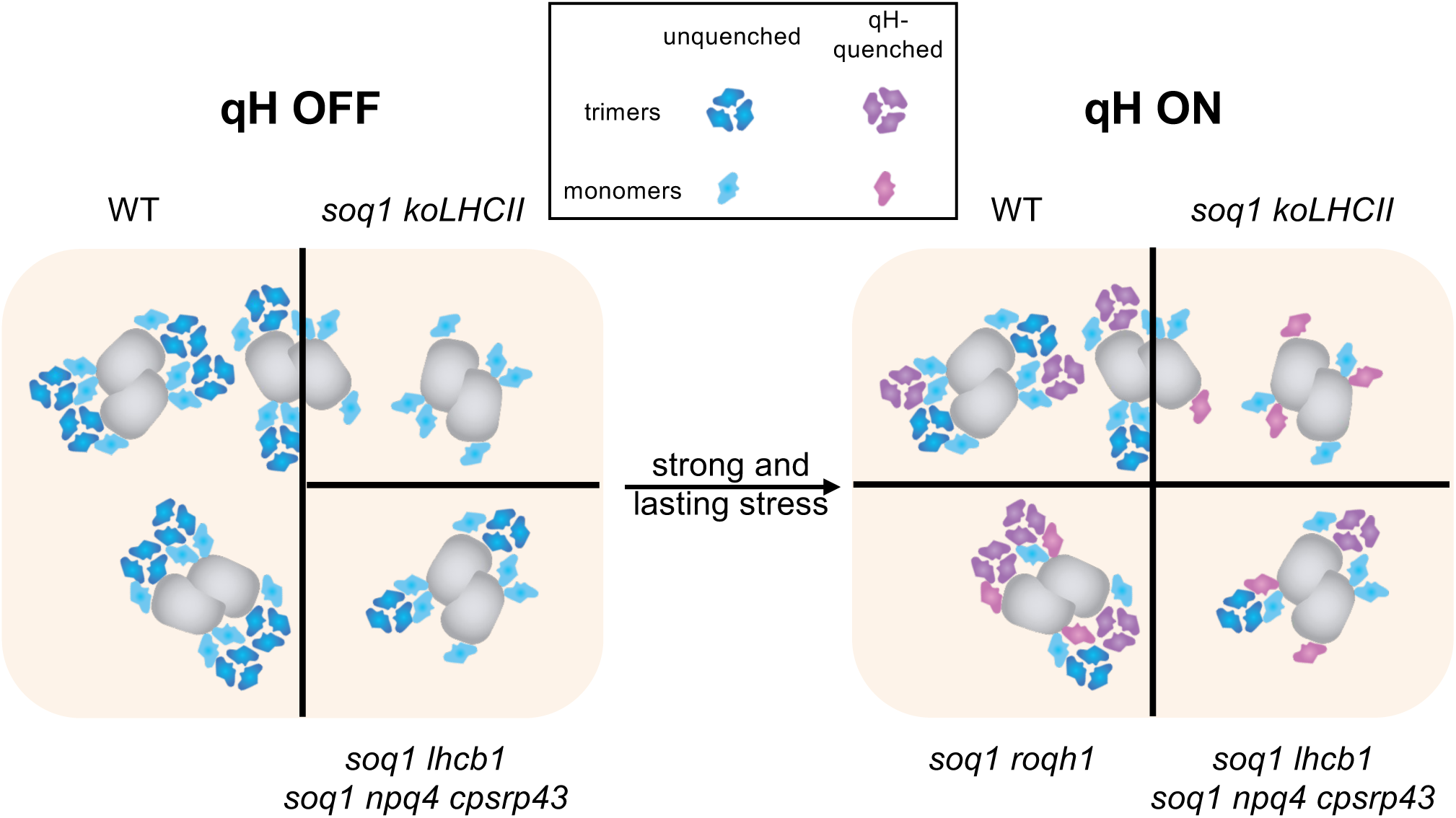
Model of qH in the antennae systems of the mutants presented in this study. Mutants containing varying amounts of antennae (left) display a different repartition of qH quenching after a strong and lasting stress exposure, such as several hours of cold and high light (right). In plants containing a complete antenna system, qH is only discernable in the major trimeric antennae, unless the stress (and therefore qH activation) is longer-lasting, such as in the qH-constitutive *soq1 roqh1* plants: minor monomeric antennae are then also affected. In mutants lacking trimeric antennae (*soq1 koLHCII*) or with lower amount (*soq1 lhcb1, soq1 npq4 cpsrp43*), both monomeric antennae and remaining trimers can be quenched by qH.

### Implications for the qH mechanism

In contrast to previously described by (Sattari Vayghan et al. 2022), we observed some LHCII trimer accumulation in the *lhcb1* mutant background (Figure 2, S3A). Different thylakoid membrane solubilization methods (α-DM (this study) vs. digitonin (prior study)) may have enabled to preserve a less stable LHCII trimer; indeed, digitonin can occupy the centre of the LHCII trimer (Graca et al. 2021) thus potentially leading to its monomerization. The remaining trimer are possibly formed of Lhcb2 homotrimer and/or Lhcb2 and Lhcb3 heterotrimer as Lhcb3 cannot form homotrimer (Pietrzykowska et al. 2014, Cutolo et al. 2023a). This result indicates that qH can occur in a homotrimer of Lhcb2. However, in the mutant lacking all minor antennae (*NoM*), despite containing 60% more trimeric antennae (Dall’Osto et al. 2020), the mutant *soq1 NoM* did not show an increased capacity to induce qH compared to *soq1* (Figure 3A). This suggests that a connection between the antennae and the PSII core would be required for effective qH induction.

### Indirect effect of Lhcb6 mutation on qH

We demonstrated that qH is independent of a specific monomeric (Figure 3) or trimeric Lhcb (Bru et al. 2022) protein, highlighting its flexibility. The flexibility of qH in its targets is particularly interesting in engineering plants with decreased antenna size for increased light distribution within the canopy or microalgal cultures (Kirst et al. 2017, Cutolo et al. 2023b). It points to photoprotection of these plants under harsh stress being possibly minimally impaired as they will still be able to perform qH. Although qH does not rely on a specific monomeric antenna, Lhcb6 appeared to modulate its induction. The *soq1 lhcb6* mutant uniquely displayed a slightly lower NPQ level than *soq1* after a cold and high-light treatment (Figure 3E). *lhcb6* presents a slower photosynthetic electron transport rate due to restricted plastoquinone diffusion (de Bianchi et al. 2008, Ilikova et al. 2021) in contrast to the *lhcb4* mutant lacking both Lhcb4 and Lhcb6, where the electron transport rates are not altered (de Bianchi et al. 2011). The slight NPQ decrease in the mutants *soq1 lhcb6,* not observed in the *soq1 lhcb4*, may therefore be explained by the slower electron transport rate possibly impacting LCNP activation (through e.g. lower oxygen evolution and ROS formed) (Yu et al. 2022, Hao et al. 2024). Alternatively, a different macro-organization and/or interconnection of PSII-LHCII units in the *lhcb6* mutant could limit the action of the qH-quenched trimers in the antenna pool, thereby reducing the overall extent of qH. (Ferguson et al. 2023) recently identified Lhcb6 as a potential candidate for allelic variation in the NPQ response of maize and this may be due to the lack of Lhcb6 having an indirect effect on modulating qH. This suggests that Lhcb6 may contribute to optimizing qH under specific stress conditions, although it is not strictly required.

### Downregulation of Lhcb6 to regulate qH in high light

Interestingly, excess light acclimation in *Arabidopsis* plants is associated with downregulation of Lhcb3 and Lhcb6 (Bailey et al. 2001, Ballottari et al. 2007, Floris et al. 2013, Kouril et al. 2013), resulting in a smaller PSII antenna size mainly caused by a decrease in the amount of M-trimer (Kovacs et al. 2006, Ballottari et al. 2007, Kouril et al. 2013, Flannery et al. 2021). During long-term stress conditions, multiple adaptation mechanisms are put in place such as a decrease of the antennae cross-section or a faster activation rate of qE (Ballottari et al. 2007, Kouril et al. 2013, Albanese et al. 2016) for photoprotection and optimization of light harvesting. The downregulation of Lhcb6 could limit qH activation, and this may constitute an adaptive strategy to prevent excessive energy dissipation, that would otherwise impair plant growth once other long-term acclimation mechanisms are established. Lhcb6 and Lhcb3 are absent in Norway spruce, resulting in unique PSII supercomplexes and megacomplexes organization (Kouril et al. 2020, Ilikova et al. 2021). In Norway spruce, the M-trimer associates with the PSII core complex in a different orientation than in *Arabidopsis* due to the lack of Lhcb6 and is closer to the S-trimer (Kouril et al. 2016). Similar observation has been made on *Chlamydomonas reinhardtii,* which also lacks Lhcb3 and Lhcb6 (Tokutsu et al. 2012, Drop et al. 2014, Kouril et al. 2016). The absence of Lhcb6 through its impact on LCNP access to its targets and/or its alteration of antenna interconnection may partly explain the diversity in PSII antenna organization affecting qH extent and regulation under prolonged stress. This is particularly noteworthy given that evergreens are more frequently exposed to severe stress during winter. Given the flexibility of qH in its Lhcb targets, it is thus likely that it can occur in non-vascular plants and evergreens, and exploring its conservation and regulation will be the subject of future work.

This study establishes qH as a flexible and adaptable photoprotective mechanism that can target both major and minor antennae under stress. Its independence from specific antennae proteins underscores its robustness, making it an attractive target for enhancing stress resilience in crops with altered light-harvesting properties. Future structural and spectroscopic studies will be crucial for unraveling the mechanistic details of qH quenching across diverse antenna configurations.

## Materials and Methods

### Plant material and growth conditions

Wild-type *Arabidopsis thaliana* and the derived mutants studied here are of the Col-0 ecotype. The following antennae mutants were used: *cpsrp43-3* (this study, in a *soq1-1 npq4-1 gl1* background), penta mutant *lhcb1.1.2.3.4.5*, and *lhcb4.1.2.3* are referred for clarity as *lhcb1* (Sattari Vayghan et al. 2022), and *lhcb4* (Betterle et al. 2009, de Bianchi et al. 2011) respectively, *lhcb3* (SALK_036200C) (Damkjaer et al. 2009), *lhcb5* (SALK_014869) (Kim et al. 2009), *lhcb6* (SALK_077953) (Kovacs et al. 2006, de Bianchi et al. 2008), *No Minor* (NoM) (Dall’Osto et al. 2014), *koLHCII* (Guardini et al. 2022b). In addition the mutants *soq1-1* (Brooks et al. 2013), *lcnp-1* (Malnoë et al. 2017) were used. Double mutants *soq1 lhcb1, 2, 3, 5* were obtained by crossing *soq1* with the corresponding *lhcb* mutant. Mutants *soq1 koLHCII, soq1 lhcb4, soq1 lhcb6, soq1 NoM* were obtained by *SOQ1* gene mutation using clustered regularly interspaced short palindromic repeats (CRISPR)-associated protein 9 (Cas9). Mutants *koLHCII lcnp, soq1 lhcb1 lcnp, soq1 npq4 cpsrp43 lcnp* were obtained by *LCNP* gene mutation using CRISPR-Cas9. Mutant *soq1 lhcb6-2* was obtained by *LHCB6* gene mutation using CRISPR-Cas9 in *soq1-1*. Finally, the mutant *soq1 koLHCII lcnp* has been obtained by crossing *soq1 koLHCII* and *koLHCII lcnp*. Seeds were surface sterilized using 70% ethanol and sown on MS plates (Murashige and Skoog Basal Salt Mixture, Duchefa Biochemie, with pH adjusted to 5.7 with KOH) and placed for 1 day in the dark at 4°C. Plates are then transferred into a growth cabinet for 2 to 3 weeks with 16 h light (Philips F17T8/TL741/ALTO 17W) at 150 µmol photons m^-2^ s^-1^ and 8 h dark at constant temperature 22°C. Seedlings were then transferred into soil (1:3 mixture of Agra-vermiculite + “yrkeskvalité K-JORD/krukjord” provided by RHP and Hasselfors garden respectively) and placed into a short-day growth room 8 h light with 150 µmol photons m^-2^ s^-1^ light (Philips F17T8/TL841/ALTO 17W) at 22°C and 16 h dark at 18°C. For cold and high light stress conditions, full plants (for thylakoid preparation) or detached leaves from the different individuals were placed on a moist paper on a plate first in a walk-in cold room at 4°C for 6 h in the dark then placed under an LED panel (custom-designed LED panel WH3000 built by JBeamBio with cool white LEDs BXRA-56C1100-B-00 (Farnell)) at 1,600 µmol photons m^-2^ s^-1^ for 8 h at 4°C. This cold and high-light treatment slightly differs from (Malnoë et al. 2017) to eliminate the need to visit the laboratory during the nighttime hours, where cold and high-light stresses were started simultaneously and were done for 6 hours; NPQ levels of the controls were similar to those previously published. For the photoprotection experiment, detached leaves were placed on a moist paper on a plate and exposed to cold (4°C) and high light (1,600 µmol photons m^-2^ s^-1^) until the leaves bleached (up to 45h). Chlorophyll content was monitored using the chlorophyll content meter CCM-200 from Opti-Sciences.

### Identification of cpsrp43 mutation by mapping by sequencing

To identify the mutation of interest, the mutant E40 (*soq1 npq4 gl1 cpsrp43-3*) was crossed to the *soq1 npq4 gl1* parental line, which was used to generate the EMS population (described in (Malnoë et al. 2017, Bru et al. 2020)). Plants displaying the mutant phenotype (low F_o_) in the F2 generation were isolated (Figure 1B) and pooled for DNA extraction (60 seedlings). Genomic DNA was also extracted from *soq1 npq4 gl1* (60 seedlings). The seedlings were ground with a mortar and pestle in liquid nitrogen, and then the powder was separated in four different tubes for DNA extractions. The four DNA extractions were prepared using a DNeasy Plant Mini kit (Qiagen), pooled, precipitated in ethanol plus sodium acetate, and eluted in 35 µL of water. DNA concentration was measured with Qubit and DNA integrity was assessed by running 50 ng of DNA on agarose gel. 600 ng of gDNA was sent to Novogene for sequencing library preparation and sequencing. The samples were run on Illumina NovaSeq 6000 to obtain 150-bp paired-end reads. The sequencing reads were analyzed using the online tool artMAP (Javorka et al. 2019). Briefly, reads from the control and mutant are quality filtered by trimgalore and aligned by BWA. Single nucleotide polymorphisms (SNPs) are identified by SAMtools and BCFtools, filtered, subtracted and annotated. The SNPs in the pooled E40 mutant F2 was then plotted relative to the genomic position (Figure S2A).

### Genetic crosses, genome editing and genotyping

Genetic crosses were done using standard techniques (Weigel and Glazebrook 2006). The mutant *soq1 lhcb5, soq1 koLHCII lcnp* have been generated by crossing *soq1-1* with *lhcb5* and *soq1 koLHCII* with *koLHCII lcnp* respectively. Genome editing assisted by CRISPR–Cas9 was used to edit *SOQ1* or *LCNP* or *LHCB6* (Supplemental Table 1). The mutant *soq1 koLHCII* was generated by targeting *SOQ1* using CRISPR-Cas9 in *koLHCII* (*SOQ1* sgRNA1, 2, 3 and 4). The mutant *koLHCII lcnp, soq1 lhcb1 lcnp* and *soq1 npq4 cpsrp43 lcnp* were generated from *koLHCII, soq1 lhcb1* and *soq1 npq4 cpsrp43* by targeting *LCNP* by CRISPR-Cas9 (*LCNP* sgRNA1, 2, 3 and 4). The mutant *soq1 lhcb6-2* was generated by targeting *LHCB6* by CRISPR-Cas9 in *soq1-1* (*LHCB6* sgRNA1, 2, 3 and 4). *SOQ1*, *LCNP* and *LHCB6* CRISPR-Cas9 was done using a construct carrying four single guide RNA (sgRNA) produced as described in (Ordon et al. 2017, Ordon et al. 2020). The *SOQ1* sgRNA was chosen according to (Concordet and Haeussler 2018), *LCNP* and *LHCB6* sgRNA were chosen according to (Labun et al. 2019). The mutants *soq1 NoM, soq1 lhcb4* and *soq1 lhcb6* (T-DNA) were generated by targeting *SOQ1* by CRISPR-Cas9 in *NoM, lhcb4* and *lhcb6* respectively, using the targeting vector CF588 provided by Christian Fankhauser, University of Lausanne, Switzerland based on (Wang et al. 2015) carrying two sgRNA chosen according to (Labun et al. 2019) (*SOQ1* sgRNA5 and 6).

Plants were transformed (generation T0) by floral dipping with Agrobacterium GV3101 pSoup containing the vector pDGE277 with the four sgRNA or CF588 containing two sgRNA. Seeds from transformed plants by the pDGE277 vector were plated and selected on Murashige and Skoog plates with 25 μg ml^−1^ hygromycin. Seeds from transformed plants by the CF588 vector were selected based on GFP seed coat fluorescence. The hygromycin-resistant plants and plants selected from seeds coat GFP signal were isolated (generation T1), and the absence of SOQ1, LCNP or Lhcb6 was confirmed by immunoblot using specific antibodies. Lines from independent transformation events (i.e. several T1) were screened for their phenotype, genotyped and one representative line was selected (in the case of *soq1 lhcb6* (Crispr-Cas9 on *SOQ1* in SALK_077953), three T1s were isolated: #3, #7 and #10 with *soq1 lhcb6* #10 initially selected until we realized it was an outlier, see Fig. S6C). T2 (or further generation) individuals from each lines were characterized.

### Chlorophyll fluorescence measurement

Detached leaves from the different individuals were placed on wet paper on a plate for 20 min in the dark to allow fast-relaxing NPQ to turn off. SpeedZen fluorescence imaging setup from JbeamBio (Johnson et al. 2009) was used to measure chlorophyll fluorescence. The minimum chlorophyll fluorescence (F_0_) was measured on the dark acclimated leaves. Following F_0_ measurement, a saturating flash pulse at 2,600 µmol photons m^-2^ s^-1^ was applied to measure the maximum chlorophyll fluorescence (F_m_) when all the reaction centers were closed. Then, NPQ was induced for 10 min at 1200 µmol photons m^-2^ s^-1^ followed by 10 min of dark to relax NPQ. Maximum fluorescence F_m’_ is measured at different time points after a saturating flash pulse during the light and dark phase. NPQ is calculated as (F_m_ - F_m_’) / F_m_’ (Brooks and Niyogi 2011). In stress conditions, the NPQ level is calculated as above with F_m_ (dark-adapted plants) and F_m_’ (after a cold and high-light treatment followed by a 5 min dark adaptation to relax fast NPQ components). Statistical analyses were performed using ANOVA Tukey’s multiple comparisons with a 95% confidence interval for statistical analysis with GraphPad Prism software.

### Protein extraction

For total protein extraction, leaves were collected in a 2 mL Eppendorf tube containing three glass beads and frozen in liquid nitrogen. Then the leaves were ground using beads mill for 30 seconds at 30 frequencies/s. A small volume of protein extraction buffer (50 mM HEPES, 150 mM NaCl, 5 mM MgCl_2_, 10% glycerol, 1% triton x100, PMSF and 2-mercaptoethanol) was then added into the tube. The extraction tube was then vortexed for 30 seconds and centrifuged to remove all solid pieces. Protein concentration was then measured with the Bradford method. Thylakoid extractions were performed according to (Iwai et al. 2015). Briefly, leaves from 6-week-old plants dark-adapted overnight or exposed to a cold and high-light treatment were harvested and ground in a blender for 30 s in 50 mL B1 cold solution (20 mM Tricine-KOH pH 7.8, 400 mM NaCl, 2 mM MgCl_2_). The solution was then filtrated through four layers of Miracloth and centrifuged 5 min at 27,000 x g at 4°C. The supernatant was discarded, and the pellet was resuspended in 15 mL B2 solution (20 mM Tricine-KOH pH 7.8, 150 mM NaCl, 5 mM MgCl_2_). B1 and B2 solution contain protease inhibitors (0.2 mM benzamidine, 1 mM aminocaproic acid, 0.2 mM PMSF). The resuspended solution was overlayed onto a 1.3 M/1.8 M sucrose cushion and ultracentrifuged for 30 min in a SW28 rotor at 19,000 rpm and 4°C. The band between the sucrose layers was collected and washed with B3 solution (20 mM Tricine-KOH pH 7.8, 15 mM NaCl, 5 mM MgCl_2_). The solution was centrifuged for 15 min at 27,000 x g and 4°C. The pellet was washed in storing solution (20 mM Tricine-KOH pH 7.8, 0.4 M sucrose, 15 mM NaCl, 5 mM MgCl_2_) and centrifuged for 10 min at 27,000 x g and 4°C. The pellet was then resuspended in storing solution. Chlorophyll concentration was measured according to (Porra et al. 1989).

### Immunoblot analysis

Thylakoids were washed and resuspended in 120 mM Tris HCl pH 6.8 followed by chlorophyll concentration measurement according to (Porra et al. 1989). Total proteins or thylakoids were then diluted in loading buffer (50 mM Tris pH 6.8, 2% SDS, 10% glycerol, 0.01% Bromophenol blue, 50 mM DTT) and boiled for 5 min. The samples were then loaded into an SDS-gel and ran until the migration front reached the bottom of the gel. The proteins were then transferred to a PVDF membrane activated in ethanol using a wet transfer system at constant 100V for 1h. The membrane was briefly rinsed with TBS-T and stained with Coomassie blue. The membrane was blocked with 10% milk for detection of LCNP or 3% milk for other proteins for 1h at room temperature. After a brief rinse with TBS-T, primary antibodies were added for 1h at room temperature or overnight at 4°C, cpSRP43_PHY2333S (1:1,000) from PhytoAB, Lhcb1_AS09 522 (1:5,000), Lhcb2_AS01 003 (1:10,000), Lhcb3_AS01 002 (1:2,000), Lhcb4_AS04 045 (1:7,000), Lhcb6_AS01 010 (1:5,000), ATP*b*_AS05 085 (1:5,000), PsbA D1_AS05 084 (1:10,000) from Agrisera, SOQ1 C-ter (1:200) (Malnoë et al. 2017), LCNP (1:200) (Bru et al. 2022), diluted in TBS-T + 3% milk. The membrane was washed 3 times 10 min with TBS-T, then incubated for 1h with the secondary goat anti-rabbit antibody conjugated to horse radish peroxidase AS09 602 (1:10,000) from Agrisera in TBS-T + 3% milk. The membrane was washed three times 10 min with TBS-T and one time 5 min with TBS. The Agrisera ECL Bright (AS16 ECL-N-100) or ECL SuperBright (AS16 ECL-S) and Azure biosystems c600 were used to reveal the bands.

### Antennae separation by clear native PAGE

Native PAGE was performed according to (Rantala et al. 2018) with modifications. Thylakoids were washed two times in the 25 mM Bis-Tris/HLC pH 7.0, 20% glycerol buffer then resuspended at 1 mg mL^-1^ chlorophyll concentration. An equal volume of 2% α-dodecyl maltoside (α-DM) was added to the samples and vortexed for 1 min then kept in ice and dark for 15 min and vortexed every 5 min for 5 s. Samples were centrifuged 20 min at 18,000 x g to remove all non-solubilized material. Amphipol (A8-35, Anatrace) were added to the samples to a final concentration of 1%. The samples were mixed by pipetting up and down multiple times then kept on ice and dark for 10 min before loading onto the gel (NativePAGE 4–16%, Bis–Tris, 1.0 mm, Mini Protein Gel, 10-well from Thermo Fisher Scientific; catalog number: BN1001BOX). The gels were run with anode buffer 50 mM Bis-Tris/HCL pH 7.0 and cathode buffer 50 mM tricine, 15 mM Bis-Tris, 0.02% α-DM for 2h at 80V. The gels were scanned, and fluorescence measurement was done by using SpeedZen fluorescence imaging setup from JBeamBio (Johnson et al. 2009) adapted from (Aro et al. 2004). Quantification of green band intensity was performed on the scanned pictures using the GelAnalyzer 19.1 software (Istvan Lazar Jr., PhD, Istvan Lazar Sr., PhD, CSc, www.gelanalyzer.com – last accessed on Feb 9 2025). Before the second dimension, the bands were soaked in Tris-HCl pH 6.8 containing 2% SDS for 30 min. Second dimension gel was prepared and ran as a regular SDS gel. The gel was covered with the fixing solution (30% ethanol + 10% acid acetic) and placed on a shaker at room temperature two times 30 min. The fixing solution was then replaced by Sypro Ruby and placed on a shaker at room temperature overnight. The gel was washed with the washing solution (10% ethanol + 10% acid acetic) for 30 min at room temperature on a shaker. Finally, the gel was rinsed briefly with water. Azure biosystems c600 was used to reveal the bands.

### Isolation of pigment–protein complexes by fast protein liquid chromatography (FPLC)

Thylakoid membranes (400 µg Chl) were solubilized at a concentration of 2 mg mL^-1^ with 4% (w/v) α-dodecyl maltoside (α-DM) for 15 min on ice, with brief mixing every 5 min. Unsolubilized membranes were then removed by centrifugation at 14,000 rpm for 5 min. Gel filtration chromatography was carried out according to (Iwai et al. 2015) using an ÄKTAmicro chromatography system and a Superdex 200 Increase 10/300 GL column (GE Healthcare), which was equilibrated with 20 mM Tris–HCl (pH 8.0), 5 mM MgCl_2_, and 0.03% (w/v) α-DM at 4 C. The flow rate was set to 1 mL min^-1^, and protein detection was done at 280 nm absorbance.

## Accession Numbers

Sequence data from this article can be found in the Arabidopsis Genome Initiative under accession numbers At1g56500 (SOQ1), At3g47860 (LCNP), At1g44575 (PsbS), At1g29920 (Lhcb1.1), At1g29910 (Lhcb1.2), At1g29930 (Lhcb1.3), At2g34430 (Lhcb1.4), At2g34420 (Lhcb1.5), At5g01530 (Lhcb4.1), At3g08940 (Lhcb4.2), At2g40100 (Lhcb4.3), At4g10340 (Lhcb5), At1g15820 (Lhcb6), At2g47450 (cpSRP43). Illumina NovaSeq 6000 sequencing data for *soq1 npq4 cpsrp43* (mutant E40) can be found in the Sequence Read Archive (https://www.ncbi.nlm.nih.gov/sra) under BioProject number PRJNA1009991 and BioSample numbers SAMN37176931.

## Supporting information

qH minor Supplemental Figure

qH minor Source Data

## Acknowledgements

We thank Stefano Caffarri, Jingfang Hao and Maria Paola Puggioni for critical discussions and Christian Fankhauser, University of Lausanne, Switzerland for providing the CF588 CRISPR-Cas9 vector. R.B. acknowledges the financial support from the European Research Council (ERC Advanced Grant 101053983-GrInSun). This research was supported by a Carl Kempe Postdoc Fellowship to A.C. (SMK-1855.2), European Commission Marie Skłodowska-Curie Actions Individual Fellowship Reintegration Panel to A.M. (845687), by a starting grant to A. M. from the Swedish Research Council Vetenskapsrådet (2018-04150), and by a consortium grant from the Swedish Foundation for Strategic Research (ARC19-0051). The research performed at Umeå University (P. B., A.C., A. M.) was further supported by grants to UPSC from the Knut and Alice Wallenberg Foundation (2016.0341, 2016.0352), and the Swedish Governmental Agency for Innovation Systems (2016-00504). The research performed at Indiana University Bloomington (P.B., A.M.) was supported by a start-up fund to A.M.

## Author Contributions

P.B., A.C., and A. M. designed the research; P.B., A.C., Y.P., Z.G. and A. M. performed research; Z.G., R.B., L.D. contributed new tools; P.B., A.C., and A. M. analyzed data; P.B., A.C., and A. M. wrote the paper with input from Z.G., L.D.

## References

1. Albanese, P., Manfredi, M., Marengo, E., Saracco, G. and Pagliano, C. (2019). Structural and functional differentiation of the light-harvesting protein Lhcb4 during land plant diversification. Physiol Plant 166(1): 336–350.

2. Albanese, P., Manfredi, M., Meneghesso, A., Marengo, E., Saracco, G., Barber, J., Morosinotto, T. and Pagliano, C. (2016). Dynamic reorganization of photosystem II supercomplexes in response to variations in light intensities. Biochim Biophys Acta 1857(10): 1651–1660.

3. Amin, P., Sy, D. A., Pilgrim, M. L., Parry, D. H., Nussaume, L. and Hoffman, N. E. (1999). Arabidopsis mutants lacking the 43- and 54-kilodalton subunits of the chloroplast signal recognition particle have distinct phenotypes. Plant Physiol 121(1): 61–70.

4. Amstutz, C. L., Fristedt, R., Schultink, A., Merchant, S. S., Niyogi, K. K. and Malnoë, A. (2020). An atypical short-chain dehydrogenase-reductase functions in the relaxation of photoprotective qH in Arabidopsis. Nat Plants 6(2): 154–166.

5. Anderson, S. A., Satyanarayan, M. B., Wessendorf, R. L., Lu, Y. and Fernandez, D. E. (2021). A homolog of GuidedEntry of Tail-anchored proteins3 functions in membrane-specific protein targeting in chloroplasts of Arabidopsis. The Plant Cell 33(8): 2812–2833.

6. Aro, E.-M., Suorsa, M., Rokka, A., Allahverdiyeva, Y., Paakkarinen, V., Saleem, A., Battchikova, N. and Rintamäki, E. (2004). Dynamics of photosystem II: a proteomic approach to thylakoid protein complexes. Journal of Experimental Botany 56(411): 347–356.

7. Bailey, S., Walters, R. G., Jansson, S. and Horton, P. (2001). Acclimation of Arabidopsis thaliana to the light environment: the existence of separate low light and high light responses. Planta 213(5): 794–801.

8. Ballottari, M., Dall’Osto, L., Morosinotto, T. and Bassi, R. (2007). Contrasting behavior of higher plant photosystem I and II antenna systems during acclimation. J Biol Chem 282(12): 8947–8958.

9. Ballottari, M., Girardon, J., Dall’osto, L. and Bassi, R. (2012). Evolution and functional properties of photosystem II light harvesting complexes in eukaryotes. Biochim Biophys Acta 1817(1): 143–157.

10. Bassi, R. and Dall’Osto, L. (2021). Dissipation of Light Energy Absorbed in Excess: The Molecular Mechanisms. Annual Review of Plant Biology 72(Volume 72, 2021): 47–76.

11. Betterle, N., Ballottari, M., Zorzan, S., de Bianchi, S., Cazzaniga, S., Dall’osto, L., Morosinotto, T. and Bassi, R. (2009). Light-induced dissociation of an antenna hetero-oligomer is needed for non-photochemical quenching induction. J Biol Chem 284(22): 15255–15266.

12. Brooks, M. D. and Niyogi, K. K. (2011). Use of a pulse-amplitude modulated chlorophyll fluorometer to study the efficiency of photosynthesis in Arabidopsis plants. Methods Mol Biol 775: 299–310.

13. Brooks, M. D., Sylak-Glassman, E. J., Fleming, G. R. and Niyogi, K. K. (2013). A thioredoxin-like/beta-propeller protein maintains the efficiency of light harvesting in *Arabidopsis*. Proc Natl Acad Sci USA 110(29): 2733–2740.

14. Bru, P., Nanda, S. and Malnoë, A. (2020). A Genetic Screen to Identify New Molecular Players Involved in Photoprotection qH in Arabidopsis thaliana. Plants (Basel) 9(11).

15. Bru, P., Steen, C. J., Park, S., Amstutz, C. L., Sylak-Glassman, E. J., Lam, L., Fekete, A., Mueller, M. J., Longoni, F., Fleming, G. R., Niyogi, K. K. and Malnoë, A. (2022). The major trimeric antenna complexes serve as a site for qH-energy dissipation in plants. J Biol Chem 298(11): 102519.

16. Bujaldon, S., Kodama, N., Rathod, M. K., Tourasse, N., Ozawa, S. I., Selles, J., Vallon, O., Takahashi, Y. and Wollman, F. A. (2020). The BF4 and p71 antenna mutants from Chlamydomonas reinhardtii. Biochim Biophys Acta Bioenerg 1861(4): 148085.

17. Caffarri, S., Croce, R., Cattivelli, L. and Bassi, R. (2004). A look within LHCII: differential analysis of the Lhcb1-3 complexes building the major trimeric antenna complex of higher-plant photosynthesis. Biochemistry 43(29): 9467–9476.

18. Caffarri, S., Kouril, R., Kereiche, S., Boekema, E. J. and Croce, R. (2009). Functional architecture of higher plant photosystem II supercomplexes. EMBO J 28(19): 3052–3063.

19. Caffarri, S., Tibiletti, T., Jennings, R. C. and Santabarbara, S. (2014). A comparison between plant photosystem I and photosystem II architecture and functioning. Curr Protein Pept Sci 15(4): 296–331.

20. Concordet, J. P. and Haeussler, M. (2018). CRISPOR: intuitive guide selection for CRISPR/Cas9 genome editing experiments and screens. Nucleic Acids Res 46(W1): W242–W245.

21. Crepin, A. and Caffarri, S. (2015). The specific localizations of phosphorylated Lhcb1 and Lhcb2 isoforms reveal the role of Lhcb2 in the formation of the PSI-LHCII supercomplex in Arabidopsis during state transitions. Biochim Biophys Acta 1847(12): 1539–1548.

22. Crepin, A. and Caffarri, S. (2018). Functions and Evolution of Lhcb Isoforms Composing LHCII, the Major Light Harvesting Complex of Photosystem II of Green Eukaryotic Organisms. Curr Protein Pept Sci 19(7): 699–713.

23. Cutolo, E. A., Caferri, R., Guardini, Z., Dall’Osto, L. and Bassi, R. (2023a). Analysis of state 1—state 2 transitions by genome editing and complementation reveals a quenching component independent from the formation of PSI-LHCI-LHCII supercomplex in Arabidopsis thaliana. Biology Direct 18(1): 49.

24. Cutolo, E. A., Guardini, Z., Dall’Osto, L. and Bassi, R. (2023b). A paler shade of green: engineering cellular chlorophyll content to enhance photosynthesis in crowded environments. New Phytol 239(5): 1567–1583.

25. Dall’Osto, L., Caffarri, S. and Bassi, R. (2005). A mechanism of nonphotochemical energy dissipation, independent from PsbS, revealed by a conformational change in the antenna protein CP26. Plant Cell 17(4): 1217–1232.

26. Dall’Osto, L., Cazzaniga, S., Bressan, M., Palecek, D., Zidek, K., Niyogi, K. K., Fleming, G. R., Zigmantas, D. and Bassi, R. (2017). Two mechanisms for dissipation of excess light in monomeric and trimeric light-harvesting complexes. Nat Plants 3: 17033.

27. Dall’Osto, L., Cazzaniga, S., Zappone, D. and Bassi, R. (2020). Monomeric light harvesting complexes enhance excitation energy transfer from LHCII to PSII and control their lateral spacing in thylakoids. Biochim Biophys Acta Bioenerg 1861(4): 148035.

28. Dall’Osto, L., Unlu, C., Cazzaniga, S. and van Amerongen, H. (2014). Disturbed excitation energy transfer in Arabidopsis thaliana mutants lacking minor antenna complexes of photosystem II. Biochim Biophys Acta 1837(12): 1981–1988.

29. Damkjaer, J. T., Kereiche, S., Johnson, M. P., Kovacs, L., Kiss, A. Z., Boekema, E. J., Ruban, A. V., Horton, P. and Jansson, S. (2009). The photosystem II light-harvesting protein Lhcb3 affects the macrostructure of photosystem II and the rate of state transitions in Arabidopsis. Plant Cell 21(10): 3245–3256.

30. de Bianchi, S., Betterle, N., Kouril, R., Cazzaniga, S., Boekema, E., Bassi, R. and Dall’Osto, L. (2011). Arabidopsis mutants deleted in the light-harvesting protein Lhcb4 have a disrupted photosystem II macrostructure and are defective in photoprotection. Plant Cell 23(7): 2659–2679.

31. de Bianchi, S., Dall’Osto, L., Tognon, G., Morosinotto, T. and Bassi, R. (2008). Minor antenna proteins CP24 and CP26 affect the interactions between photosystem II subunits and the electron transport rate in grana membranes of Arabidopsis. Plant Cell 20(4): 1012–1028.

32. De Souza, A. P., Burgess, S. J., Doran, L., Hansen, J., Manukyan, L., Maryn, N., Gotarkar, D., Leonelli, L., Niyogi, K. K. and Long, S. P. (2022). Soybean photosynthesis and crop yield are improved by accelerating recovery from photoprotection. Science 377(6608): 851–854.

33. Drop, B., Webber-Birungi, M., Yadav, S. K., Filipowicz-Szymanska, A., Fusetti, F., Boekema, E. J. and Croce, R. (2014). Light-harvesting complex II (LHCII) and its supramolecular organization in Chlamydomonas reinhardtii. Biochim Biophys Acta 1837(1): 63–72.

34. Dwyer, M. E. and Hangarter, R. P. (2022). Light-induced displacement of PLASTID MOVEMENT IMPAIRED1 precedes light-dependent chloroplast movements. Plant Physiol 189(3): 1866–1880.

35. Ferguson, J. N., Caproni, L., Walter, J., Shaw, K., Thein, M. S., Mager, S., Taylor, G., Cackett, L., Mathan, J., Vath, R. L., Martin, L., Genty, B., Pe, E., Lawson, T., Dell’Acqua, M. and Kromdijk, J. (2023). The genetic basis of dynamic non-photochemical quenching and photosystem II efficiency in fluctuating light reveals novel molecular targets for maize (Zea mays) improvement. bioRxiv: 2023.2011.2001.565118.

36. Flannery, S. E., Hepworth, C., Wood, W. H. J., Pastorelli, F., Hunter, C. N., Dickman, M. J., Jackson, P. J. and Johnson, M. P. (2021). Developmental acclimation of the thylakoid proteome to light intensity in Arabidopsis. Plant J 105(1): 223–244.

37. Floris, M., Bassi, R., Robaglia, C., Alboresi, A. and Lanet, E. (2013). Post-transcriptional control of light-harvesting genes expression under light stress. Plant Mol Biol 82(1-2): 147–154.

38. Foyer, C. H. and Hanke, G. (2022). ROS production and signalling in chloroplasts: cornerstones and evolving concepts. Plant J 111(3): 642–661.

39. Graca, A. T., Hall, M., Persson, K. and Schroder, W. P. (2021). High-resolution model of Arabidopsis Photosystem II reveals the structural consequences of digitonin-extraction. Sci Rep 11(1): 15534.

40. Guardini, Z., Gomez, R. L., Caferri, R., Dall’Osto, L. and Bassi, R. (2022a). Loss of a single chlorophyll in CP29 triggers re-organization of the Photosystem II supramolecular assembly. Biochimica et Biophysica Acta (BBA) - Bioenergetics 1863(5): 148555.

41. Guardini, Z., Gomez, R. L., Caferri, R., Stuttmann, J., Dall’Osto, L. and Bassi, R. (2022b). Thylakoid grana stacking revealed by multiplex genome editing of LHCII encoding genes. bioRxiv: 2021.2012.2031.474624.

42. Hao, J., Johansson, A., Svensson Fall, J., Duan, J., Hertle, A. P., Brooks, M. D., Niyogi, K. K., Yoshida, K., Hisabori, T. and Malnoë, A. (2024). Suppressor of quenching 1 functions as a methionine sulfoxide reductase in the chloroplast lumen for regulation of photoprotective qH in Arabidopsis. bioRxiv: 2024.2011.2001.621559.

43. Holzwarth, A. R., Miloslavina, Y., Nilkens, M. and Jahns, P. (2009). Identification of two quenching sites active in the regulation of photosynthetic light-harvesting studied by time-resolved fluorescence. Chemical Physics Letters 483(4): 262–267.

44. Horton, P., Ruban, A. V. and Walters, R. G. (1996). Regulation of Light Harvesting in Green Plants. Annu Rev Plant Physiol Plant Mol Biol 47: 655–684.

45. Hutin, C., Havaux, M., Carde, J. P., Kloppstech, K., Meiherhoff, K., Hoffman, N. and Nussaume, L. (2002). Double mutation cpSRP43--/cpSRP54-- is necessary to abolish the cpSRP pathway required for thylakoid targeting of the light-harvesting chlorophyll proteins. Plant J 29(5): 531–543.

46. Ilikova, I., Ilik, P., Opatikova, M., Arshad, R., Nosek, L., Karlicky, V., Kucerova, Z., Roudnicky, P., Pospisil, P., Lazar, D., Bartos, J. and Kouril, R. (2021). Towards spruce-type photosystem II: consequences of the loss of light-harvesting proteins LHCB3 and LHCB6 in Arabidopsis. Plant Physiol 187(4): 2691–2715.

47. Iwai, M., Patel-Tupper, D. and Niyogi, K. K. (2024). Structural Diversity in Eukaryotic Photosynthetic Light Harvesting. Annu Rev Plant Biol 75(1): 119–152.

48. Iwai, M., Yokono, M., Kono, M., Noguchi, K., Akimoto, S. and Nakano, A. (2015). Light-harvesting complex Lhcb9 confers a green alga-type photosystem I supercomplex to the moss Physcomitrella patens. Nat Plants 1: 14008.

49. Javorka, P., Raxwal, V. K., Najvarek, J. and Riha, K. (2019). artMAP: A user-friendly tool for mapping ethyl methanesulfonate-induced mutations in Arabidopsis. Plant Direct 3(6): e00146.

50. Jeong, J., Baek, K., Kirst, H., Melis, A. and Jin, E. (2017). Loss of CpSRP54 function leads to a truncated light-harvesting antenna size in Chlamydomonas reinhardtii. Biochim Biophys Acta Bioenerg 1858(1): 45–55.

51. Johnson, X., Vandystadt, G., Bujaldon, S., Wollman, F. A., Dubois, R., Roussel, P., Alric, J. and Beal, D. (2009). A new setup for in vivo fluorescence imaging of photosynthetic activity. Photosynth Res 102(1): 85–93.

52. Khaipho-Burch, M., Cooper, M., Crossa, J., de Leon, N., Holland, J., Lewis, R., McCouch, S., Murray, S. C., Rabbi, I., Ronald, P., Ross-Ibarra, J., Weigel, D. and Buckler, E. S. (2023). Genetic modification can improve crop yields - but stop overselling it. Nature 621(7979): 470–473.

53. Khorobrykh, S., Havurinne, V., Mattila, H. and Tyystjärvi, E. (2020). Oxygen and ROS in Photosynthesis. Plants 9(1): 91.

54. Kim, E.-H., Li, X.-P., Razeghifard, R., Anderson, J. M., Niyogi, K. K., Pogson, B. J. and Chow, W. S. (2009). The multiple roles of light-harvesting chlorophyll a/b-protein complexes define structure and optimize function of Arabidopsis chloroplasts: A study using two chlorophyll b-less mutants. Biochimica et Biophysica Acta (BBA) - Bioenergetics 1787(8): 973–984.

55. Kirst, H., Gabilly, S. T., Niyogi, K. K., Lemaux, P. G. and Melis, A. (2017). Photosynthetic antenna engineering to improve crop yields. Planta 245(5): 1009–1020.

56. Klimyuk, V. I., Persello-Cartieaux, F., Havaux, M., Contard-David, P., Schuenemann, D., Meiherhoff, K., Gouet, P., Jones, J. D., Hoffman, N. E. and Nussaume, L. (1999). A chromodomain protein encoded by the arabidopsis CAO gene is a plant-specific component of the chloroplast signal recognition particle pathway that is involved in LHCP targeting. Plant Cell 11(1): 87–99.

57. Kouril, R., Nosek, L., Bartos, J., Boekema, E. J. and Ilik, P. (2016). Evolutionary loss of light-harvesting proteins Lhcb6 and Lhcb3 in major land plant groups--break-up of current dogma. New Phytol 210(3): 808–814.

58. Kouril, R., Nosek, L., Opatikova, M., Arshad, R., Semchonok, D. A., Chamrad, I., Lenobel, R., Boekema, E. J. and Ilik, P. (2020). Unique organization of photosystem II supercomplexes and megacomplexes in Norway spruce. Plant J 104(1): 215–225.

59. Kouril, R., Wientjes, E., Bultema, J. B., Croce, R. and Boekema, E. J. (2013). High-light vs. low-light: effect of light acclimation on photosystem II composition and organization in Arabidopsis thaliana. Biochim Biophys Acta 1827(3): 411–419.

60. Kovacs, L., Damkjaer, J., Kereiche, S., Ilioaia, C., Ruban, A. V., Boekema, E. J., Jansson, S. and Horton, P. (2006). Lack of the light-harvesting complex CP24 affects the structure and function of the grana membranes of higher plant chloroplasts. Plant Cell 18(11): 3106–3120.

61. Kromdijk, J., Glowacka, K., Leonelli, L., Gabilly, S. T., Iwai, M., Niyogi, K. K. and Long, S. P. (2016). Improving photosynthesis and crop productivity by accelerating recovery from photoprotection. Science 354(6314): 857–861.

62. Labun, K., Montague, T. G., Krause, M., Torres Cleuren, Y. N., Tjeldnes, H. and Valen, E. (2019). CHOPCHOP v3: expanding the CRISPR web toolbox beyond genome editing. Nucleic Acids Res 47(W1): W171–W174.

63. Li, Z., Wakao, S., Fischer, B. B. and Niyogi, K. K. (2009). Sensing and Responding to Excess Light. Annual Review of Plant Biology 60(Volume 60, 2009): 239–260.

64. Long, S. P., Taylor, S. H., Burgess, S. J., Carmo-Silva, E., Lawson, T., De Souza, A. P., Leonelli, L. and Wang, Y. (2022). Into the Shadows and Back into Sunlight: Photosynthesis in Fluctuating Light. Annu Rev Plant Biol 73: 617–648.

65. Longoni, P., Douchi, D., Cariti, F., Fucile, G. and Goldschmidt-Clermont, M. (2015). Phosphorylation of the Light-Harvesting Complex II Isoform Lhcb2 Is Central to State Transitions. Plant Physiol 169(4): 2874–2883.

66. Malnoë, A. (2018). Photoinhibition or photoprotection of photosynthesis? Update on the (newly termed) sustained quenching component qH. Environmental and Experimental Botany 154: 123–133.

67. Malnoë, A., Schultink, A., Shahrasbi, S., Rumeau, D., Havaux, M. and Niyogi, K. K. (2017). The Plastid Lipocalin LCNP is Required for Sustained Photoprotective Energy Dissipation in Arabidopsis. Plant Cell 30: 196–208.

68. Müller, P., Li, X. P. and Niyogi, K. K. (2001). Non-photochemical quenching. A response to excess light energy. Plant Physiol 125(4): 1558–1566.

69. Opatikova, M., Semchonok, D. A., Kopecny, D., Ilik, P., Pospisil, P., Ilikova, I., Roudnicky, P., Zeljkovic, S. C., Tarkowski, P., Kyrilis, F. L., Hamdi, F., Kastritis, P. L. and Kouril, R. (2023). Cryo-EM structure of a plant photosystem II supercomplex with light-harvesting protein Lhcb8 and alpha-tocopherol. Nat Plants 9(8): 1359–1369.

70. Ordon, J., Bressan, M., Kretschmer, C., Dall’Osto, L., Marillonnet, S., Bassi, R. and Stuttmann, J. (2020). Optimized Cas9 expression systems for highly efficient Arabidopsis genome editing facilitate isolation of complex alleles in a single generation. Funct Integr Genomics 20(1): 151–162.

71. Ordon, J., Gantner, J., Kemna, J., Schwalgun, L., Reschke, M., Streubel, J., Boch, J. and Stuttmann, J. (2017). Generation of chromosomal deletions in dicotyledonous plants employing a user-friendly genome editing toolkit. Plant J 89(1): 155–168.

72. Pan, X., Ma, J., Su, X., Cao, P., Chang, W., Liu, Z., Zhang, X. and Li, M. (2018). Structure of the maize photosystem I supercomplex with light-harvesting complexes I and II. Science 360(6393): 1109–1113.

73. Pietrzykowska, M., Suorsa, M., Semchonok, D. A., Tikkanen, M., Boekema, E. J., Aro, E. M. and Jansson, S. (2014). The light-harvesting chlorophyll a/b binding proteins Lhcb1 and Lhcb2 play complementary roles during state transitions in Arabidopsis. Plant Cell 26(9): 3646–3660.

74. Porra, R. J., Thompson, W. A. and Kriedemann, P. E. (1989). Determination of accurate extinction coefficients and simultaneous equations for assaying chlorophylls a and b extracted with four different solvents: verification of the concentration of chlorophyll standards by atomic absorption spectroscopy. Biochimica et Biophysica Acta (BBA) - Bioenergetics 975(3): 384–394.

75. Rantala, M., Paakkarinen, V. and Aro, E. M. (2018). Analysis of Thylakoid Membrane Protein Complexes by Blue Native Gel Electrophoresis. J Vis Exp(139).

76. Sattari Vayghan, H., Nawrocki, W. J., Schiphorst, C., Tolleter, D., Hu, C., Douet, V., Glauser, G., Finazzi, G., Croce, R., Wientjes, E. and Longoni, F. (2022). Photosynthetic Light Harvesting and Thylakoid Organization in a CRISPR/Cas9 Arabidopsis Thaliana LHCB1 Knockout Mutant. Front Plant Sci 13: 833032.

77. Su, X., Ma, J., Wei, X., Cao, P., Zhu, D., Chang, W., Liu, Z., Zhang, X. and Li, M. (2017). Structure and assembly mechanism of plant C(2)S(2)M(2)-type PSII-LHCII supercomplex. Science 357(6353): 815–820.

78. Tokutsu, R., Kato, N., Bui, K. H., Ishikawa, T. and Minagawa, J. (2012). Revisiting the supramolecular organization of photosystem II in Chlamydomonas reinhardtii. J Biol Chem 287(37): 31574–31581.

79. Wang, Y., Burgess, S. J., de Becker, E. M. and Long, S. P. (2020). Photosynthesis in the fleeting shadows: an overlooked opportunity for increasing crop productivity? Plant J 101(4): 874–884.

80. Wang, Z. P., Xing, H. L., Dong, L., Zhang, H. Y., Han, C. Y., Wang, X. C. and Chen, Q. J. (2015). Egg cell-specific promoter-controlled CRISPR/Cas9 efficiently generates homozygous mutants for multiple target genes in Arabidopsis in a single generation. Genome Biol 16(1): 144.

81. Weigel, D. and Glazebrook, J. (2006). Setting up Arabidopsis crosses. CSH Protoc 2006(5).

82. Yu, G., Hao, J., Pan, X., Shi, L., Zhang, Y., Wang, J., Fan, H., Xiao, Y., Yang, F., Lou, J., Chang, W., Malnoë, A. and Li, M. (2022). Structure of Arabidopsis SOQ1 lumenal region unveils C-terminal domain essential for negative regulation of photoprotective qH. Nat Plants 8(7): 840–855.

83. Zhu, X. G., Ort, D. R., Whitmarsh, J. and Long, S. P. (2004). The slow reversibility of photosystem II thermal energy dissipation on transfer from high to low light may cause large losses in carbon gain by crop canopies: a theoretical analysis. J Exp Bot 55(400): 1167–1175.

84. Ziehe, D., Dunschede, B. and Schunemann, D. (2018). Molecular mechanism of SRP-dependent light-harvesting protein transport to the thylakoid membrane in plants. Photosynth Res 138(3): 303–313.

