## Supplementary material for "The minor antennae of photosystem II contribute to qH-energy dissipation in *Arabidopsis*": qH minor Supplemental Figure

**Affiliations**

**Short title:** Photoprotection by qH in the minor antennae

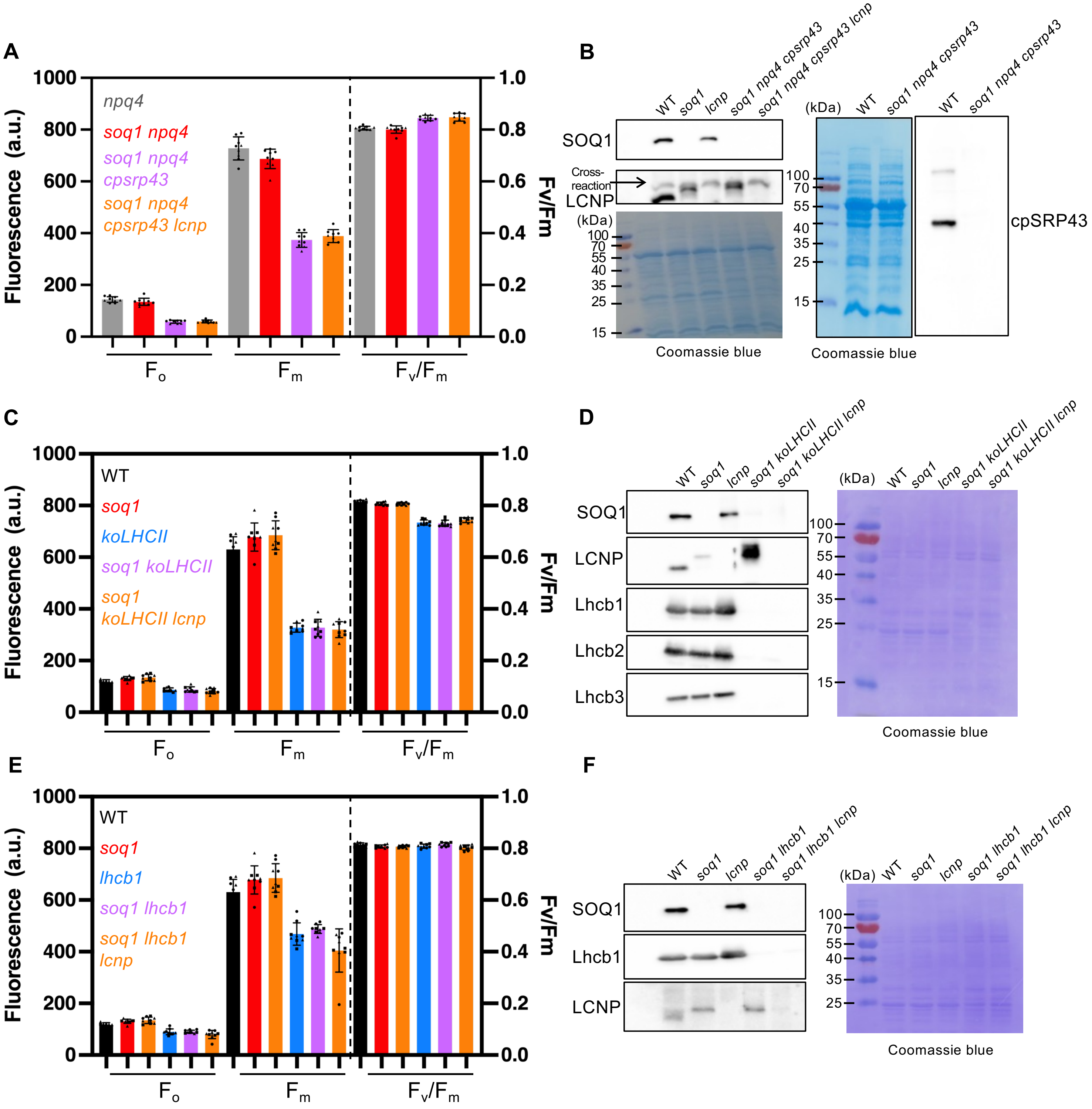


**Supplemental Figure 1. Fluorescence parameters and immunoblot analysis (supports Figure 1).** (A, C, E) Fluorescence parameters $F_0$, $F_m$ and $F_v/F_m$ of the same plant individuals measured in Figure 1 C, D (kinetics of *soq1 lhcb1* published in Bru et al. JBC 2022). Data represent means ± SD (n = 9, three detached leaves from independent individuals from three independent biological replicates, denoted by different symbols). (B, D, F) Immunoblot from 20 µg total proteins using antibodies against SOQ1, LCNP and cpSRP43 (B) and from 0.5 µg chlorophyll thylakoid using antibodies against SOQ1, LCNP, Lhcb1, Lhcb2 and Lhcb3 (D, F). Increased LCNP accumulation in *soq1 koLHCII* was not observed consistently.

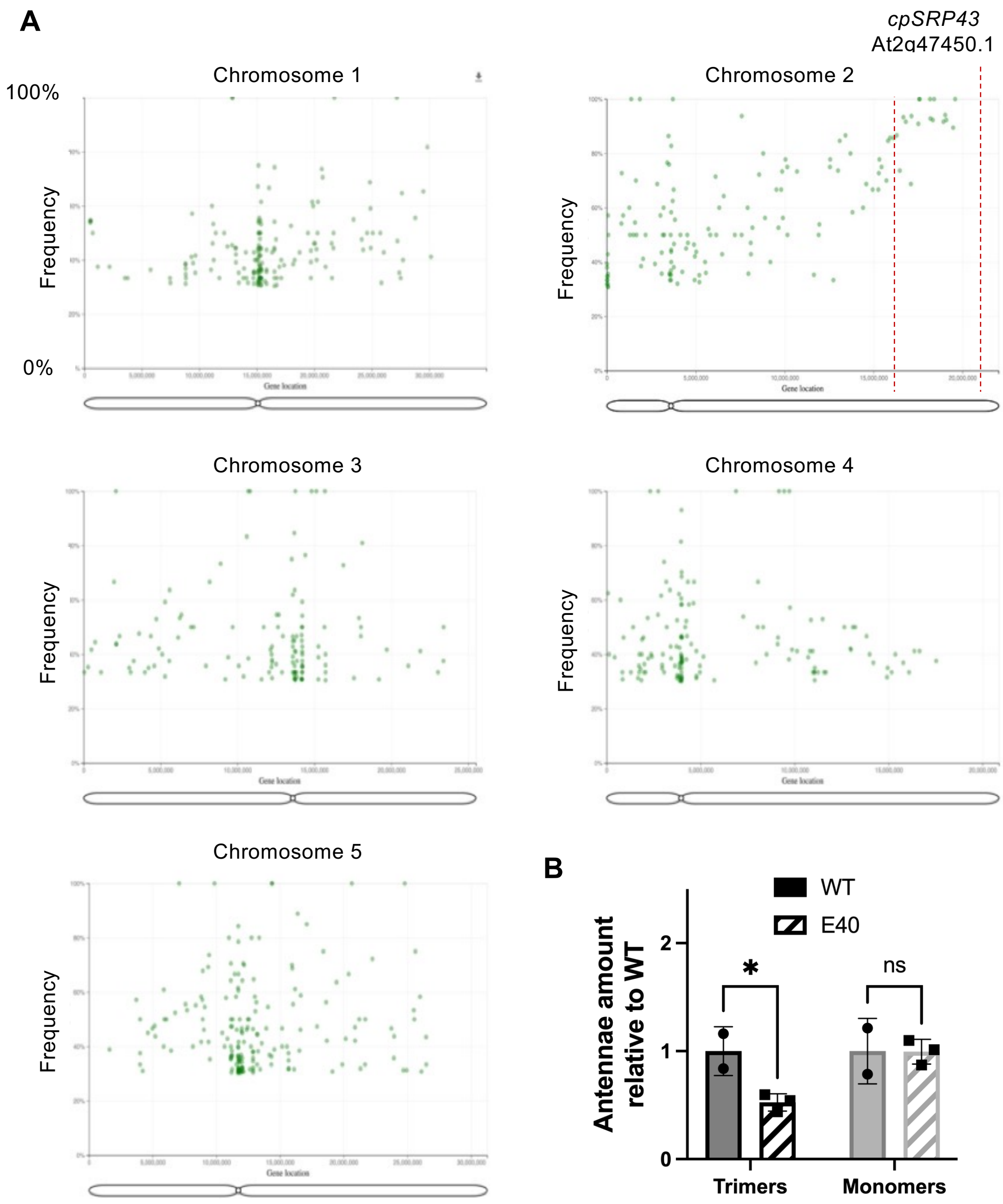


**Supplemental Figure 2. Mapping by sequencing identifies *cpsrp43* mutation in the E40 mutant (*soq1 npq4 cpsrp43*) (supports Figure 1).**

(A) Mapping-by-sequencing data analyzed by artMAP (Javorka et al. 2019). The figure represents the five Arabidopsis chromosomes with the detected single nucleotide polymorphisms (SNPs) of the pooled mutant F2 individuals from the cross *soq1 npq4 gl1 x E40* that had the same NPQ phenotype as E40. SNPs were filtered for quality and for the ones present in the parental line, the remaining SNPs were plotted with the allele frequency on the Y-axis and position on the X-axis for each chromosome. An increase in the allelic frequency of mutations approaching 100% was observed in the region between 15 and 20 Mb on chromosome 2, identifying this region as the one containing the causative mutation. (B) Bar graph representing PSII antennae quantification relative to WT as calculated from n=2 (WT) or n=3 (E40) separate CN-PAGE gels (see Figure 2).

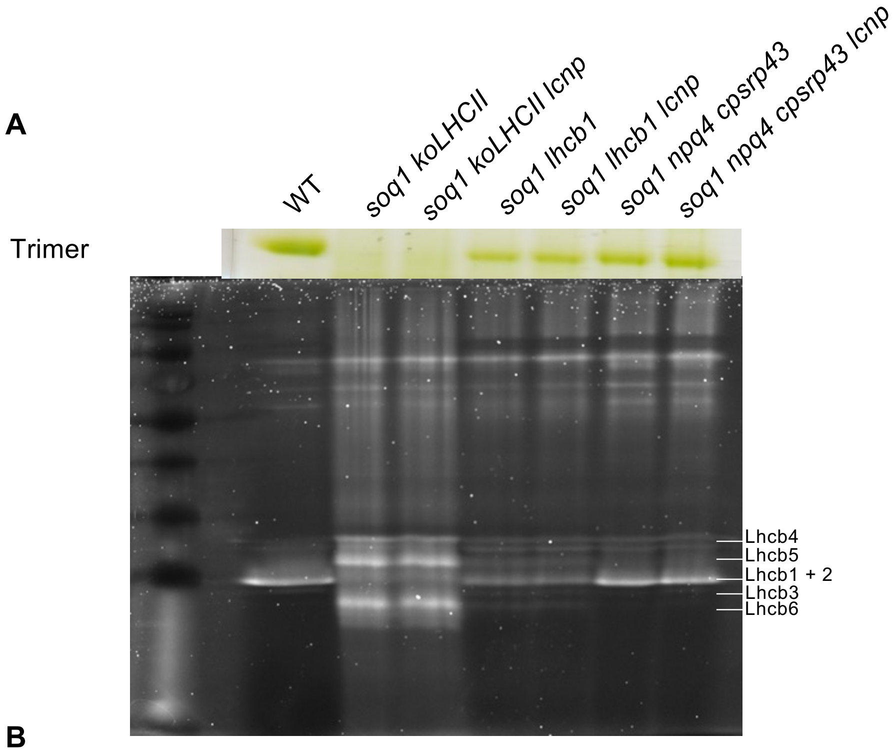


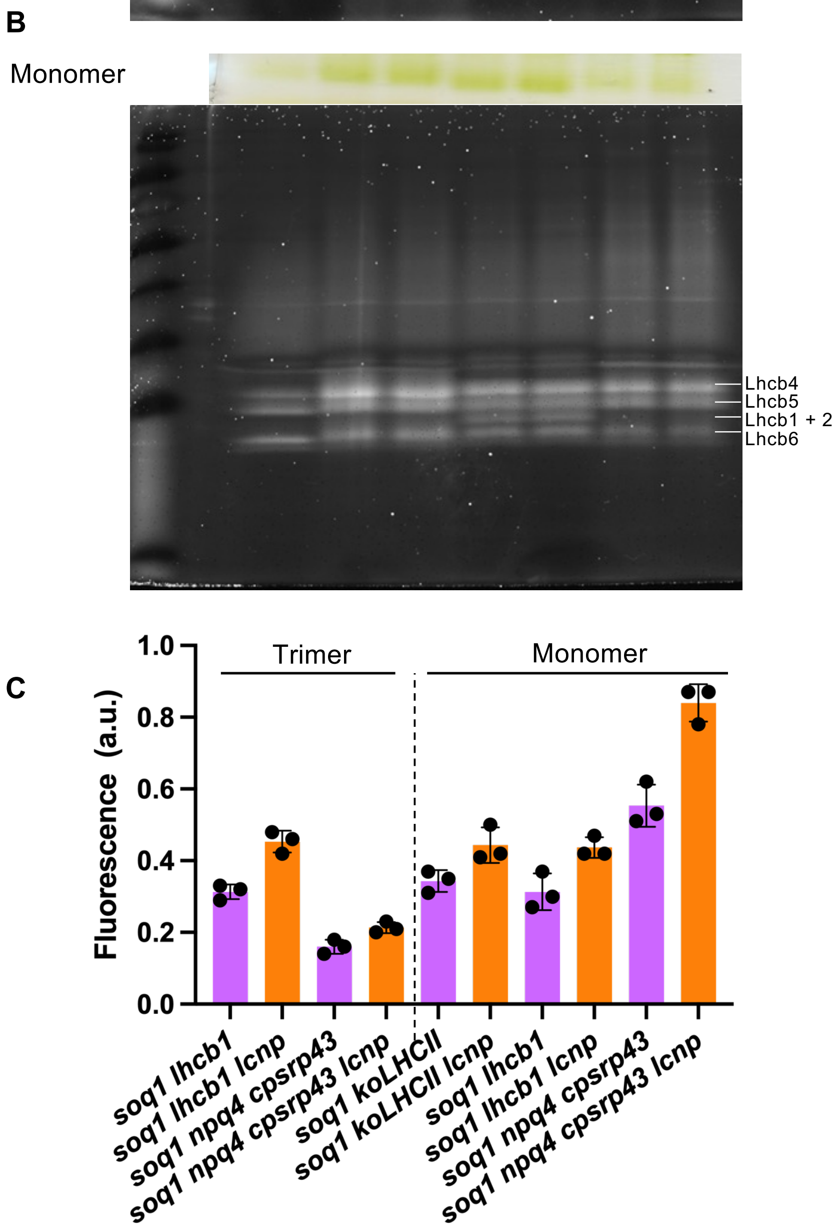


**Supplemental Figure 3. The bands corresponding to trimeric and monomeric antennae contain a similar composition of antennae subunits between active and inactive qH (supports Figure 2).**
SDS-PAGE followed by Sypro ruby staining from WT, *soq1 koLHCII, soq1 koLHCII lcnp, soq1 lhcb1, soq1 lhcb1 lcnp, soq1 npq4 cpsrp43* and *soq1 npq4 cpsrp43 lcnp* trimer (A) and monomer (B) bands excised from the CN-PAGE shown in Figure 2A. Representative experiment from three independent biological replicates. (C) Trimer and monomer band chlorophyll fluorescence quantification normalized by the intensity of the corresponding green band from the CN-PAGE. All the mutants were exposed to cold and high light stress (n=3 biological replicates).

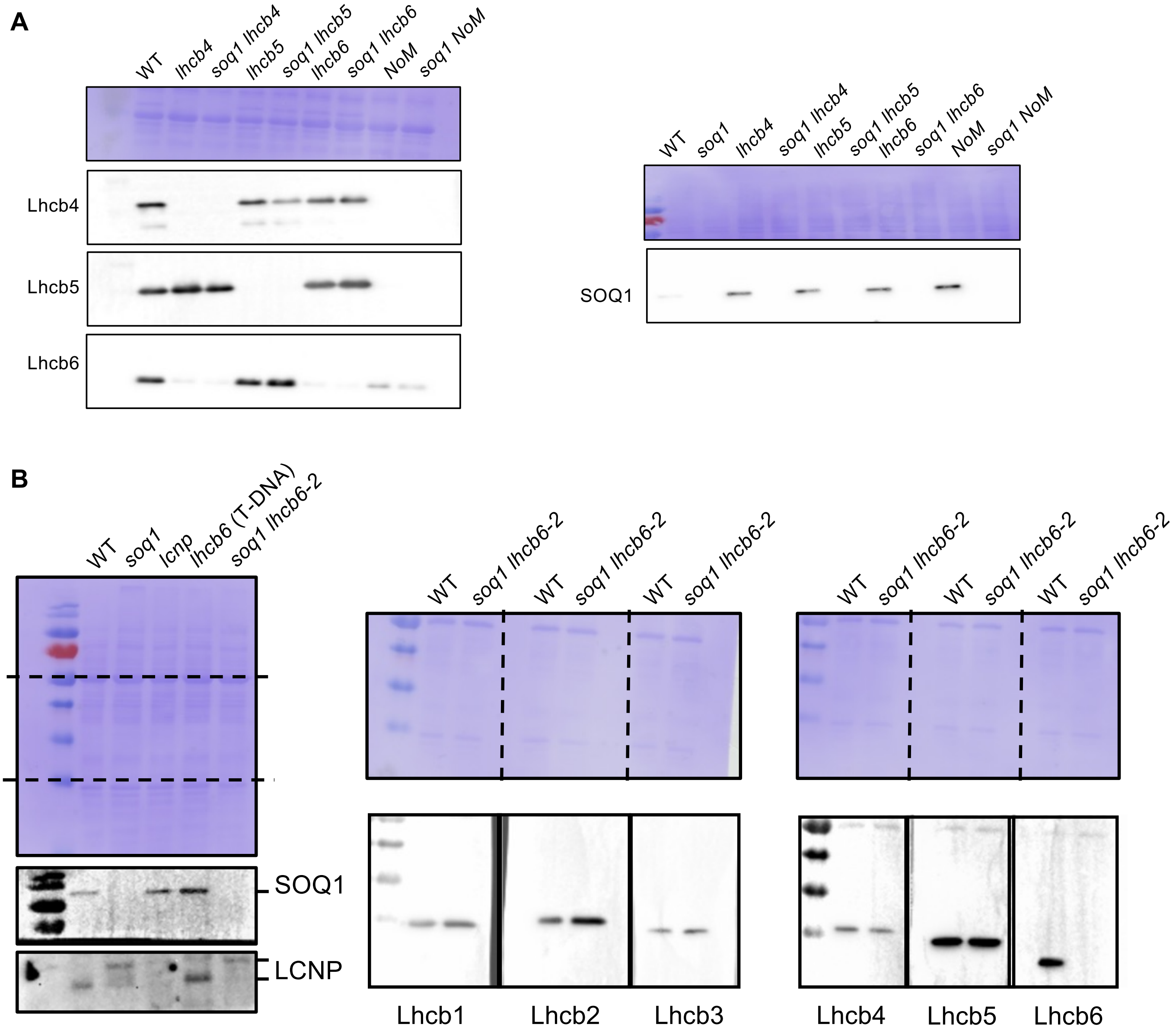


**Supplemental Figure 4. Mutants *soq1 NoM, soq1 lhcb4 and soq1 lhcb6* (T-DNA) accumulate a low amount of Lhcb6 (supports Figure 3).**
(A) Isolated thylakoid (0.5 µg chlorophyll). Immunoblot against Lhcb4, 5, 6 and SOQ1 in WT and the mutants *soq1, lhcb4, 5, 6* and *NoM* associated with the *soq1* mutation or not. (B) Total protein (1 µg). Immunoblot against SOQ1, LCNP in WT, *soq1, lcnp, lhcb6* (T-DNA) and *soq1 lhcb6-2* and Lhcb1, 2, 3, 4, 5 and 6 in WT and *soq1 lhcb6-2*.

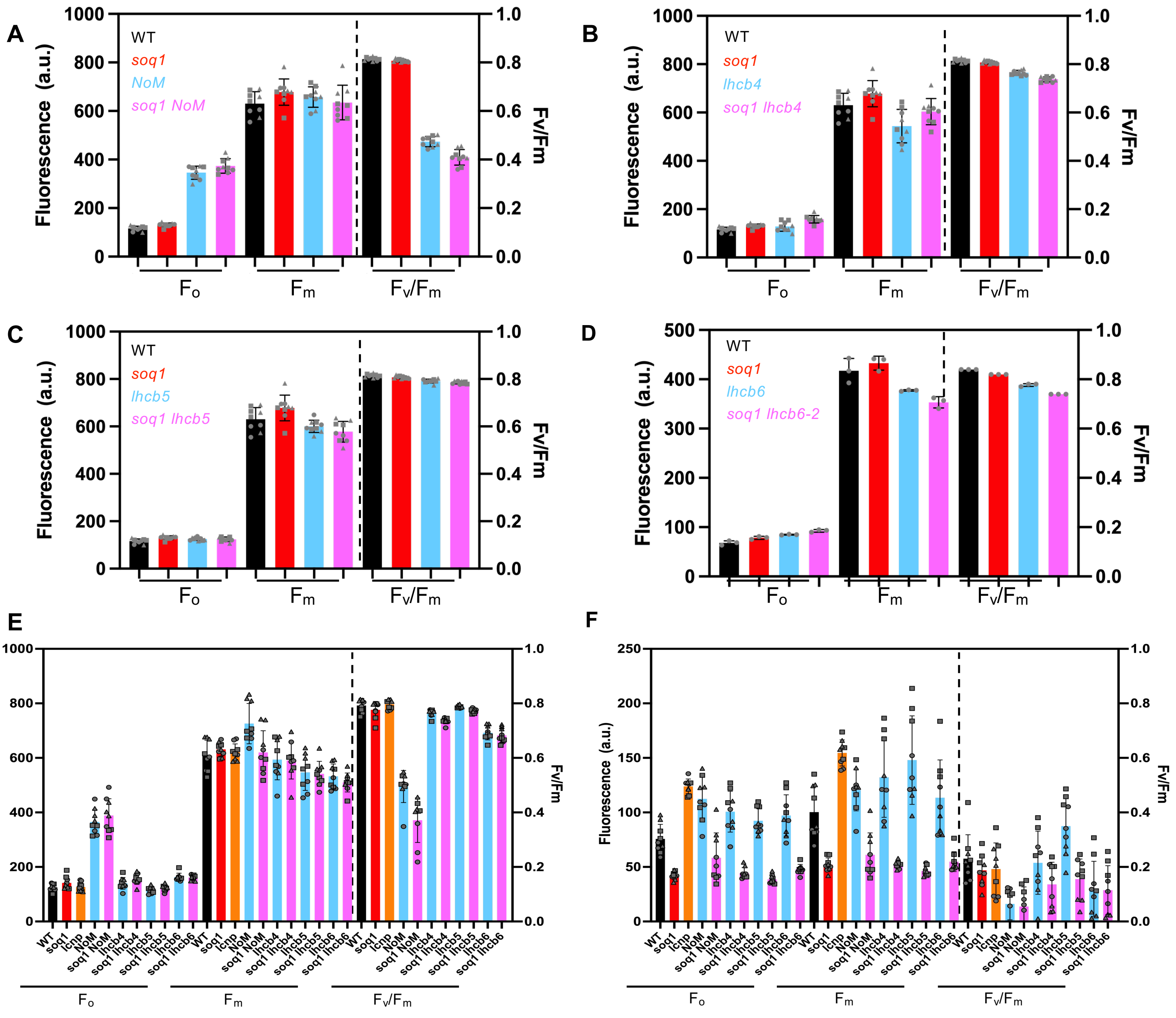


**Supplemental Figure 5. Fluorescence parameters (supports Figure 3).**
(A, B, C and D) fluorescence parameters $F_0$, $F_m$ and $F_v/F_m$ corresponding to the same plants in Figure 3A, B, C and D, respectively. WT (black), *soq1* (red), *lhcb* (blue) and *soq1 lhcb* (pink). All measurements were taken from 5 (E,F) or 20 (A,B,C,D) min dark-adapted detached leaves from 6-week-old plants. Data are presented as mean ± SD (n = 3 detached leaves from independent individuals). The results display a representative experiment, or all data, from three independent biological replicates, denoted by different symbols in (E,F). NPQ measurements were performed from plants grown at the same time for A, B and C. For D, the measurements were performed at a separate time. (E, F) Fluorescence parameter before (E) and after (F) cold and high light, $F_0$, $F_m$ and $F_v/F_m$ corresponding to the same plants shown in Figure 3E.

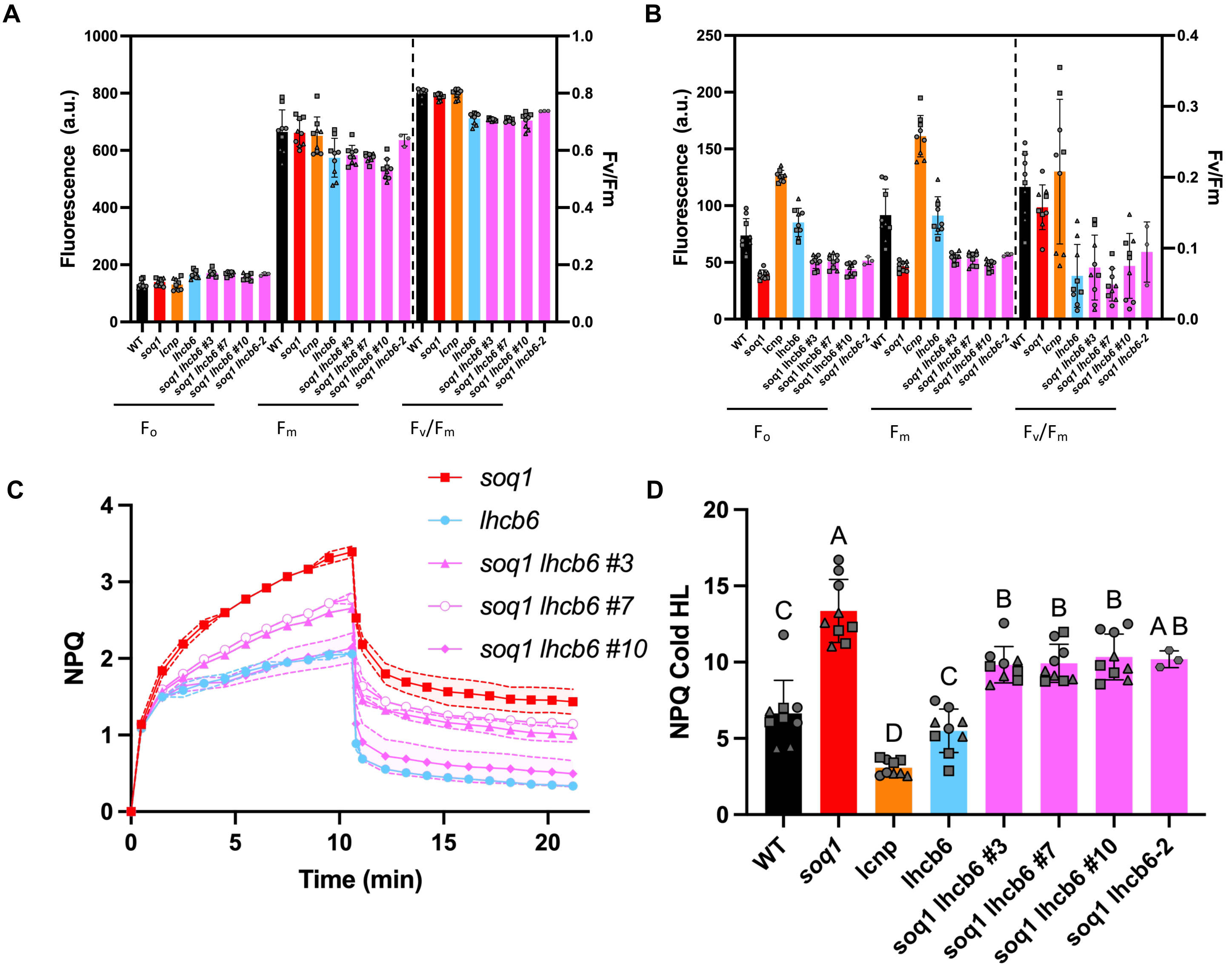


**Supplemental Figure 6. Independent *soq1 lhcb6* (T-DNA or CRISPR-Cas9) lines display a similar NPQ phenotype: slight decrease in qH compared to *soq1* after Cold HL (supports Figure 3).**
(A, B) Fluorescence parameter minimum ($F_0$), maximum ($F_m$) and $F_v/F_m$ fluorescence measurement before (A) and after (B) cold and high. (C) NPQ kinetics of *soq1* (red square), *lhcb6* (T-DNA) (blue circle) and *soq1 lhcb6 #3, #7, #10* (pink). (D) Bar graph representing NPQ after 6 h cold treatment followed by 8 h cold and high light (Cold HL). All measurements were taken from 5 (A,B,D) or 20 (C) min dark-adapted detached leaves from 6-week-old plants. Data are presented as mean ± SD (n = 3 detached leaves from independent individuals). The results display a representative experiment, or all data, from three independent biological replicates, denoted by different symbols in (A,B,D). Statistical analyses were made using ANOVA Tukey's multiple comparison test and a 95% confidence interval, revealing a significant increase in NPQ level of *soq1 lhcb6* compared to *lhcb6* and significant decrease in NPQ level of *soq1 lhcb6* compared to *soq1* ($p$ <0.05).

Of note, *soq1 lhcb6 #10* consistently showed a lower level of NPQ during short term kinetics experiment compared to #3, #7 (C) which may be due to an additional mutation elsewhere in the genome; we could not identify an obvious change by whole genome sequencing (not shown). However, after cold and HL treatment (D) #10 behaved as the two independent lines (#3, #7) and as *soq1 lhcb6-2*. We thus conclude that Lhcb6 indirectly, and only partly, affects qH.

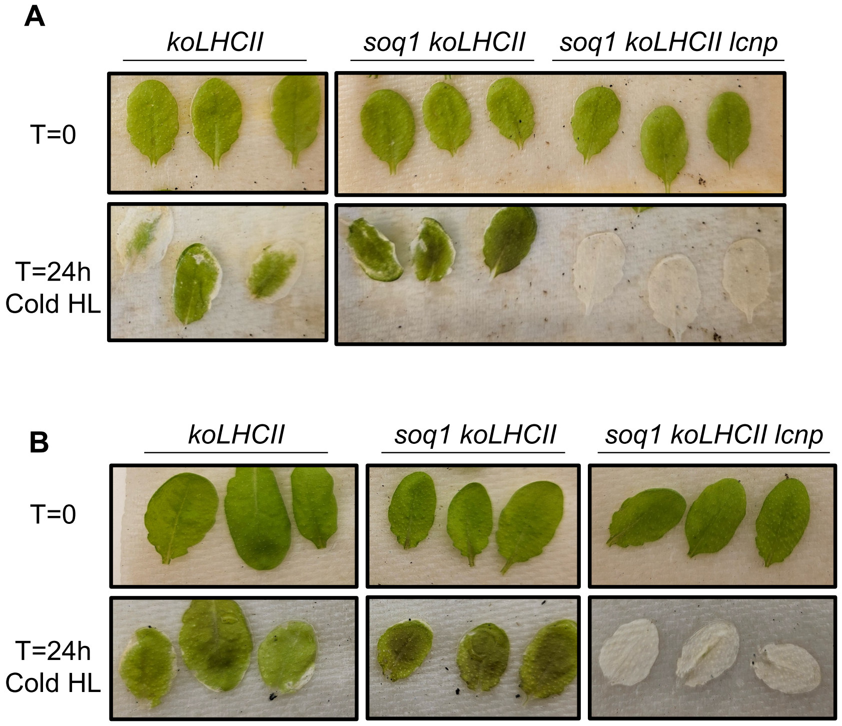


**Supplemental Figure 7. Replicates for the photobleaching experiment (supports Figure 5).**
(A,B) Images of detached leaves from *koLHCII*, *soq1 koLHCII*, *soq1 koLHCII lcnp* before (top) and after a 24h cold and high light (Cold HL) stress (bottom). All leaves from the different mutant lines were exposed together at the same time and were from plants grown at the same time. Three independent biological replicates, i.e. plants grown at a separate time, were performed.

**Supplemental Table 1. sgRNA used to produce mutation in *SOQ1*, *LCNP* or *LHCB6* by CRISPR-Cas9**

| | *SOQ1* | *LCNP* | *LHCB6* |
|---|---|---|---|
| sgRNA1 | CTCCGTAAAAACATCCACGG | CTTGTTGAAGTGGCAGCAGG | CTGGAGCTCAGAGAGTTGGC |
| sgRNA2 | GTCTACAAAACTGAGTGTGG | CTCACGTTACTGTCAGAAGA | ATAGTTCGCGAAATTCTCGG |
| sgRNA3 | CAATTGCGACGGATGATTGG | TGACATCATAAGGCAACTTG | CCGGCTCATAAACACCGTCG |
| sgRNA4 | GTCGTCTACAAAACTGAGTG | TCAGTCACTTCACAGTCCTG | GGTGTGGCATGGTTTGAAGC |
| sgRNA5 | GGATTGGGGGAAAGTATCGG | | |
| sgRNA6 | TGTTCTCTTCAAGCATTGCC | | |
