## Supplementary material for "The minor antennae of photosystem II contribute to qH-energy dissipation in *Arabidopsis*": qH minor Source Data

Fig1C

Rep 1

| Time (Min) | npq4 | | | soq1 npq4 | | | soq1 npq4 cpsrp43 | | | soq1 npq4 cpsrp43 lcnp | | |
|---|---|---|---|---|---|---|---|---|---|---|---|---|
| 0 | 0 | 0 | 0 | 0 | 0 | 0 | 0 | 0 | 0 | 0 | 0 | 0 |
| 0.50483 | 0.18089 | 0.15835 | 0.15059 | 0.23329 | 0.22147 | 0.22411 | 0.28619 | 0.25725 | 0.29182 | 0.17841 | 0.19867 | 0.18552 |
| 1.50967 | 0.35049 | 0.35115 | 0.34676 | 0.56992 | 0.53584 | 0.546 | 0.74149 | 0.67322 | 0.74025 | 0.45755 | 0.47709 | 0.41649 |
| 2.5145 | 0.52463 | 0.509 | 0.51583 | 0.88563 | 0.81131 | 0.83195 | 1.16541 | 1.05886 | 1.15332 | 0.69466 | 0.68493 | 0.65918 |
| 3.51934 | 0.65331 | 0.60289 | 0.64571 | 1.11528 | 1.06705 | 1.09038 | 1.55463 | 1.39878 | 1.46327 | 0.85837 | 0.81492 | 0.80669 |
| 4.52417 | 0.75715 | 0.68303 | 0.77684 | 1.31442 | 1.25837 | 1.28183 | 1.86936 | 1.71225 | 1.75747 | 0.95421 | 0.90365 | 0.88417 |
| 5.52901 | 0.82235 | 0.76716 | 0.87232 | 1.50391 | 1.42728 | 1.46768 | 2.15751 | 1.96009 | 2.02665 | 1.04379 | 0.97897 | 0.96167 |
| 6.53384 | 0.87557 | 0.8185 | 0.99409 | 1.65237 | 1.58021 | 1.61556 | 2.50594 | 2.191 | 2.23293 | 1.12009 | 1.01989 | 1.01837 |
| 7.53868 | 0.93558 | 0.8653 | 1.08424 | 1.77229 | 1.71936 | 1.76722 | 2.71781 | 2.39503 | 2.41659 | 1.19211 | 1.07154 | 1.06631 |
| 8.54351 | 0.97469 | 0.91341 | 1.17542 | 1.91166 | 1.8346 | 1.8942 | 2.94442 | 2.6227 | 2.59636 | 1.22604 | 1.12272 | 1.10114 |
| 9.54834 | 1.02559 | 0.96492 | 1.27576 | 2.0312 | 1.9591 | 2.03471 | 3.13235 | 2.78181 | 2.79293 | 1.27144 | 1.13915 | 1.17463 |
| 10.5532 | 1.06431 | 0.99096 | 1.35321 | 2.14137 | 2.05203 | 2.15559 | 3.36535 | 2.91221 | 2.92907 | 1.3058 | 1.18595 | 1.21538 |
| 10.8085 | 1.18195 | 1.12867 | 1.4404 | 2.40388 | 2.24569 | 2.36287 | 3.61177 | 3.11844 | 3.09311 | 1.47632 | 1.34144 | 1.37816 |
| 11.1463 | 1.15036 | 1.09258 | 1.46004 | 2.33366 | 2.21619 | 2.31359 | 3.39786 | 2.96434 | 2.89813 | 1.40738 | 1.24878 | 1.2972 |
| 12.1508 | 1.01689 | 1.02006 | 1.39648 | 2.24878 | 2.09491 | 2.21864 | 3.22014 | 2.81978 | 2.71577 | 1.17851 | 1.09309 | 1.05913 |
| 13.1553 | 0.90837 | 0.93353 | 1.3501 | 2.16421 | 1.97775 | 2.10295 | 2.92377 | 2.6912 | 2.58957 | 1.02117 | 0.94838 | 0.92492 |
| 14.1598 | 0.86765 | 0.90759 | 1.33018 | 2.10464 | 1.90902 | 2.07267 | 2.94743 | 2.62838 | 2.47483 | 0.92384 | 0.87404 | 0.82318 |
| 15.1643 | 0.83151 | 0.86809 | 1.31301 | 2.04359 | 1.84516 | 2.00741 | 2.79692 | 2.48662 | 2.39555 | 0.88118 | 0.82497 | 0.75695 |
| 16.1688 | 0.79518 | 0.85304 | 1.27859 | 1.99838 | 1.79675 | 1.96858 | 2.70777 | 2.3963 | 2.35949 | 0.84542 | 0.7698 | 0.74521 |
| 17.1733 | 0.77794 | 0.82868 | 1.26366 | 1.96854 | 1.74343 | 1.93904 | 2.73335 | 2.38178 | 2.38033 | 0.82342 | 0.7342 | 0.73066 |
| 18.1778 | 0.74134 | 0.76768 | 1.2544 | 1.88469 | 1.71162 | 1.90076 | 2.7216 | 2.35728 | 2.27729 | 0.82671 | 0.7205 | 0.71453 |
| 19.1823 | 0.72002 | 0.73662 | 1.24189 | 1.78045 | 1.65436 | 1.83194 | 2.58013 | 2.33909 | 2.21087 | 0.79027 | 0.69927 | 0.69787 |
| 20.1868 | 0.71372 | 0.71991 | 1.2238 | 1.77206 | 1.65697 | 1.81025 | 2.58609 | 2.33095 | 2.20416 | 0.80388 | 0.70281 | 0.69328 |
| 21.1913 | 0.70065 | 0.72208 | 1.24351 | 1.7478 | 1.65287 | 1.76931 | 2.61994 | 2.36139 | 2.28421 | 0.77903 | 0.67933 | 0.68096 |

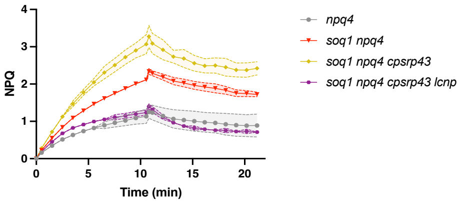


Rep2

| Time (Min) | npq4 | | | soq1 npq4 | | | soq1 npq4 cpsrp43 | | | soq1 npq4 cpsrp43 lcnp | | |
|---|---|---|---|---|---|---|---|---|---|---|---|---|
| 0 | 0 | 0 | 0 | 0 | 0 | 0 | 0 | 0 | 0 | 0 | 0 | 0 |
| 0.50483 | 0.14316 | 0.12547 | 0.15685 | 0.20966 | 0.26055 | 0.20227 | 0.2412 | 0.16973 | 0.19977 | 0.13072 | 0.1116 | 0.11378 |
| 1.50967 | 0.32641 | 0.30069 | 0.35875 | 0.53792 | 0.63296 | 0.52699 | 0.70436 | 0.57582 | 0.57077 | 0.34098 | 0.32341 | 0.32034 |
| 2.5145 | 0.49072 | 0.43923 | 0.51755 | 0.83107 | 0.93397 | 0.80557 | 1.1352 | 0.92884 | 0.90201 | 0.50685 | 0.48613 | 0.49335 |
| 3.51934 | 0.60569 | 0.55395 | 0.64036 | 1.07212 | 1.13883 | 1.03112 | 1.57306 | 1.2447 | 1.20442 | 0.62545 | 0.60479 | 0.61875 |
| 4.52417 | 0.70379 | 0.63123 | 0.71809 | 1.29641 | 1.34197 | 1.22292 | 1.83881 | 1.51636 | 1.48322 | 0.73224 | 0.72062 | 0.71539 |
| 5.52901 | 0.77287 | 0.69218 | 0.78327 | 1.48468 | 1.49859 | 1.36514 | 2.13757 | 1.74501 | 1.69659 | 0.82577 | 0.78761 | 0.80172 |
| 6.53384 | 0.82696 | 0.74381 | 0.84527 | 1.63199 | 1.6635 | 1.51175 | 2.52199 | 1.99809 | 1.95414 | 0.89229 | 0.878 | 0.85372 |
| 7.53868 | 0.88603 | 0.78557 | 0.91684 | 1.79146 | 1.78792 | 1.64927 | 2.71964 | 2.162 | 2.17764 | 0.98602 | 0.92536 | 0.90601 |
| 8.54351 | 0.94065 | 0.83134 | 0.97133 | 1.95397 | 1.92332 | 1.78233 | 3.05401 | 2.40428 | 2.3668 | 1.01196 | 0.99307 | 0.96077 |
| 9.54834 | 0.99326 | 0.8717 | 1.01734 | 2.13125 | 2.01681 | 1.90914 | 3.29814 | 2.62104 | 2.5819 | 1.09526 | 1.06462 | 1.03473 |
| 10.5532 | 1.04079 | 0.89182 | 1.06135 | 2.19749 | 2.10675 | 1.99532 | 3.42165 | 2.75424 | 2.77093 | 1.11286 | 1.08397 | 1.06485 |
| 10.8085 | 1.12923 | 0.99236 | 1.20796 | 2.44532 | 2.26629 | 2.19984 | 3.84399 | 3.06998 | 2.99598 | 1.28279 | 1.27916 | 1.30053 |
| 11.1463 | 1.14098 | 0.98208 | 1.19921 | 2.41463 | 2.25843 | 2.1919 | 3.96967 | 3.01365 | 2.94436 | 1.28276 | 1.2584 | 1.28017 |
| 12.1508 | 1.07018 | 0.90557 | 1.10366 | 2.36523 | 2.12712 | 2.06338 | 3.80731 | 2.94332 | 3.02397 | 1.16149 | 1.12097 | 1.14337 |
| 13.1553 | 0.99533 | 0.85036 | 1.02873 | 2.24123 | 2.04324 | 1.96813 | 3.55267 | 2.75753 | 2.77407 | 1.04234 | 1.00842 | 1.01359 |
| 14.1598 | 0.95331 | 0.81194 | 0.99262 | 2.14122 | 1.97823 | 1.89891 | 3.40674 | 2.69466 | 2.67885 | 1.00855 | 0.96329 | 0.9215 |
| 15.1643 | 0.94181 | 0.78666 | 0.96285 | 2.08924 | 1.97223 | 1.89058 | 3.32468 | 2.65801 | 2.72337 | 0.98732 | 0.93467 | 0.83317 |
| 16.1688 | 0.91816 | 0.74676 | 0.93672 | 2.00638 | 1.92966 | 1.81267 | 3.30443 | 2.5509 | 2.68224 | 0.9231 | 0.91079 | 0.79446 |
| 17.1733 | 0.89644 | 0.72328 | 0.9203 | 1.97935 | 1.9148 | 1.79331 | 3.3572 | 2.66377 | 2.69188 | 0.93626 | 0.92411 | 0.78042 |
| 18.1778 | 0.88865 | 0.69732 | 0.86087 | 1.91243 | 1.87806 | 1.72342 | 3.1202 | 2.5823 | 2.66975 | 0.92147 | 0.87637 | 0.77736 |
| 19.1823 | 0.8582 | 0.69247 | 0.82268 | 1.90215 | 1.81801 | 1.68242 | 3.11399 | 2.57515 | 2.71875 | 0.90909 | 0.86103 | 0.7569 |
| 20.1868 | 0.848 | 0.68668 | 0.82342 | 1.88704 | 1.78065 | 1.67721 | 3.12241 | 2.57649 | 2.67161 | 0.90162 | 0.86322 | 0.73202 |
| 21.1913 | 0.83509 | 0.68339 | 0.81891 | 1.85896 | 1.75725 | 1.64845 | 3.09161 | 2.55249 | 2.62878 | 0.89542 | 0.87088 | 0.73605 |

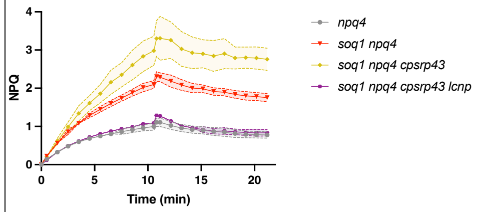


Rep 3

| Time (Min) | npq4 | | | soq1 npq4 | | | soq1 npq4 cpsrp43 | | | soq1 npq4 cpsrp43 lcnp | | |
|---|---|---|---|---|---|---|---|---|---|---|---|---|
| 0 | 0 | 0 | 0 | 0 | 0 | 0 | 0 | 0 | 0 | 0 | 0 | 0 |
| 0.50483 | 0.138744 | 0.16865 | 0.167123 | 0.208139 | 0.196149 | 0.198427 | 0.22434 | 0.260123 | 0.208735 | 0.158412 | 0.153968 | 0.154868 |
| 1.50967 | 0.317118 | 0.355494 | 0.35008 | 0.560114 | 0.533868 | 0.544662 | 0.625909 | 0.724105 | 0.605393 | 0.431816 | 0.367712 | 0.388025 |
| 2.5145 | 0.469673 | 0.480611 | 0.468484 | 0.850956 | 0.814044 | 0.828122 | 0.962361 | 1.113714 | 0.967718 | 0.619969 | 0.511876 | 0.566946 |
| 3.51934 | 0.598536 | 0.558143 | 0.541364 | 1.08509 | 1.043286 | 1.048231 | 1.211555 | 1.439213 | 1.26087 | 0.746314 | 0.61235 | 0.683451 |
| 4.52417 | 0.679152 | 0.614585 | 0.597292 | 1.289835 | 1.229947 | 1.228081 | 1.430164 | 1.751757 | 1.514657 | 0.847817 | 0.696356 | 0.768294 |
| 5.52901 | 0.737488 | 0.658665 | 0.653875 | 1.447121 | 1.393483 | 1.400723 | 1.627383 | 2.087297 | 1.708416 | 0.926236 | 0.758986 | 0.846075 |
| 6.53384 | 0.800801 | 0.709816 | 0.690631 | 1.604572 | 1.549201 | 1.550279 | 1.793445 | 2.414587 | 1.928906 | 0.990221 | 0.817206 | 0.908749 |
| 7.53868 | 0.858761 | 0.748773 | 0.739465 | 1.76465 | 1.692546 | 1.704685 | 1.959129 | 2.704116 | 2.093522 | 1.062075 | 0.870578 | 0.958918 |
| 8.54351 | 0.90718 | 0.787399 | 0.774766 | 1.926157 | 1.828933 | 1.834694 | 2.082959 | 3.011787 | 2.265024 | 1.114819 | 0.908108 | 1.005528 |
| 9.54834 | 0.955123 | 0.82333 | 0.819657 | 2.058312 | 1.970981 | 1.943984 | 2.211861 | 3.29115 | 2.407068 | 1.161482 | 0.940496 | 1.038639 |
| 10.5532 | 0.996596 | 0.855182 | 0.847017 | 2.169374 | 2.090202 | 2.072245 | 2.318191 | 3.472102 | 2.554224 | 1.208504 | 0.99677 | 1.074113 |
| 10.8085 | 1.12787 | 0.925236 | 0.952372 | 2.358766 | 2.280486 | 2.247319 | 2.442108 | 3.607566 | 2.721836 | 1.298266 | 1.019001 | 1.195207 |
| 11.1463 | 1.083869 | 0.880355 | 0.905909 | 2.321922 | 2.281649 | 2.233109 | 2.369239 | 3.488833 | 2.685591 | 1.214435 | 0.952839 | 1.119195 |
| 12.1508 | 0.991061 | 0.786221 | 0.786056 | 2.230922 | 2.178588 | 2.149114 | 2.209523 | 3.196046 | 2.524762 | 1.04436 | 0.84211 | 0.974369 |
| 13.1553 | 0.941295 | 0.722315 | 0.721803 | 2.150357 | 2.135783 | 2.049598 | 2.121309 | 3.012239 | 2.442277 | 0.950427 | 0.792549 | 0.88759 |
| 14.1598 | 0.890767 | 0.702891 | 0.698181 | 2.074296 | 2.053406 | 1.994703 | 2.063913 | 2.949174 | 2.317424 | 0.912864 | 0.766156 | 0.831575 |
| 15.1643 | 0.86771 | 0.705371 | 0.677546 | 2.058438 | 2.02182 | 1.943233 | 2.024214 | 2.87044 | 2.251844 | 0.877803 | 0.742043 | 0.803456 |
| 16.1688 | 0.838835 | 0.687654 | 0.677836 | 2.009327 | 2.021476 | 1.913142 | 2.02492 | 2.873155 | 2.246619 | 0.858378 | 0.735623 | 0.792567 |
| 17.1733 | 0.815311 | 0.673208 | 0.65705 | 2.002267 | 1.99258 | 1.906926 | 1.984287 | 2.816531 | 2.164466 | 0.816588 | 0.734242 | 0.758548 |
| 18.1778 | 0.787504 | 0.672269 | 0.657151 | 1.960277 | 1.978326 | 1.905102 | 1.971886 | 2.807564 | 2.131306 | 0.807184 | 0.724018 | 0.738921 |
| 19.1823 | 0.761319 | 0.659161 | 0.658743 | 1.96226 | 1.95812 | 1.908562 | 1.965713 | 2.743644 | 2.077535 | 0.778068 | 0.714502 | 0.711416 |
| 20.1868 | 0.746293 | 0.654685 | 0.646108 | 1.931005 | 1.959594 | 1.891764 | 1.960155 | 2.735633 | 2.04876 | 0.775296 | 0.720493 | 0.689545 |
| 21.1913 | 0.728449 | 0.652955 | 0.642511 | 1.937711 | 1.93291 | 1.865305 | 1.945703 | 2.70749 | 2.015782 | 0.752236 | 0.707349 | 0.654904 |

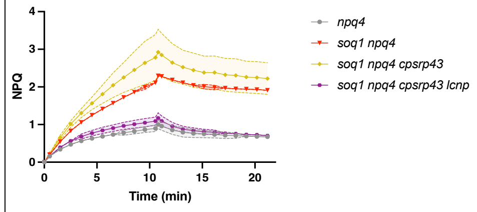


Fig1E and F

| | WT | soq1 | lcnp | lhcb1 | soq1 lhcb1 | q1 lhcb1 lcr | koLHCII | oq1 koLHC | 1 koLHCII lc | npq4 | soq1 npq4 | 1 npq4 cpsr | npq4 cpsrp43 lcnp |
|---|---|---|---|---|---|---|---|---|---|---|---|---|---|
| Rep 1 | 6.984126 | 11.22049 | 3.568467 | 6.155417 | 11.57901 | 5.331599 | 2.060759 | 7.472602 | 2.069366 | 11.76619 | 12.21218 | 13.64181 | 4.90516 |
| | 6.079087 | 12.3017 | 3.761288 | 6.908752 | 10.98044 | 2.502752 | 4.949891 | 7.688689 | 2.050938 | 8.879419 | 11.42191 | 11.92787 | 4.233995 |
| | 6.348455 | 12.07763 | 3.408579 | 6.364854 | 10.04532 | 3.539627 | 6.281439 | 7.0174 | 2.713579 | 8.193559 | 11.1595 | 15.68359 | 8.357904 |
| Rep 2 | 3.079066 | 9.428509 | 2.592094 | 4.054051 | 9.353052 | 3.857776 | 3.420625 | 7.590902 | 2.144753 | 3.341008 | 11.34938 | 8.781546 | 4.286061 |
| | 4.438568 | 10.97467 | 2.858726 | 3.318474 | 10.43788 | 2.111063 | 3.963524 | 5.737133 | 2.304409 | 9.802564 | 11.95144 | 13.07452 | 4.204707 |
| | 5.499522 | 8.716724 | 3.237524 | 2.71324 | 10.94726 | 3.002853 | 5.127475 | 6.536606 | 2.786768 | 10.33865 | 12.67154 | 12.82677 | 5.993534 |
| Rep 3 | 6.770212 | 12.59477 | 2.714203 | 3.566897 | 10.54139 | 4.221265 | 5.105338 | 6.538679 | 2.001513 | 10.73226 | 9.646322 | 16.56797 | 7.102891 |
| | 4.440138 | 11.03376 | 2.535785 | 6.593781 | 9.620975 | 4.935101 | 5.43095 | 7.196518 | 1.872761 | 8.675843 | 10.1558 | 14.49133 | 3.961067 |
| | 4.337826 | 13.24643 | 2.727231 | 6.113859 | 10.26253 | 6.998991 | 5.266141 | 6.897749 | 1.85177 | 6.875721 | 10.81481 | 13.75265 | 4.752618 |

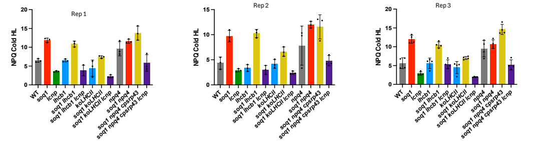


Fig1D

Rep 1

| Time (min) | WT | | | soq1 | | | koLHCII | | | soq1 koLHCII | | | soq1 koLHCII lcnp | | |
|---|---|---|---|---|---|---|---|---|---|---|---|---|---|---|---|
| 0 | 0 | 0 | 0 | 0 | 0 | 0 | 0 | 0 | 0 | 0 | 0 | 0 | 0 | 0 | 0 |
| 0.50483 | 1.03164 | 1.03037 | 1.15116 | 0.74248 | 1.01228 | 1.0908 | 0.30713 | 0.31775 | 0.34775 | 0.3598 | 0.36762 | 0.34536 | 0.31597 | 0.32524 | 0.30122 |
| 1.50967 | 1.59068 | 1.62556 | 1.91385 | 1.23425 | 1.66747 | 1.92602 | 0.514 | 0.526 | 0.54554 | 0.61084 | 0.62973 | 0.59042 | 0.53218 | 0.53186 | 0.50648 |
| 2.5145 | 1.73025 | 1.81297 | 2.21499 | 1.44066 | 1.91701 | 2.35708 | 0.63065 | 0.65617 | 0.67789 | 0.79737 | 0.80252 | 0.7692 | 0.6542 | 0.65941 | 0.62108 |
| 3.51934 | 1.92602 | 1.98065 | 2.32354 | 1.80345 | 2.1843 | 2.61932 | 0.72297 | 0.73944 | 0.76475 | 0.94069 | 0.94623 | 0.91265 | 0.72654 | 0.74857 | 0.6997 |
| 4.52417 | 2.12375 | 2.13748 | 2.43278 | 2.08917 | 2.32564 | 2.82131 | 0.79403 | 0.80736 | 0.81806 | 1.04492 | 1.08245 | 1.01108 | 0.77404 | 0.78788 | 0.7499 |
| 5.52901 | 2.21705 | 2.16465 | 2.45526 | 2.31797 | 2.43539 | 2.94372 | 0.83635 | 0.8572 | 0.86881 | 1.15129 | 1.19094 | 1.12384 | 0.81795 | 0.84687 | 0.8088 |
| 6.53384 | 2.29896 | 2.21202 | 2.49817 | 2.5051 | 2.55569 | 3.09591 | 0.88193 | 0.89121 | 0.9107 | 1.29178 | 1.29979 | 1.21969 | 0.85362 | 0.89608 | 0.85039 |
| 7.53868 | 2.31812 | 2.24169 | 2.59091 | 2.64732 | 2.65599 | 3.29155 | 0.91852 | 0.94534 | 0.93846 | 1.37252 | 1.38715 | 1.30452 | 0.87232 | 0.91337 | 0.86696 |
| 8.54351 | 2.31058 | 2.23935 | 2.63501 | 2.76293 | 2.7698 | 3.39265 | 0.96617 | 0.98696 | 0.96763 | 1.4524 | 1.50776 | 1.37504 | 0.88938 | 0.95903 | 0.91185 |
| 9.54834 | 2.35477 | 2.25207 | 2.67149 | 2.89581 | 2.89028 | 3.55115 | 0.9874 | 1.02245 | 0.99063 | 1.49975 | 1.5826 | 1.44653 | 0.91978 | 0.98267 | 0.94535 |
| 10.5532 | 2.34779 | 2.23049 | 2.69304 | 3.03386 | 3.01337 | 3.58152 | 1.02162 | 1.0456 | 1.01528 | 1.61382 | 1.63675 | 1.54676 | 0.94219 | 1.01427 | 0.96514 |
| 10.8085 | 1.27054 | 1.18058 | 1.72621 | 2.15004 | 2.17344 | 2.81812 | 0.79356 | 0.75202 | 0.80485 | 1.3594 | 1.58731 | 1.35674 | 0.64438 | 0.76556 | 0.75501 |
| 11.1463 | 0.84921 | 0.81025 | 1.21514 | 1.7678 | 1.79212 | 2.32842 | 0.73303 | 0.71286 | 0.72947 | 1.31923 | 1.47684 | 1.32884 | 0.58935 | 0.71695 | 0.6899 |
| 12.1508 | 0.56346 | 0.56215 | 0.77047 | 1.50223 | 1.42478 | 1.83435 | 0.64828 | 0.63245 | 0.63835 | 1.24688 | 1.39843 | 1.2003 | 0.50134 | 0.62153 | 0.60816 |
| 13.1553 | 0.50832 | 0.50912 | 0.67274 | 1.46893 | 1.36787 | 1.70111 | 0.60441 | 0.57269 | 0.57053 | 1.1779 | 1.34683 | 1.12975 | 0.46219 | 0.57035 | 0.55498 |
| 14.1598 | 0.4636 | 0.48018 | 0.62632 | 1.41182 | 1.31625 | 1.62096 | 0.54642 | 0.5313 | 0.52971 | 1.09812 | 1.25327 | 1.08241 | 0.41431 | 0.5011 | 0.51264 |
| 15.1643 | 0.45522 | 0.46042 | 0.58962 | 1.36685 | 1.28129 | 1.59575 | 0.5148 | 0.48772 | 0.50283 | 1.05411 | 1.20137 | 1.01881 | 0.40392 | 0.45938 | 0.45888 |
| 16.1688 | 0.42574 | 0.44282 | 0.56366 | 1.33953 | 1.26493 | 1.53033 | 0.49357 | 0.4642 | 0.47543 | 1.00885 | 1.16007 | 0.94823 | 0.38718 | 0.44308 | 0.43751 |
| 17.1733 | 0.40758 | 0.4273 | 0.55356 | 1.29297 | 1.23984 | 1.52035 | 0.45624 | 0.44863 | 0.44222 | 0.97619 | 1.13532 | 0.91756 | 0.37096 | 0.41665 | 0.42079 |
| 18.1778 | 0.39026 | 0.41167 | 0.53557 | 1.27332 | 1.21609 | 1.5013 | 0.42833 | 0.43531 | 0.40683 | 0.95551 | 1.068 | 0.86873 | 0.36336 | 0.37048 | 0.38723 |
| 19.1823 | 0.3767 | 0.39186 | 0.54238 | 1.2543 | 1.19516 | 1.456 | 0.40243 | 0.4211 | 0.3747 | 0.94838 | 1.06945 | 0.83232 | 0.36056 | 0.35954 | 0.36987 |
| 20.1868 | 0.3707 | 0.37678 | 0.53039 | 1.24013 | 1.17056 | 1.42539 | 0.38595 | 0.38369 | 0.36219 | 0.92337 | 1.0233 | 0.80735 | 0.33438 | 0.34879 | 0.34611 |
| 21.1913 | 0.35516 | 0.35909 | 0.53046 | 1.22984 | 1.12183 | 1.44027 | 0.38243 | 0.36408 | 0.35403 | 0.90936 | 0.98755 | 0.82105 | 0.32314 | 0.33356 | 0.32758 |

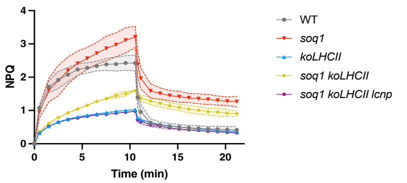


Rep2

| Time (min) | WT | | | soq1 | | | koLHCII | | | soq1 koLHCII | | | soq1 koLHCII lcnp | | |
|---|---|---|---|---|---|---|---|---|---|---|---|---|---|---|---|
| 0 | 0 | 0 | 0 | 0 | 0 | 0 | 0 | 0 | 0 | 0 | 0 | 0 | 0 | 0 | 0 |
| 0.50483 | 1.16269 | 1.10776 | 1.17643 | 1.17707 | 1.22034 | 1.15586 | 0.27914 | 0.3066775 | 0.30503 | 0.30239 | 0.36127 | 0.35251 | 0.31574 | 0.3414353 | 0.32416 |
| 1.50967 | 1.91567 | 1.79183 | 1.98411 | 1.90705 | 2.02819 | 2.03937 | 0.48329 | 0.4834514 | 0.51833 | 0.56094 | 0.62659 | 0.57139 | 0.51578 | 0.5463133 | 0.51558 |
| 2.5145 | 2.19431 | 2.07411 | 2.31357 | 2.19487 | 2.34341 | 2.4504 | 0.59942 | 0.5905555 | 0.63024 | 0.76055 | 0.82152 | 0.75894 | 0.64772 | 0.6690803 | 0.63545 |
| 3.51934 | 2.34862 | 2.21545 | 2.44736 | 2.39209 | 2.54051 | 2.63404 | 0.6751 | 0.6846557 | 0.71336 | 0.90737 | 0.95987 | 0.87741 | 0.73529 | 0.7557514 | 0.73939 |
| 4.52417 | 2.37239 | 2.2967 | 2.52932 | 2.57837 | 2.71002 | 2.79439 | 0.71134 | 0.7492822 | 0.76609 | 1.03003 | 1.11078 | 0.99568 | 0.8035 | 0.834477 | 0.80833 |
| 5.52901 | 2.48376 | 2.34974 | 2.63238 | 2.72693 | 2.82263 | 2.97766 | 0.7927 | 0.7985714 | 0.81334 | 1.16236 | 1.23228 | 1.09254 | 0.86326 | 0.8722028 | 0.86547 |
| 6.53384 | 2.54042 | 2.44602 | 2.66516 | 2.86679 | 2.90881 | 3.11359 | 0.84364 | 0.8721218 | 0.83918 | 1.27952 | 1.35673 | 1.18896 | 0.91428 | 0.9440243 | 0.92897 |
| 7.53868 | 2.59333 | 2.56277 | 2.70564 | 2.96525 | 2.98572 | 3.20154 | 0.85682 | 0.8895677 | 0.85988 | 1.36953 | 1.45205 | 1.28286 | 0.93511 | 0.9793865 | 0.9785 |
| 8.54351 | 2.61937 | 2.53906 | 2.75346 | 3.05707 | 3.09498 | 3.34791 | 0.88523 | 0.9191996 | 0.88597 | 1.49281 | 1.52893 | 1.3827 | 0.96098 | 0.99762 | 0.99688 |
| 9.54834 | 2.62486 | 2.59461 | 2.80661 | 3.11896 | 3.10259 | 3.39641 | 0.92486 | 0.9403595 | 0.9093 | 1.55016 | 1.62031 | 1.47159 | 0.99189 | 1.036839 | 1.01197 |
| 10.5532 | 2.70547 | 2.64047 | 2.79759 | 3.23856 | 3.21043 | 3.51902 | 0.93414 | 0.9655552 | 0.92496 | 1.65681 | 1.70734 | 1.54597 | 1.03205 | 1.039688 | 1.02648 |
| 10.8085 | 1.75274 | 2.1457 | 2.08767 | 2.71474 | 2.17909 | 2.97982 | 0.72476 | 0.7621358 | 0.64296 | 1.48934 | 1.54883 | 1.40006 | 0.80453 | 0.8189712 | 0.78434 |
| 11.1463 | 1.24312 | 1.57833 | 1.52494 | 2.1613 | 1.62604 | 2.46905 | 0.68634 | 0.721435 | 0.59208 | 1.45776 | 1.51274 | 1.35032 | 0.76208 | 0.7578666 | 0.71157 |
| 12.1508 | 0.70357 | 0.82511 | 0.81007 | 1.41567 | 1.12494 | 1.71884 | 0.59711 | 0.6458156 | 0.51416 | 1.34607 | 1.39597 | 1.23308 | 0.66359 | 0.6582908 | 0.61123 |
| 13.1553 | 0.6081 | 0.68403 | 0.70724 | 1.27014 | 1.00958 | 1.54321 | 0.53382 | 0.5903563 | 0.45962 | 1.30221 | 1.34833 | 1.15456 | 0.61158 | 0.600249 | 0.54299 |
| 14.1598 | 0.54473 | 0.62996 | 0.65604 | 1.16981 | 0.94837 | 1.47129 | 0.47536 | 0.5325162 | 0.40753 | 1.2357 | 1.28258 | 1.07064 | 0.56102 | 0.5469767 | 0.4866 |
| 15.1643 | 0.50928 | 0.60074 | 0.61778 | 1.13586 | 0.88724 | 1.40241 | 0.4405 | 0.4759495 | 0.38287 | 1.13251 | 1.20587 | 1.00935 | 0.52837 | 0.4996471 | 0.44919 |
| 16.1688 | 0.47272 | 0.57171 | 0.57545 | 1.07284 | 0.84767 | 1.35077 | 0.40525 | 0.4342681 | 0.35183 | 1.11173 | 1.17097 | 0.96711 | 0.48679 | 0.474207 | 0.41413 |
| 17.1733 | 0.45102 | 0.55134 | 0.5645 | 1.05851 | 0.81941 | 1.30843 | 0.37201 | 0.4023421 | 0.33059 | 1.04778 | 1.12779 | 0.90726 | 0.43673 | 0.4408499 | 0.39373 |
| 18.1778 | 0.43191 | 0.52638 | 0.52764 | 1.00348 | 0.77439 | 1.25645 | 0.3297 | 0.3791127 | 0.29641 | 0.9917 | 1.05183 | 0.82809 | 0.40232 | 0.3917818 | 0.37007 |
| 19.1823 | 0.42712 | 0.50651 | 0.48931 | 0.9689 | 0.75979 | 1.19803 | 0.31837 | 0.3585176 | 0.27966 | 0.97061 | 0.99579 | 0.80561 | 0.3827 | 0.3791729 | 0.34465 |
| 20.1868 | 0.42206 | 0.48535 | 0.47942 | 0.95469 | 0.72702 | 1.17655 | 0.30023 | 0.3507155 | 0.27113 | 0.94758 | 0.96966 | 0.7962 | 0.36881 | 0.3599717 | 0.33108 |
| 21.1913 | 0.41659 | 0.47569 | 0.46666 | 0.93641 | 0.72706 | 1.12366 | 0.29816 | 0.345552 | 0.2634 | 0.94492 | 0.95345 | 0.77702 | 0.37562 | 0.359159 | 0.33029 |

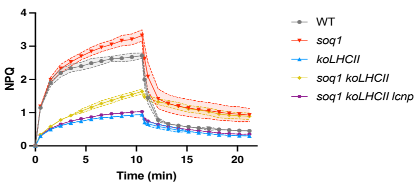


Rep 3

| Time (mib) | WT | | | soq1 | | | koLHCII | | | soq1 koLHCII | | | soq1 koLHCII lcnp | | |
|---|---|---|---|---|---|---|---|---|---|---|---|---|---|---|---|
| 0 | 0 | 0 | 0 | 0 | 0 | 0 | 0 | 0 | 0 | 0 | 0 | 0 | 0 | 0 | 0 |
| 0.50483 | 0.6355537 | 1.055509 | 1.263699 | 1.124963 | 1.206055 | 0.6988617 | 0.3159442 | 0.2671848 | 0.2955913 | 0.3156456 | 0.3117095 | 0.2532007 | 0.3467724 | 0.4257034 | 0.3174273 |
| 1.50967 | 1.388138 | 1.771968 | 1.940878 | 1.929976 | 1.933771 | 1.511127 | 0.5126051 | 0.4622588 | 0.4904828 | 0.5900184 | 0.5878568 | 0.4998272 | 0.5694957 | 0.6545324 | 0.5157994 |
| 2.5145 | 1.68843 | 2.051156 | 2.177185 | 2.256117 | 2.22831 | 1.964716 | 0.6389461 | 0.5878814 | 0.602227 | 0.7922143 | 0.7794604 | 0.6534936 | 0.6875449 | 0.7812812 | 0.6418377 |
| 3.51934 | 1.829732 | 2.177484 | 2.245317 | 2.448503 | 2.42059 | 2.213537 | 0.72642 | 0.6935293 | 0.6842412 | 0.9690781 | 0.9560986 | 0.7917206 | 0.8171122 | 0.8702461 | 0.7119142 |
| 4.52417 | 1.874316 | 2.254222 | 2.300897 | 2.555525 | 2.487034 | 2.412762 | 0.7856398 | 0.7755089 | 0.7437257 | 1.097568 | 1.088393 | 0.9041631 | 0.8627058 | 0.959283 | 0.7585329 |
| 5.52901 | 1.894781 | 2.289593 | 2.332479 | 2.680255 | 2.572055 | 2.524489 | 0.8387226 | 0.8181026 | 0.7898123 | 1.198067 | 1.181962 | 0.9736383 | 0.9158193 | 1.009181 | 0.8054945 |
| 6.53384 | 1.967222 | 2.342154 | 2.388914 | 2.786379 | 2.665073 | 2.672933 | 0.8754622 | 0.8656839 | 0.8254559 | 1.307481 | 1.279713 | 1.06038 | 0.9452314 | 1.046297 | 0.8307317 |
| 7.53868 | 1.995413 | 2.419417 | 2.430862 | 2.852053 | 2.744411 | 2.777568 | 0.9039718 | 0.8965361 | 0.8529165 | 1.371794 | 1.353379 | 1.110536 | 1.001866 | 1.099372 | 0.8663861 |
| 8.54351 | 2.027182 | 2.447705 | 2.437227 | 2.954358 | 2.776323 | 2.862653 | 0.9469062 | 0.9384477 | 0.8953709 | 1.443218 | 1.417565 | 1.177564 | 1.033184 | 1.148756 | 0.8783018 |
| 9.54834 | 2.057136 | 2.524152 | 2.488411 | 3.014436 | 2.854937 | 3.00923 | 0.9543018 | 0.9553013 | 0.9088113 | 1.515873 | 1.484331 | 1.222099 | 1.02228 | 1.1695 | 0.9141026 |
| 10.5532 | 2.090605 | 2.553446 | 2.540464 | 3.077111 | 2.96295 | 3.123558 | 0.9809431 | 0.9782109 | 0.9397563 | 1.568051 | 1.517497 | 1.28624 | 1.055247 | 1.219231 | 0.9319662 |
| 10.8085 | 1.319926 | 1.728854 | 1.69183 | 2.117651 | 1.96878 | 2.257405 | 0.7163858 | 0.7462668 | 0.6753297 | 1.485285 | 1.417614 | 1.130945 | 0.8115211 | 0.9637875 | 0.6345492 |
| 11.1463 | 0.956297 | 1.22385 | 1.176266 | 1.60671 | 1.516869 | 1.917616 | 0.6158431 | 0.6452234 | 0.591653 | 1.41172 | 1.346465 | 1.068468 | 0.7026341 | 0.8378522 | 0.5631734 |
| 12.1508 | 0.5797145 | 0.7167878 | 0.6539458 | 1.182261 | 1.102058 | 1.639193 | 0.4965538 | 0.5215379 | 0.5049952 | 1.2779 | 1.220373 | 0.9801661 | 0.5557687 | 0.6432594 | 0.5002297 |
| 13.1553 | 0.5325503 | 0.6463999 | 0.5612805 | 1.11563 | 1.026911 | 1.586229 | 0.4376608 | 0.4575522 | 0.4495744 | 1.206285 | 1.146204 | 0.9138586 | 0.4949276 | 0.5515825 | 0.4525321 |
| 14.1598 | 0.4802626 | 0.6045197 | 0.5210197 | 1.053343 | 1.0009 | 1.530871 | 0.3973498 | 0.4168731 | 0.4088579 | 1.158557 | 1.079436 | 0.8596326 | 0.4450516 | 0.4881431 | 0.4110664 |
| 15.1643 | 0.4649301 | 0.5850615 | 0.4905306 | 1.010123 | 0.9744635 | 1.51728 | 0.3798707 | 0.3900752 | 0.385157 | 1.104698 | 1.039907 | 0.8157358 | 0.4151167 | 0.4332905 | 0.3748802 |
| 16.1688 | 0.4501755 | 0.5681302 | 0.468234 | 0.9648067 | 0.9497319 | 1.499642 | 0.3596421 | 0.3588382 | 0.3696035 | 1.091402 | 1.002317 | 0.7948596 | 0.3842362 | 0.3867623 | 0.3530166 |
| 17.1733 | 0.4488063 | 0.5489935 | 0.451812 | 0.9247051 | 0.9338358 | 1.48827 | 0.3374216 | 0.3232237 | 0.3591379 | 1.032602 | 0.9570689 | 0.7657132 | 0.363252 | 0.365889 | 0.3302629 |
| 18.1778 | 0.4394132 | 0.5235168 | 0.4275225 | 0.9028957 | 0.9056707 | 1.477147 | 0.3103375 | 0.2832394 | 0.3436423 | 0.9810134 | 0.8997012 | 0.7516348 | 0.3367545 | 0.3483786 | 0.3199203 |
| 19.1823 | 0.423386 | 0.491839 | 0.3992118 | 0.887763 | 0.8829742 | 1.446843 | 0.2831875 | 0.2649568 | 0.3325365 | 0.9115717 | 0.8223134 | 0.7400916 | 0.3051307 | 0.3344704 | 0.3050276 |
| 20.1868 | 0.4257021 | 0.46698 | 0.3777049 | 0.8675089 | 0.8228216 | 1.455215 | 0.2607136 | 0.2551211 | 0.3077749 | 0.8664469 | 0.7797299 | 0.7231667 | 0.2721293 | 0.3320247 | 0.3051788 |
| 21.1913 | 0.419368 | 0.4443707 | 0.3563268 | 0.8542453 | 0.7891608 | 1.442521 | 0.243231 | 0.2488546 | 0.2802658 | 0.8302234 | 0.7724198 | 0.7167614 | 0.2439815 | 0.3274906 | 0.2813894 |

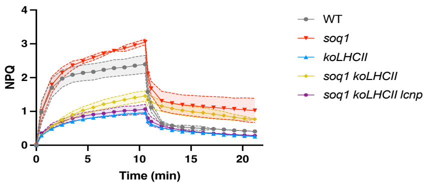

Figure 2 A, B and C

Normalized data

Trimer

| | soq1 lhcb1 | soq1 lhcb1 lcnp | soq1 npq4 cpsrp43 | soq1 npq4 cpsrp43 lcnp |
|---|---|---|---|---|
| Rep 1 | 0.71 | 1.02 | 0.67 | 0.97 |
| Rep 2 | 0.73 | 1.06 | 0.76 | 1.08 |
| Rep 3 | 0.65 | 0.92 | 0.82 | 0.95 |

Monomer

| | soq1 koLHCII | soq1 koLHCII lcnp | soq1 lhcb1 | soq1 lhcb1 lcnp | soq1 npq4 cpsrp43 | soq1 npq4 cpsrp43 lcnp |
|---|---|---|---|---|---|---|
| Rep 1 | 0.78 | 0.93 | 0.7 | 0.96 | 0.73 | 1.04 |
| Rep 2 | 0.7 | 0.96 | 0.63 | 0.97 | 0.61 | 1.03 |
| Rep 3 | 0.84 | 1.12 | 0.85 | 1.07 | 0.63 | 0.93 |

Raw data

Green band

Trimer

| | soq1 lhcb1 | soq1 lhcb1 lcnp | soq1 npq4 cpsrp43 | soq1 npq4 cpsrp43 lcnp |
|---|---|---|---|---|
| Rep 1 | 1160 | 957 | 1959 | 1770 |
| Rep 2 | 1157 | 944 | 2322 | 2184 |
| Rep 3 | 1245 | 1066 | 2147 | 2158 |

Monomer

| | soq1 koLHCII | soq1 koLHCII lcnp | soq1 lhcb1 | soq1 lhcb1 lcnp | soq1 npq4 cpsrp43 | soq1 npq4 cpsrp43 lcnp |
|---|---|---|---|---|---|---|
| Rep 1 | 945 | 966 | 1403 | 1090 | 364 | 331 |
| Rep 2 | 1251 | 1035 | 1601 | 1180 | 505 | 375 |
| Rep 3 | 1093 | 894 | 1216 | 1077 | 494 | 365 |

Fluorescence

Trimer

| | soq1 lhcb1 | soq1 lhcb1 lcnp | soq1 npq4 cpsrp43 | soq1 npq4 cpsrp43 lcnp |
|---|---|---|---|---|
| Rep 1 | 375.0453 | 442.3736 | 283.4478 | 368.0559 |
| Rep 2 | 382.9078 | 455.2534 | 377.9633 | 509.2353 |
| Rep 3 | 366.0556 | 444.2 | 379.3 | 439.5909 |

Monomer

| | soq1 koLHCII | soq1 koLHCII lcnp | soq1 lhcb1 | soq1 lhcb1 lcnp | soq1 npq4 cpsrp43 | soq1 npq4 cpsrp43 lcnp |
|---|---|---|---|---|---|---|
| Rep 1 | 329.18 | 397.3145 | 427.3289 | 456.6718 | 223.9285 | 288.495 |
| Rep 2 | 389.33 | 439.5497 | 436.1424 | 497.772 | 259.8954 | 325.13 |
| Rep 3 | 406.55 | 444.6728 | 450.4506 | 501.0679 | 261.3642 | 285.759 |

Rep 1

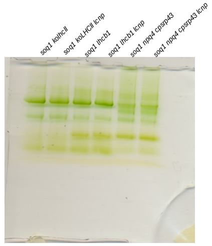


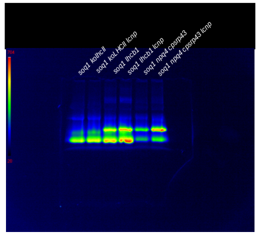

Rep 2

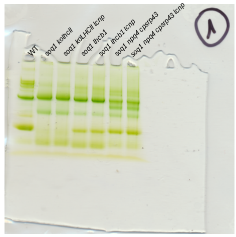

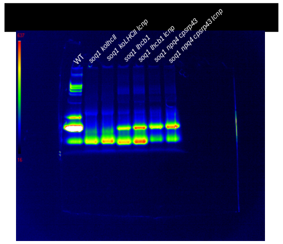

Rep 3

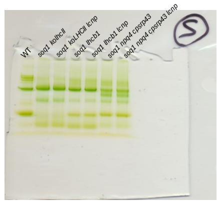

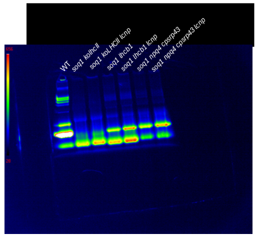

Figure 2 D

Trimer

| | Rep 1 | | Rep 2 | | Rep 3 | | Rep 4 | | Rep 5 | |
|---|---|---|---|---|---|---|---|---|---|---|
| | soq1 roqh1 | soq1 roqh1 lcnp | soq1 roqh1 | soq1 roqh1 lcnp | soq1 roqh1 | soq1 roqh1 lcnp | soq1 roqh1 | soq1 roqh1 lcnp | soq1 roqh1 | soq1 roqh1 lcnp |
| Raw value 680nm | 1033651.53 | 1989130.998 | 966100.995 | 1933424.757 | 1173496.14 | 2079309.194 | 1320345.24 | 2076368.202 | 1394991.97 | 2185595.048 |
| Normalized | 0.50354093 | 0.969000532 | 0.47063385 | 0.941863367 | 0.57166591 | 1.012930631 | 0.64320311 | 1.011497933 | 0.67956709 | 1.064707537 |

Monomer

| | Rep 1 | | Rep 2 | | Rep 3 | | Rep 4 | | Rep 5 | |
|---|---|---|---|---|---|---|---|---|---|---|
| | soq1 roqh1 | soq1 roqh1 lcnp | soq1 roqh1 | soq1 roqh1 lcnp | soq1 roqh1 | soq1 roqh1 lcnp | soq1 roqh1 | soq1 roqh1 lcnp | soq1 roqh1 | soq1 roqh1 lcnp |
| Raw value 680nm | 1440171.94 | 2098469.264 | 1411752.53 | 1912389.895 | 1614919.67 | 2446466.513 | 1919132.1 | 2251629.063 | 1586570.04 | 2411659.587 |
| Normalized | 0.64752355 | 0.9435042 | 0.63474575 | 0.859840041 | 0.72609283 | 1.099969139 | 0.86287144 | 1.012367212 | 0.7133464 | 1.084319408 |

Figure 3 A, B, C

Rep 1

| Time (min) | *soq1* | | | NoM | | | soq1 NoM | | | lhcb4 | | |
|---|---|---|---|---|---|---|---|---|---|---|---|---|
| 0 | 0 | 0 | 0 | 0 | 0 | 0 | 0 | 0 | 0 | 0 | 0 | 0 |
| 0.50483 | 0.74248 | 1.01228 | 1.0908 | 0.97137 | 1.04634 | 1.10967 | 1.28512 | 1.01707 | 1.08504 | 0.79054 | 0.69989 | 0.93351 |
| 1.50967 | 1.23425 | 1.66747 | 1.92602 | 0.70194 | 0.76547 | 0.9211 | 1.49618 | 0.94352 | 1.31212 | 0.8501 | 0.84652 | 1.18288 |
| 2.5145 | 1.44066 | 1.91701 | 2.35708 | 0.76977 | 0.76424 | 0.97799 | 1.63711 | 1.01046 | 1.45312 | 0.87102 | 0.87114 | 1.24087 |
| 3.51934 | 1.80345 | 2.1843 | 2.61932 | 1.02643 | 0.92764 | 1.19665 | 1.77194 | 1.23904 | 1.67405 | 0.96918 | 0.99664 | 1.28324 |
| 4.52417 | 2.08917 | 2.32564 | 2.82131 | 1.42337 | 1.22341 | 1.50923 | 1.97393 | 1.50955 | 1.88543 | 1.09421 | 1.15382 | 1.35349 |
| 5.52901 | 2.31797 | 2.43539 | 2.94372 | 1.86585 | 1.68789 | 1.84192 | 2.20027 | 1.80611 | 2.17342 | 1.19063 | 1.26517 | 1.44296 |
| 6.53384 | 2.5051 | 2.55569 | 3.09591 | 2.15031 | 2.08244 | 2.09692 | 2.44634 | 2.11523 | 2.43197 | 1.24805 | 1.29939 | 1.53132 |
| 7.53868 | 2.64732 | 2.65599 | 3.29155 | 2.32596 | 2.36112 | 2.30356 | 2.64249 | 2.4109 | 2.65959 | 1.30128 | 1.34508 | 1.58481 |
| 8.54351 | 2.76293 | 2.7698 | 3.39265 | 2.40819 | 2.45609 | 2.40689 | 2.78875 | 2.58752 | 2.76094 | 1.33785 | 1.35303 | 1.58042 |
| 9.54834 | 2.89581 | 2.89028 | 3.55115 | 2.47253 | 2.52199 | 2.50689 | 2.92556 | 2.78841 | 2.94396 | 1.37565 | 1.39281 | 1.6374 |
| 10.5532 | 3.03386 | 3.01337 | 3.58152 | 2.49698 | 2.56571 | 2.53594 | 3.02069 | 2.85834 | 2.99592 | 1.37297 | 1.39588 | 1.63078 |
| 10.8085 | 2.15004 | 2.17344 | 2.81812 | 0.68737 | 0.60353 | 1.04632 | 1.83442 | 1.66468 | 1.8005 | 0.52345 | 0.54236 | 0.70683 |
| 11.1463 | 1.7678 | 1.79212 | 2.32842 | 0.38518 | 0.34539 | 0.58095 | 1.34437 | 1.26278 | 1.40457 | 0.34665 | 0.43912 | 0.48304 |
| 12.1508 | 1.50223 | 1.42478 | 1.83435 | 0.2223 | 0.21005 | 0.23279 | 0.91698 | 0.91282 | 1.00766 | 0.2952 | 0.3779 | 0.39712 |
| 13.1553 | 1.46893 | 1.36787 | 1.70111 | 0.19015 | 0.17477 | 0.19512 | 0.85296 | 0.8675 | 0.96594 | 0.28924 | 0.36442 | 0.37482 |
| 14.1598 | 1.41182 | 1.31625 | 1.62096 | 0.17818 | 0.15774 | 0.17647 | 0.82052 | 0.85103 | 0.9285 | 0.31312 | 0.35886 | 0.36294 |
| 15.1643 | 1.36685 | 1.28129 | 1.59575 | 0.16379 | 0.1478 | 0.17164 | 0.78391 | 0.82722 | 0.91882 | 0.31894 | 0.34975 | 0.35111 |
| 16.1688 | 1.33953 | 1.26493 | 1.53033 | 0.1611 | 0.13868 | 0.16281 | 0.75332 | 0.80761 | 0.89306 | 0.33546 | 0.33426 | 0.34453 |
| 17.1733 | 1.29297 | 1.23984 | 1.52035 | 0.15428 | 0.13879 | 0.15314 | 0.74605 | 0.7966 | 0.89364 | 0.34245 | 0.32477 | 0.34398 |
| 18.1778 | 1.27332 | 1.21609 | 1.5013 | 0.14658 | 0.13754 | 0.14758 | 0.7208 | 0.79866 | 0.87442 | 0.3477 | 0.32253 | 0.33526 |
| 19.1823 | 1.2543 | 1.19516 | 1.456 | 0.13877 | 0.13551 | 0.14487 | 0.71905 | 0.78811 | 0.86964 | 0.35102 | 0.32808 | 0.34196 |
| 20.1868 | 1.24013 | 1.17056 | 1.42539 | 0.1352 | 0.13852 | 0.14689 | 0.71146 | 0.79019 | 0.86615 | 0.36144 | 0.3453 | 0.34208 |
| 21.1913 | 1.22984 | 1.12183 | 1.44027 | 0.12891 | 0.14205 | 0.15243 | 0.72927 | 0.79411 | 0.87081 | 0.37394 | 0.36431 | 0.34523 |

| Time (min) | soq1 lhcb4 | | | lhcb5 | | | soq1 lhcb5 | | | WT | | |
|---|---|---|---|---|---|---|---|---|---|---|---|---|
| 0 | 0 | 0 | 0 | 0 | 0 | 0 | 0 | 0 | 0 | 0 | 0 | 0 |
| 0.50483 | 0.84769 | 0.828 | 0.96313 | 1.0536 | 1.25632 | 1.06199 | 1.2877 | 1.39939 | 1.22042 | 1.03164 | 1.03037 | 1.15116 |
| 1.50967 | 0.75821 | 0.70345 | 1.01086 | 1.60833 | 2.04114 | 1.71449 | 2.097 | 2.23622 | 1.92255 | 1.59068 | 1.62556 | 1.91385 |
| 2.5145 | 0.87361 | 0.73934 | 1.03888 | 1.91592 | 2.29656 | 1.96035 | 2.3825 | 2.57811 | 2.21606 | 1.73025 | 1.81297 | 2.21499 |
| 3.51934 | 1.10344 | 0.90071 | 1.13409 | 2.11255 | 2.38234 | 2.05575 | 2.59773 | 2.70329 | 2.39658 | 1.92602 | 1.98065 | 2.32354 |
| 4.52417 | 1.41067 | 1.11993 | 1.296 | 2.25721 | 2.44145 | 2.18333 | 2.69948 | 2.79389 | 2.49259 | 2.12375 | 2.13748 | 2.43278 |
| 5.52901 | 1.70809 | 1.37615 | 1.44776 | 2.38325 | 2.47084 | 2.35153 | 2.8258 | 2.93989 | 2.58687 | 2.21705 | 2.16465 | 2.45526 |
| 6.53384 | 1.90833 | 1.63291 | 1.61757 | 2.47319 | 2.54237 | 2.3718 | 2.92649 | 2.99033 | 2.63645 | 2.29896 | 2.21202 | 2.49817 |
| 7.53868 | 2.11977 | 1.88239 | 1.73873 | 2.53445 | 2.60421 | 2.41163 | 2.97856 | 3.08268 | 2.76623 | 2.31812 | 2.24169 | 2.59091 |
| 8.54351 | 2.24402 | 2.08034 | 1.83601 | 2.55591 | 2.63021 | 2.48637 | 3.07732 | 3.06873 | 2.83823 | 2.31058 | 2.23935 | 2.63501 |
| 9.54834 | 2.36844 | 2.2516 | 1.92624 | 2.64019 | 2.69293 | 2.52686 | 3.201 | 3.19696 | 2.8457 | 2.35477 | 2.25207 | 2.67149 |
| 10.5532 | 2.44757 | 2.38053 | 2.02158 | 2.62261 | 2.68021 | 2.55452 | 3.23559 | 3.19772 | 2.96498 | 2.34779 | 2.23049 | 2.69304 |
| 10.8085 | 1.33663 | 1.44341 | 1.11767 | 1.53791 | 1.83714 | 1.92999 | 2.61387 | 2.55241 | 2.50594 | 1.27054 | 1.18058 | 1.72621 |
| 11.1463 | 1.28349 | 1.37057 | 0.94337 | 1.07655 | 1.28851 | 1.40564 | 2.103 | 1.91652 | 2.02966 | 0.84921 | 0.81025 | 1.21514 |
| 12.1508 | 1.2087 | 1.31618 | 0.85171 | 0.58849 | 0.61545 | 0.64947 | 1.38829 | 1.17032 | 1.32678 | 0.56346 | 0.56215 | 0.77047 |
| 13.1553 | 1.1334 | 1.22745 | 0.80138 | 0.49099 | 0.52494 | 0.53813 | 1.22358 | 1.01847 | 1.17671 | 0.50832 | 0.50912 | 0.67274 |
| 14.1598 | 1.0925 | 1.16968 | 0.78725 | 0.4564 | 0.48171 | 0.49965 | 1.1648 | 0.96612 | 1.10167 | 0.4636 | 0.48018 | 0.62632 |
| 15.1643 | 1.09318 | 1.18139 | 0.80241 | 0.45075 | 0.4776 | 0.48866 | 1.13243 | 0.91742 | 1.08082 | 0.45522 | 0.46042 | 0.58962 |
| 16.1688 | 1.08778 | 1.14574 | 0.79926 | 0.44722 | 0.4583 | 0.45844 | 1.08061 | 0.87139 | 1.03354 | 0.42574 | 0.44282 | 0.56366 |
| 17.1733 | 1.08494 | 1.16545 | 0.80778 | 0.45394 | 0.44442 | 0.45437 | 1.03058 | 0.86801 | 0.98984 | 0.40758 | 0.4273 | 0.55356 |
| 18.1778 | 1.08618 | 1.17545 | 0.8011 | 0.45145 | 0.42509 | 0.45235 | 0.98001 | 0.8401 | 0.95794 | 0.39026 | 0.41167 | 0.53557 |
| 19.1823 | 1.08628 | 1.16649 | 0.81372 | 0.46853 | 0.41741 | 0.44732 | 0.96831 | 0.82514 | 0.93241 | 0.3767 | 0.39186 | 0.54238 |
| 20.1868 | 1.10012 | 1.19032 | 0.81106 | 0.46832 | 0.40185 | 0.44971 | 0.94865 | 0.80738 | 0.93368 | 0.3707 | 0.37678 | 0.53039 |
| 21.1913 | 1.11731 | 1.21063 | 0.81922 | 0.48381 | 0.39618 | 0.45452 | 0.93573 | 0.81094 | 0.91819 | 0.35516 | 0.35909 | 0.53046 |

Rep 2

| Time (min) | *soq1* | | | NoM | | | soq1 NoM | | | lhcb4 | | |
|---|---|---|---|---|---|---|---|---|---|---|---|---|
| 0 | 0 | 0 | 0 | 0 | 0 | 0 | 0 | 0 | 0 | 0 | 0 | 0 |
| 0.50483 | 1.17707 | 1.22034 | 1.15586 | 0.99624 | 1.43768 | 1.36231 | 0.95551 | 1.00962 | 0.77043 | 0.88101 | 0.89948 | 1.07009 |
| 1.50967 | 1.90705 | 2.02819 | 2.03937 | 0.86851 | 1.1311 | 1.00038 | 0.7304 | 0.733 | 0.57358 | 0.89511 | 1.00599 | 1.21997 |
| 2.5145 | 2.19487 | 2.34341 | 2.4504 | 1.23997 | 1.11081 | 0.96152 | 0.74611 | 0.8519 | 0.66251 | 1.05297 | 1.11711 | 1.15488 |
| 3.51934 | 2.39209 | 2.54051 | 2.63404 | 1.78053 | 1.29768 | 1.10859 | 0.93018 | 1.14423 | 0.92143 | 1.23684 | 1.28242 | 1.30585 |
| 4.52417 | 2.57837 | 2.71002 | 2.79439 | 2.13492 | 1.64912 | 1.55773 | 1.25746 | 1.54953 | 1.4289 | 1.35402 | 1.41871 | 1.5782 |
| 5.52901 | 2.72693 | 2.82263 | 2.97766 | 2.31112 | 2.14454 | 2.10831 | 1.69107 | 2.05016 | 1.92355 | 1.37401 | 1.46453 | 1.69447 |
| 6.53384 | 2.86679 | 2.90881 | 3.11359 | 2.36202 | 2.57768 | 2.43227 | 2.13339 | 2.44284 | 2.25261 | 1.42208 | 1.51227 | 1.75263 |
| 7.53868 | 2.96525 | 2.98572 | 3.20154 | 2.39831 | 2.71581 | 2.59617 | 2.43575 | 2.70171 | 2.47196 | 1.39806 | 1.53551 | 1.7596 |
| 8.54351 | 3.05707 | 3.09498 | 3.34791 | 2.4464 | 2.7682 | 2.69067 | 2.62878 | 2.86418 | 2.58512 | 1.42893 | 1.56372 | 1.77613 |
| 9.54834 | 3.11896 | 3.10259 | 3.39641 | 2.46229 | 2.85554 | 2.74208 | 2.75811 | 3.01721 | 2.74534 | 1.47088 | 1.58703 | 1.78294 |
| 10.5532 | 3.23856 | 3.21043 | 3.51902 | 2.46106 | 2.83752 | 2.76952 | 2.86227 | 3.11872 | 2.85098 | 1.49457 | 1.58687 | 1.81327 |
| 10.8085 | 2.71474 | 2.17909 | 2.97982 | 0.95051 | 1.27794 | 1.136 | 1.32507 | 1.72263 | 1.27287 | 0.62134 | 0.48562 | 0.88659 |
| 11.1463 | 2.1613 | 1.62604 | 2.46905 | 0.55801 | 0.64524 | 0.5896 | 1.07662 | 1.3308 | 1.00071 | 0.43615 | 0.38635 | 0.57293 |
| 12.1508 | 1.41567 | 1.12494 | 1.71884 | 0.29426 | 0.27641 | 0.29064 | 0.9414 | 1.09825 | 0.82524 | 0.32136 | 0.29841 | 0.36934 |
| 13.1553 | 1.27014 | 1.00958 | 1.54321 | 0.25623 | 0.22807 | 0.23685 | 0.89461 | 1.05783 | 0.80984 | 0.30943 | 0.28699 | 0.32272 |
| 14.1598 | 1.16981 | 0.94837 | 1.47129 | 0.24018 | 0.19869 | 0.20433 | 0.87679 | 1.03202 | 0.79815 | 0.31491 | 0.2934 | 0.29641 |
| 15.1643 | 1.13586 | 0.88724 | 1.40241 | 0.20708 | 0.17558 | 0.18474 | 0.86172 | 1.00962 | 0.7847 | 0.31266 | 0.2966 | 0.29065 |
| 16.1688 | 1.07284 | 0.84767 | 1.35077 | 0.18799 | 0.16839 | 0.17184 | 0.86125 | 0.98655 | 0.77401 | 0.31843 | 0.296 | 0.28436 |
| 17.1733 | 1.05851 | 0.81941 | 1.30843 | 0.18399 | 0.16182 | 0.17188 | 0.84173 | 0.9824 | 0.77429 | 0.33113 | 0.30937 | 0.29189 |
| 18.1778 | 1.00348 | 0.77439 | 1.25645 | 0.18133 | 0.16843 | 0.17779 | 0.83304 | 0.96742 | 0.76115 | 0.34399 | 0.32081 | 0.28943 |
| 19.1823 | 0.9689 | 0.75979 | 1.19803 | 0.18588 | 0.17073 | 0.18067 | 0.83216 | 0.94045 | 0.75271 | 0.36148 | 0.34079 | 0.28461 |
| 20.1868 | 0.95469 | 0.72702 | 1.17655 | 0.19449 | 0.17857 | 0.18779 | 0.82428 | 0.91407 | 0.76647 | 0.37345 | 0.34469 | 0.29948 |
| 21.1913 | 0.93641 | 0.72706 | 1.12366 | 0.20107 | 0.18852 | 0.20241 | 0.82519 | 0.92012 | 0.76662 | 0.38582 | 0.36161 | 0.31139 |

| Time (min) | soq1 lhcb4 | | | lhcb5 | | | soq1 lhcb5 | | | WT | | |
|---|---|---|---|---|---|---|---|---|---|---|---|---|
| 0 | 0 | 0 | 0 | 0 | 0 | 0 | 0 | 0 | 0 | 0 | 0 | 0 |
| 0.50483 | 0.81988 | 1.04227 | 0.78556 | 1.07196 | 1.25674 | 1.10847 | 1.29042 | 1.42303 | 1.28382 | 1.16269 | 1.10776 | 1.17643 |
| 1.50967 | 0.78423 | 1.18379 | 1.00564 | 1.86109 | 1.89545 | 1.55612 | 2.07108 | 2.28262 | 2.04618 | 1.91567 | 1.79183 | 1.98411 |
| 2.5145 | 0.96929 | 1.25302 | 1.13829 | 2.19931 | 2.15951 | 1.7889 | 2.41174 | 2.59442 | 2.32529 | 2.19431 | 2.07411 | 2.31357 |
| 3.51934 | 1.18099 | 1.50346 | 1.33608 | 2.32527 | 2.2586 | 1.95403 | 2.52764 | 2.75718 | 2.50399 | 2.34862 | 2.21545 | 2.44736 |
| 4.52417 | 1.42232 | 1.76768 | 1.55291 | 2.48784 | 2.33756 | 2.04392 | 2.70101 | 2.8554 | 2.68342 | 2.37239 | 2.2967 | 2.52932 |
| 5.52901 | 1.59607 | 1.86833 | 1.76834 | 2.57395 | 2.41233 | 2.14631 | 2.87934 | 2.95298 | 2.81766 | 2.48376 | 2.34974 | 2.63238 |
| 6.53384 | 1.72999 | 1.96552 | 1.9812 | 2.63061 | 2.47259 | 2.25243 | 2.9306 | 3.00916 | 2.94575 | 2.54042 | 2.44602 | 2.66516 |
| 7.53868 | 1.84362 | 2.02157 | 2.13221 | 2.66318 | 2.4775 | 2.27937 | 3.03273 | 3.07859 | 2.96794 | 2.59333 | 2.56277 | 2.70564 |
| 8.54351 | 1.94803 | 2.07173 | 2.24544 | 2.70324 | 2.52944 | 2.28664 | 3.07412 | 3.1137 | 3.06971 | 2.61937 | 2.53906 | 2.75346 |
| 9.54834 | 2.05028 | 2.14098 | 2.38554 | 2.7389 | 2.56185 | 2.38553 | 3.17509 | 3.15247 | 3.16379 | 2.62486 | 2.59461 | 2.80661 |
| 10.5532 | 2.12144 | 2.13843 | 2.47576 | 2.73639 | 2.58639 | 2.39116 | 3.23902 | 3.17235 | 3.16506 | 2.70547 | 2.64047 | 2.79759 |
| 10.8085 | 1.36428 | 1.316 | 1.65301 | 1.70357 | 1.79709 | 1.61454 | 2.44952 | 2.3419 | 2.47408 | 1.75274 | 2.1457 | 2.08767 |
| 11.1463 | 1.12425 | 0.98212 | 1.49666 | 1.20746 | 1.25461 | 1.13623 | 1.85157 | 1.69821 | 1.93718 | 1.24312 | 1.57833 | 1.52494 |
| 12.1508 | 1.00885 | 0.76451 | 1.40765 | 0.60777 | 0.51455 | 0.52972 | 1.13859 | 0.93818 | 1.19963 | 0.70357 | 0.82511 | 0.81007 |
| 13.1553 | 0.95456 | 0.70367 | 1.37234 | 0.4943 | 0.42796 | 0.43352 | 0.99123 | 0.80455 | 1.06624 | 0.6081 | 0.68403 | 0.70724 |
| 14.1598 | 0.9437 | 0.68217 | 1.30303 | 0.45156 | 0.42289 | 0.40479 | 0.93046 | 0.75835 | 0.99529 | 0.54473 | 0.62996 | 0.65604 |
| 15.1643 | 0.93045 | 0.66022 | 1.24547 | 0.43296 | 0.40807 | 0.38655 | 0.87286 | 0.70955 | 0.93539 | 0.50928 | 0.60074 | 0.61778 |
| 16.1688 | 0.92493 | 0.62715 | 1.22752 | 0.42954 | 0.40513 | 0.37859 | 0.81209 | 0.66876 | 0.92173 | 0.47272 | 0.57171 | 0.57545 |
| 17.1733 | 0.94334 | 0.60932 | 1.21232 | 0.41697 | 0.3839 | 0.37349 | 0.78613 | 0.64326 | 0.8798 | 0.45102 | 0.55134 | 0.5645 |
| 18.1778 | 0.94661 | 0.59643 | 1.18288 | 0.42228 | 0.38085 | 0.36373 | 0.76496 | 0.61376 | 0.85674 | 0.43191 | 0.52638 | 0.52764 |
| 19.1823 | 0.93338 | 0.58543 | 1.15887 | 0.41767 | 0.37472 | 0.36348 | 0.73342 | 0.60559 | 0.8227 | 0.42712 | 0.50651 | 0.48931 |
| 20.1868 | 0.94849 | 0.60055 | 1.15561 | 0.42737 | 0.38006 | 0.36346 | 0.72513 | 0.60236 | 0.83795 | 0.42206 | 0.48535 | 0.47942 |
| 21.1913 | 0.95292 | 0.59502 | 1.17533 | 0.44398 | 0.38712 | 0.36931 | 0.72039 | 0.60191 | 0.81808 | 0.41659 | 0.47569 | 0.46666 |

Rep 3

| Time (min) | *soq1* | | | NoM | | | soq1 NoM | | | lhcb4 | | |
|---|---|---|---|---|---|---|---|---|---|---|---|---|
| 0 | 0 | 0 | 0 | 0 | 0 | 0 | 0 | 0 | 0 | 0 | 0 | 0 |
| 0.50483 | 1.124963 | 1.206055 | 0.6988617 | 1.232619 | 0.8641459 | 0.9410954 | 1.090183 | 1.313801 | 0.9475612 | 0.4183571 | 0.8844355 | 0.7527773 |
| 1.50967 | 1.929976 | 1.933771 | 1.511127 | 0.8150716 | 0.7535405 | 0.7956012 | 0.8623232 | 1.243943 | 0.8348687 | 0.7891983 | 1.260387 | 1.051627 |
| 2.5145 | 2.256117 | 2.22831 | 1.964716 | 0.6408886 | 0.6372229 | 0.7766458 | 0.8538069 | 1.267929 | 0.8490708 | 0.8637214 | 1.377844 | 1.170352 |
| 3.51934 | 2.448503 | 2.42059 | 2.213537 | 0.7186786 | 0.7682648 | 0.977667 | 1.059531 | 1.461014 | 1.028552 | 1.0077 | 1.424145 | 1.242621 |
| 4.52417 | 2.555525 | 2.487034 | 2.412762 | 0.9848011 | 1.055664 | 1.31403 | 1.423263 | 1.789266 | 1.315957 | 1.178811 | 1.480807 | 1.330266 |
| 5.52901 | 2.680255 | 2.572055 | 2.524489 | 1.3698 | 1.399798 | 1.676007 | 1.800147 | 2.127964 | 1.649373 | 1.303968 | 1.535107 | 1.421918 |
| 6.53384 | 2.786379 | 2.665073 | 2.672933 | 1.852378 | 1.745327 | 2.006639 | 2.18182 | 2.496086 | 1.958063 | 1.40813 | 1.576334 | 1.471739 |
| 7.53868 | 2.852053 | 2.744411 | 2.777568 | 2.220734 | 2.044849 | 2.233653 | 2.478919 | 2.741369 | 2.248683 | 1.481776 | 1.603018 | 1.515605 |
| 8.54351 | 2.954358 | 2.776323 | 2.862653 | 2.410581 | 2.233579 | 2.379228 | 2.698678 | 2.9246 | 2.472487 | 1.542935 | 1.639081 | 1.542557 |
| 9.54834 | 3.014436 | 2.854937 | 3.00923 | 2.501688 | 2.327472 | 2.456985 | 2.888197 | 3.018792 | 2.634405 | 1.585455 | 1.653186 | 1.556827 |
| 10.5532 | 3.077111 | 2.96295 | 3.123558 | 2.544174 | 2.413031 | 2.515046 | 3.005449 | 3.078657 | 2.801234 | 1.625111 | 1.67063 | 1.577578 |
| 10.8085 | 2.117651 | 1.96878 | 2.257405 | 0.6762259 | 0.6810675 | 0.7733859 | 1.544541 | 1.712468 | 1.631679 | 0.6612036 | 0.8022087 | 0.6530057 |
| 11.1463 | 1.60671 | 1.516869 | 1.917616 | 0.337749 | 0.3610638 | 0.4279364 | 1.043398 | 1.217809 | 1.108483 | 0.503043 | 0.5534985 | 0.4794171 |
| 12.1508 | 1.182261 | 1.102058 | 1.639193 | 0.1929713 | 0.2129784 | 0.196118 | 0.7423036 | 0.7817808 | 0.7447452 | 0.414747 | 0.4155521 | 0.3848771 |
| 13.1553 | 1.11563 | 1.026911 | 1.586229 | 0.164609 | 0.1824524 | 0.1645088 | 0.7063322 | 0.7230313 | 0.6947425 | 0.3881034 | 0.3896722 | 0.3610115 |
| 14.1598 | 1.053343 | 1.0009 | 1.530871 | 0.145554 | 0.168236 | 0.150337 | 0.6785372 | 0.6859396 | 0.6797313 | 0.375999 | 0.3754874 | 0.3599994 |
| 15.1643 | 1.010123 | 0.9744635 | 1.51728 | 0.1345464 | 0.1602973 | 0.1419902 | 0.6684471 | 0.6758931 | 0.6621494 | 0.369808 | 0.3696582 | 0.3583794 |
| 16.1688 | 0.9648067 | 0.9497319 | 1.499642 | 0.1273092 | 0.1536163 | 0.1315096 | 0.6477914 | 0.6516408 | 0.6436586 | 0.3702378 | 0.3463524 | 0.3495016 |
| 17.1733 | 0.9247051 | 0.9338358 | 1.48827 | 0.1221345 | 0.153457 | 0.1287741 | 0.6406881 | 0.6356132 | 0.6386173 | 0.3634103 | 0.3293508 | 0.3469031 |
| 18.1778 | 0.9028957 | 0.9056707 | 1.477147 | 0.1129887 | 0.1495423 | 0.1257892 | 0.6305754 | 0.6155103 | 0.6287296 | 0.3605233 | 0.3150378 | 0.3421192 |
| 19.1823 | 0.887763 | 0.8829742 | 1.446843 | 0.1095801 | 0.1496171 | 0.12496 | 0.6257836 | 0.6024109 | 0.6220562 | 0.3611355 | 0.3111528 | 0.3421648 |
| 20.1868 | 0.8675089 | 0.8228216 | 1.455215 | 0.1053532 | 0.1485551 | 0.124444 | 0.6269792 | 0.5917295 | 0.6191429 | 0.3598681 | 0.3123669 | 0.3490618 |
| 21.1913 | 0.8542453 | 0.7891608 | 1.442521 | 0.1004969 | 0.1484744 | 0.1238374 | 0.6202285 | 0.5784556 | 0.611003 | 0.3585713 | 0.3163257 | 0.3515652 |

| Time (min) | soq1 lhcb4 | | | lhcb5 | | | soq1 lhcb5 | | | WT | | |
|---|---|---|---|---|---|---|---|---|---|---|---|---|
| 0 | 0 | 0 | 0 | 0 | 0 | 0 | 0 | 0 | 0 | 0 | 0 | 0 |
| 0.50483 | 0.8965808 | 0.450508 | 0.5831082 | 0.5674618 | 0.7541801 | 0.6405826 | 0.5929266 | 1.131534 | 1.142444 | 0.6355537 | 1.055509 | 1.263699 |
| 1.50967 | 1.266951 | 0.8995498 | 1.01988 | 1.291522 | 1.514334 | 1.426781 | 1.327855 | 1.901792 | 1.958098 | 1.388138 | 1.771968 | 1.940878 |
| 2.5145 | 1.378382 | 1.105968 | 1.150361 | 1.525515 | 1.723601 | 1.702146 | 1.694723 | 2.218389 | 2.258479 | 1.68843 | 2.051156 | 2.177185 |
| 3.51934 | 1.490162 | 1.316732 | 1.298243 | 1.746185 | 1.869696 | 1.869713 | 1.999766 | 2.390795 | 2.436834 | 1.829732 | 2.177484 | 2.245317 |
| 4.52417 | 1.650576 | 1.560741 | 1.54128 | 1.906931 | 1.961075 | 1.974453 | 2.291638 | 2.603052 | 2.670177 | 1.874316 | 2.254222 | 2.300897 |
| 5.52901 | 1.791141 | 1.756055 | 1.765866 | 2.018127 | 2.024179 | 2.051201 | 2.520764 | 2.751314 | 2.8287 | 1.894781 | 2.289593 | 2.332479 |
| 6.53384 | 1.884818 | 1.922593 | 1.959351 | 2.086617 | 2.084392 | 2.102954 | 2.692466 | 2.867426 | 2.954989 | 1.967222 | 2.342154 | 2.388914 |
| 7.53868 | 1.96769 | 2.086918 | 2.133042 | 2.162904 | 2.139783 | 2.161994 | 2.83286 | 2.983339 | 3.082042 | 1.995413 | 2.419417 | 2.430862 |
| 8.54351 | 2.038805 | 2.215333 | 2.288195 | 2.208909 | 2.189422 | 2.179552 | 2.991956 | 3.06598 | 3.179114 | 2.027182 | 2.447705 | 2.437227 |
| 9.54834 | 2.135059 | 2.331572 | 2.402508 | 2.264652 | 2.212198 | 2.21227 | 3.103847 | 3.159651 | 3.275426 | 2.057136 | 2.524152 | 2.488411 |
| 10.5532 | 2.16935 | 2.430739 | 2.505901 | 2.299496 | 2.23818 | 2.263462 | 3.236473 | 3.249259 | 3.381004 | 2.090605 | 2.553446 | 2.540464 |
| 10.8085 | 1.40169 | 1.460149 | 1.548066 | 1.22263 | 1.201368 | 1.139649 | 2.352181 | 2.46894 | 2.584904 | 1.319926 | 1.728854 | 1.69183 |
| 11.1463 | 1.14942 | 1.426583 | 1.479617 | 0.8214874 | 0.8379373 | 0.7714145 | 2.004026 | 1.96847 | 2.102162 | 0.956297 | 1.22385 | 1.176266 |
| 12.1508 | 0.9502186 | 1.36615 | 1.426721 | 0.472469 | 0.4296669 | 0.4571859 | 1.632015 | 1.325985 | 1.436107 | 0.5797145 | 0.7167878 | 0.6539458 |
| 13.1553 | 0.9001029 | 1.319125 | 1.373986 | 0.4171615 | 0.3681246 | 0.3990382 | 1.570205 | 1.216138 | 1.319162 | 0.5325503 | 0.6463999 | 0.5612805 |
| 14.1598 | 0.8715699 | 1.304334 | 1.355456 | 0.3904019 | 0.3414043 | 0.376825 | 1.53886 | 1.166363 | 1.25637 | 0.4802626 | 0.6045197 | 0.5210197 |
| 15.1643 | 0.8458838 | 1.290133 | 1.341705 | 0.3800133 | 0.3313391 | 0.3625391 | 1.507496 | 1.128473 | 1.225173 | 0.4649301 | 0.5850615 | 0.4905306 |
| 16.1688 | 0.81366 | 1.276311 | 1.32126 | 0.3716457 | 0.3223863 | 0.354125 | 1.479497 | 1.084606 | 1.19632 | 0.4501755 | 0.5681302 | 0.468234 |
| 17.1733 | 0.7885646 | 1.272335 | 1.317833 | 0.3679874 | 0.317761 | 0.3474329 | 1.476349 | 1.057495 | 1.160993 | 0.4488063 | 0.5489935 | 0.451812 |
| 18.1778 | 0.755239 | 1.249041 | 1.300754 | 0.3591162 | 0.3073091 | 0.3393754 | 1.452276 | 1.018685 | 1.124581 | 0.4394132 | 0.5235168 | 0.4275225 |
| 19.1823 | 0.7410301 | 1.260605 | 1.294536 | 0.3503586 | 0.3050877 | 0.3301924 | 1.440921 | 0.984429 | 1.094223 | 0.423386 | 0.491839 | 0.3992118 |
| 20.1868 | 0.7345179 | 1.238838 | 1.272975 | 0.3426551 | 0.303446 | 0.3195834 | 1.431948 | 0.9600091 | 1.067578 | 0.4257021 | 0.46698 | 0.3777049 |
| 21.1913 | 0.7254937 | 1.235286 | 1.259582 | 0.3339893 | 0.3004335 | 0.3101873 | 1.407058 | 0.9363948 | 1.044531 | 0.419368 | 0.4443707 | 0.3563268 |

Figure 3 E

NPQ

| | WT | *soq1* | *lcnp* | *NoM* | *soq1 NoM* | *lhcb4* | *soq1 lhcb4* | *lhcb5* | *soq1 lhcb5* | *lhcb6* | *soq1 lhcb6* |
|---|---|---|---|---|---|---|---|---|---|---|---|
| | 6.98412647 | 11.2204917 | 3.56846696 | 7.12388193 | 10.0936992 | 2.80202875 | 10.7837474 | 2.89784946 | 9.77517415 | 5.15322985 | 9.7895844 |
| Rep1 | 6.07908731 | 12.3017004 | 3.76128769 | 4.21807892 | 10.9332299 | 5.19519668 | 10.3817558 | 2.07193248 | 10.0500821 | 2.88173809 | 8.55723534 |
| | 6.34845503 | 12.0776335 | 3.40857881 | 4.50362502 | 9.25188048 | 3.56008492 | 12.7368157 | 2.74819427 | 9.31852747 | 4.02277152 | 9.30965042 |
| | 3.07906588 | 9.42850924 | 2.59209426 | 5.4422967 | 11.4219271 | 2.25559636 | 9.76210418 | 2.04490472 | 11.5052854 | 2.77312386 | 5.68090764 |
| Rep2 | 4.43856826 | 10.9746749 | 2.85872633 | 3.76197237 | 10.7828578 | 2.35932379 | 10.8282564 | 1.91310177 | 9.33859632 | 1.68384295 | 8.22936304 |
| | 5.49952232 | 8.71672402 | 3.2375235 | 5.00056587 | 11.9964157 | 3.03962409 | 10.2488241 | 1.35718952 | 10.8973419 | 3.0459403 | 6.19131363 |
| | 6.77021182 | 12.5947683 | 2.71420268 | 5.64600014 | 9.09542099 | 5.45850326 | 8.66591304 | 4.17708522 | 12.8472518 | 6.05321876 | 9.59187126 |
| Rep3 | 4.44013818 | 11.0337605 | 2.53578546 | 5.74134453 | 6.29388393 | 3.98156285 | 9.3597273 | 5.37232979 | 11.9301011 | 6.08530925 | 10.3930951 |
| | 4.33782571 | 13.24643 | 2.72723083 | 4.74981659 | 7.22467937 | 5.22407987 | 11.0218146 | 4.22548533 | 11.1922145 | 5.12624011 | 8.8267219 |

Figure 3 D

| | WT | | | soq1 | | | lhcb6 | | | soq1 lhcb6-2 | | |
|---|---|---|---|---|---|---|---|---|---|---|---|---|
| 0 | 0 | 0 | 0 | 0 | 0 | 0 | 0 | 0 | 0 | 0 | 0 | 0 |
| 0.50483 | 1.06777 | 1.066259 | 0.8932549 | 0.9264961 | 1.005302 | 1.154751 | 1.097376 | 1.094102 | 1.022738 | 1.023275 | 1.093111 | 0.987899 |
| 1.50967 | 1.611833 | 1.594566 | 1.468587 | 1.60838 | 1.547515 | 1.752377 | 1.347142 | 1.238827 | 1.243098 | 1.352914 | 1.319864 | 1.264603 |
| 2.5145 | 1.690765 | 1.679132 | 1.783383 | 2.039657 | 1.862214 | 1.97494 | 1.192575 | 1.235732 | 1.07636 | 1.331584 | 1.277929 | 1.246946 |
| 3.51934 | 1.80043 | 1.796791 | 1.826445 | 2.196758 | 1.988677 | 2.105398 | 1.245865 | 1.366618 | 1.129924 | 1.49657 | 1.432122 | 1.345757 |
| 4.52417 | 1.850923 | 1.870036 | 1.870568 | 2.318737 | 2.104859 | 2.223819 | 1.448808 | 1.525931 | 1.325846 | 1.723874 | 1.661791 | 1.513009 |
| 5.52901 | 1.91757 | 1.946451 | 1.895005 | 2.401095 | 2.177609 | 2.327986 | 1.595218 | 1.593572 | 1.485874 | 1.934031 | 1.799082 | 1.604178 |
| 6.53384 | 1.950563 | 1.985265 | 1.916076 | 2.497923 | 2.263756 | 2.394724 | 1.689567 | 1.670505 | 1.536942 | 2.069013 | 1.890017 | 1.720651 |
| 7.53868 | 1.971518 | 2.009674 | 1.929542 | 2.577001 | 2.337729 | 2.497624 | 1.701217 | 1.756993 | 1.615183 | 2.099277 | 1.966269 | 1.808814 |
| 8.54351 | 1.997481 | 2.040217 | 1.964319 | 2.650821 | 2.399913 | 2.562749 | 1.783293 | 1.78864 | 1.65835 | 2.24815 | 2.055052 | 1.867914 |
| 9.54834 | 2.037013 | 2.064206 | 1.983197 | 2.713932 | 2.485333 | 2.619678 | 1.891515 | 1.841358 | 1.709368 | 2.296921 | 2.24918 | 2.001918 |
| 10.5532 | 2.061592 | 2.099856 | 2.0155 | 2.766195 | 2.555766 | 2.671075 | 1.947495 | 1.888212 | 1.811764 | 2.443382 | 2.360485 | 2.118852 |
| 10.8085 | 1.1224 | 1.104794 | 1.109283 | 1.689189 | 1.647341 | 1.664609 | 0.9650418 | 0.7083819 | 0.884738 | 1.45271 | 1.222604 | 1.046951 |
| 11.1463 | 0.7763088 | 0.7461641 | 0.7784807 | 1.358611 | 1.302135 | 1.347679 | 0.7262391 | 0.5828716 | 0.6768402 | 1.280557 | 0.981057 | 0.868472 |
| 12.1508 | 0.4561544 | 0.433951 | 0.3857512 | 0.9957214 | 0.9517416 | 1.046136 | 0.5278013 | 0.498101 | 0.4867547 | 1.156821 | 0.835491 | 0.7278119 |
| 13.1553 | 0.4023679 | 0.3851349 | 0.3041942 | 0.896148 | 0.8758316 | 0.9832965 | 0.479526 | 0.4729404 | 0.4369023 | 1.1208 | 0.8053256 | 0.6653565 |
| 14.1598 | 0.3726054 | 0.3528354 | 0.2577711 | 0.8137546 | 0.8109166 | 0.930471 | 0.4602121 | 0.431925 | 0.4038495 | 1.071422 | 0.7672749 | 0.6187866 |
| 15.1643 | 0.3514527 | 0.3271732 | 0.2337172 | 0.7650914 | 0.7553175 | 0.8892339 | 0.4293 | 0.4226908 | 0.3740488 | 1.031726 | 0.7437063 | 0.5951264 |
| 16.1688 | 0.3441161 | 0.3176062 | 0.2297764 | 0.7310191 | 0.7122623 | 0.8609693 | 0.4240438 | 0.4127223 | 0.3692026 | 1.018774 | 0.7174884 | 0.5675073 |
| 17.1733 | 0.3384958 | 0.3125022 | 0.2299742 | 0.7140927 | 0.6869961 | 0.8454857 | 0.4125982 | 0.4109679 | 0.3562152 | 0.97512 | 0.6894713 | 0.5133461 |
| 18.1778 | 0.332547 | 0.304487 | 0.2278157 | 0.7013307 | 0.6694974 | 0.8323864 | 0.4114929 | 0.3896299 | 0.3484862 | 0.97195 | 0.6924344 | 0.5221265 |
| 19.1823 | 0.3257957 | 0.2967868 | 0.225275 | 0.6872754 | 0.6490949 | 0.8158765 | 0.3963851 | 0.3794295 | 0.3334745 | 0.937774 | 0.675668 | 0.5004642 |
| 20.1868 | 0.3149947 | 0.288872 | 0.2277446 | 0.6727546 | 0.637122 | 0.8049725 | 0.3933354 | 0.3867012 | 0.3239401 | 0.892662 | 0.6336532 | 0.4657488 |
| 21.1913 | 0.3069634 | 0.286865 | 0.2282788 | 0.6640512 | 0.6283824 | 0.7830969 | 0.3852603 | 0.3727086 | 0.3295154 | 0.86129 | 0.6293389 | 0.4429923 |

Figure 4 C

| | Raw data | WT | *WT CHL* | *soq1* | *soq1 CHL* | *soq1 lhcb6* | soq1 lhcb6 CHL |
|---|---|---|---|---|---|---|---|
| Rep1 | | WT | *WT CHL* | *soq1* | *soq1 CHL* | *soq1 lhcb6* | soq1 lhcb6 CHL |
| | Green band | 4606 | 4468 | 4446 | 4642 | 5439 | 5045 |
| | Fluorescence | 730.3669 | 603.8694 | 692.3906 | 455.8577 | 692.014 | 463.7902 |
| | Fluorescence/ green band | 0.15856858 | 0.1351543 | 0.15573338 | 0.09820287 | 0.12723184 | 0.09193066 |
| Rep 2 | | WT | *WT CHL* | *soq1* | *soq1 CHL* | *soq1 lhcb6* | soq1 lhcb6 CHL |
| | Green band | 4088 | 3616 | 3779 | 4302 | 4478 | 4574 |
| | Fluorescence | 681.7627 | 564.1541 | 646.1095 | 447.9729 | 672.4935 | 440.8537 |
| | Fluorescence/ green band | 0.1667717 | 0.15601607 | 0.17097367 | 0.10413131 | 0.1501772 | 0.09638253 |
| Rep 3 | | WT | *WT CHL* | *soq1* | *soq1 CHL* | *soq1 lhcb6* | soq1 lhcb6 CHL |
| | Green band | 4068 | 4176 | 3801 | 4278 | 4643 | 4813 |
| | Fluorescence | 734.438 | 616.7712 | 708.0906 | 452.093 | 700.2479 | 482.7437 |
| | Fluorescence/ green band | 0.18054031 | 0.14769425 | 0.18629061 | 0.10567859 | 0.15081798 | 0.10029996 |

| Normalized data to WT | WT | *WT CHL* | *soq1* | *soq1 CHL* | *soq1 lhcb6* | soq1 lhcb6 CHL |
|---|---|---|---|---|---|---|
| Rep1 | 0.94035185 | 0.8014992 | 0.92353836 | 0.58236785 | 0.75451704 | 0.54517211 |
| Rep2 | 0.98899838 | 0.92521477 | 1.01391715 | 0.61752503 | 0.89058881 | 0.57157281 |
| Rep3 | 1.07064977 | 0.87586431 | 1.10475046 | 0.62670078 | 0.89438882 | 0.59480414 |

Figure 4 A, B

Rep1

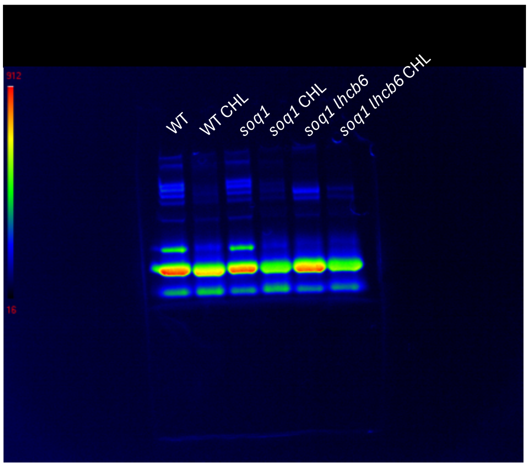


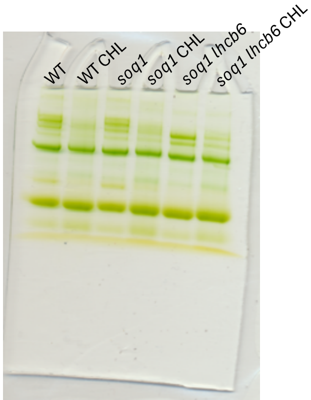


Rep2

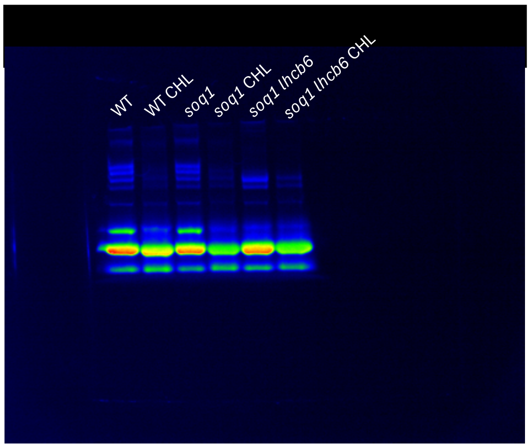


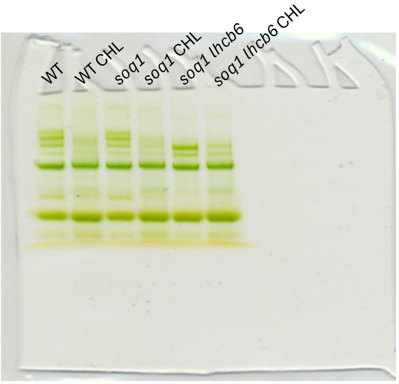


Rep3

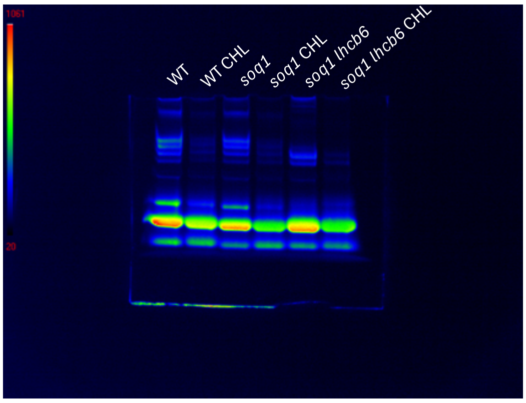


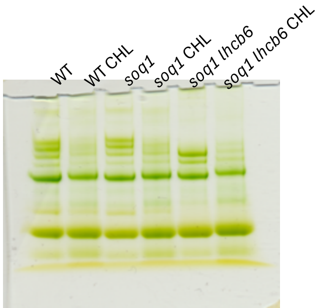

Figure 5

| | Rep1 | Rep2 | Rep3 | Rep1 | Rep2 | Rep3 | Rep1 | Rep2 | Rep3 |
|---|---|---|---|---|---|---|---|---|---|
| Time (H) | *koLHCII* | | | *soq1 koLHCII* | | | *soq1 koLHCII lcnp* | | |
| 0 | 2.6 | 2.8 | 3.2 | 2.7 | 3 | 3.3 | 2.3 | 2.8 | 3.3 |
| 6 | 1.9 | 1.7 | 1.9 | 1.8 | 1.8 | 2.1 | 1.6 | 1.7 | 2.1 |
| 20 | 1.7 | 1.2 | 1.1 | 1.8 | 1.8 | 2 | 0 | 0 | 0 |
| 24 | 1.3 | 0.6 | 0.8 | 1.7 | 1.9 | 2 | 0 | 0 | 0 |
| 30 | 0.8 | 0 | 0.1 | 1.6 | 1.4 | 2.1 | 0 | 0 | 0 |
| 45 | 0.1 | 0 | 0 | 1.2 | 0.3 | 0.4 | 0 | 0 | 0 |

Rep1 Rep2 Rep3

T=oH

T=24H

T=30H

T=45H

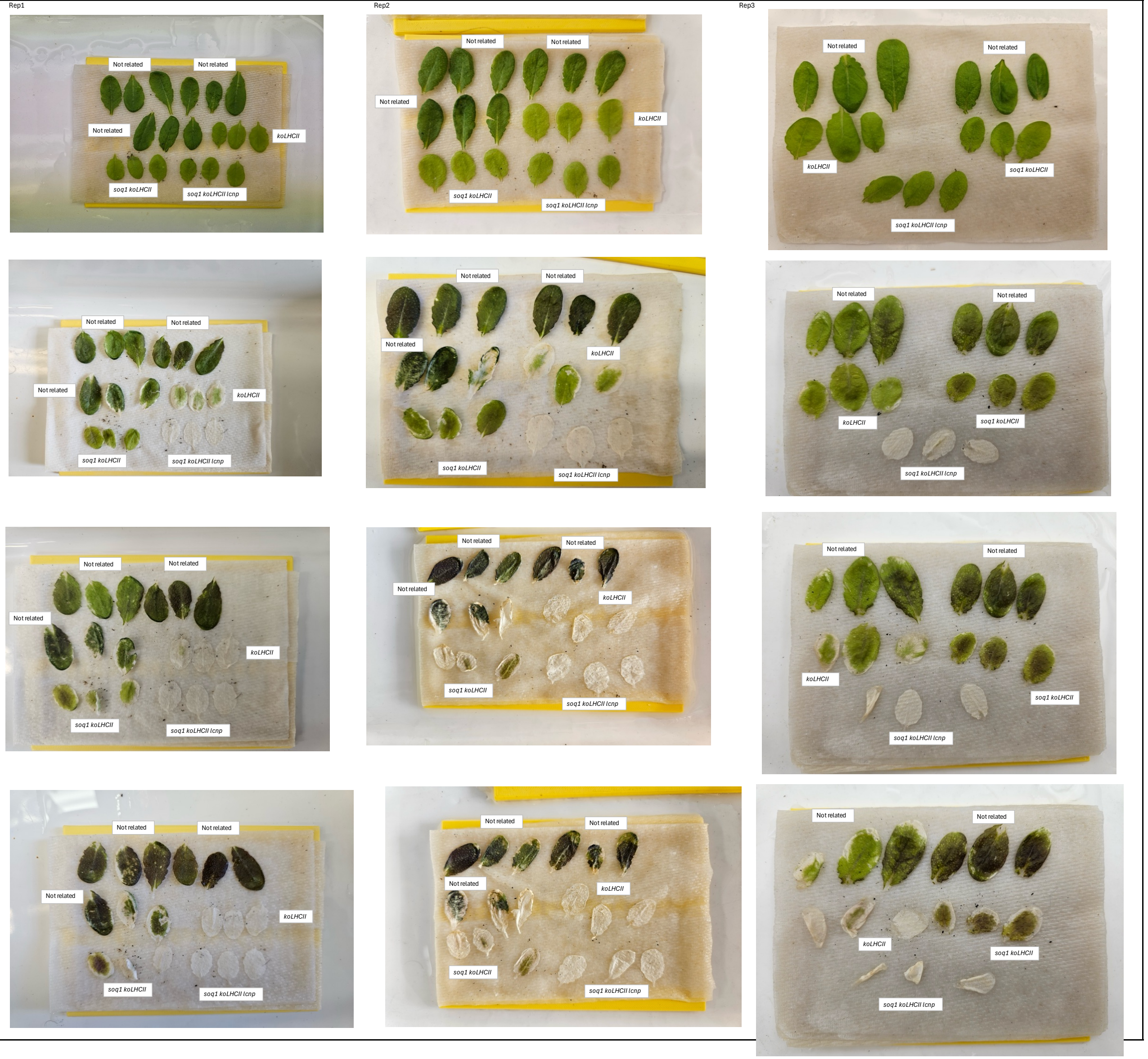

Rep1

| Fo | | | | | | | | | | | | |
|---|---|---|---|---|---|---|---|---|---|---|---|---|
| WT | soq1 | lcnp | lhcb1 | soq1 lhcb1 | soq1 lhcb1 lcnp | koLHCII | soq1 koLHCII | soq1 koLHCII lcnp | soq1 npq4 | npq4 | soq1 npq4 cpsrp43 | soq1 npq4 cpsrp43 lcnp |
| 101.603 | 128.862 | 122.245 | 84.8125 | 84.2693 | 79.6966 | 77.06 | 78.5902 | 74.7878 | 139.684 | 129.511 | 51.042 | 60.2765 |
| 112.072 | 132.911 | 116.961 | 82.2446 | 87.8248 | 76.7137 | 82.9359 | 95.6767 | 79.8248 | 141.165 | 131.289 | 50.3506 | 55.2889 |
| 118.301 | 131.783 | 151.141 | 74.2446 | 86.8372 | 82.66 | 81.8989 | 79.973 | 85.6449 | 140.573 | 116.277 | 49.4617 | 55.2889 |

| Fm | | | | | | | | | | | | |
|---|---|---|---|---|---|---|---|---|---|---|---|---|
| WT | soq1 | lcnp | lhcb1 | soq1 lhcb1 | soq1 lhcb1 lcnp | koLHCII | soq1 koLHCII | soq1 koLHCII lcnp | soq1 npq4 | npq4 | soq1 npq4 cpsrp43 | soq1 npq4 cpsrp43 lcnp |
| 554.636 | 646.586 | 631.673 | 466.34 | 476.068 | 405.813 | 306.108 | 321.796 | 299.969 | 720.237 | 669.916 | 368.434 | 391.2 |
| 600.611 | 695.426 | 614.34 | 443.772 | 490.241 | 398.29 | 315.821 | 350.982 | 307.969 | 738.459 | 664.829 | 330.064 | 367.842 |
| 605.718 | 694.656 | 653.422 | 411.08 | 489.895 | 413.558 | 328.364 | 312.265 | 343.842 | 701.471 | 604.187 | 347.545 | 382.311 |

| Fv/Fm | | | | | | | | | | | | |
|---|---|---|---|---|---|---|---|---|---|---|---|---|
| WT | soq1 | lcnp | lhcb1 | soq1 lhcb1 | soq1 lhcb1 lcnp | koLHCII | soq1 koLHCII | soq1 koLHCII lcnp | soq1 npq4 | npq4 | soq1 npq4 cpsrp43 | soq1 npq4 cpsrp43 lcnp |
| 0.81681 | 0.8007 | 0.80647 | 0.81813 | 0.82299 | 0.80361 | 0.74826 | 0.75578 | 0.75068 | 0.80606 | 0.80668 | 0.86146 | 0.84592 |
| 0.8134 | 0.80888 | 0.80962 | 0.81467 | 0.82085 | 0.80739 | 0.7374 | 0.7274 | 0.7408 | 0.80884 | 0.80252 | 0.84745 | 0.84969 |
| 0.80469 | 0.81029 | 0.80119 | 0.81939 | 0.82274 | 0.80012 | 0.75059 | 0.74389 | 0.75092 | 0.7996 | 0.80755 | 0.85768 | 0.85538 |

Rep2

| Fo | | | | | | | | | | | | |
|---|---|---|---|---|---|---|---|---|---|---|---|---|
| WT | soq1 | lcnp | lhcb1 | soq1 lhcb1 | soq1 lhcb1 lcnp | koLHCII | soq1 koLHCII | soq1 koLHCII lcnp | soq1 npq4 | npq4 | soq1 npq4 cpsrp43 | soq1 npq4 cpsrp43 lcnp |
| 118.091 | 136.56 | 131.572 | 85.8436 | 88.609 | 87.4732 | 97.1579 | 95.7695 | 65.0535 | 129.412 | 131.486 | 64.1778 | 53.116 |
| 129.3 | 132.658 | 122.584 | 89.5967 | 89.0041 | 72.8559 | 85.1522 | 87.6214 | 91.2757 | 157.659 | 140.77 | 61.8568 | 54.5975 |
| 99.9802 | 142.548 | 142.943 | 85.251 | 82.4856 | 82.6831 | 84.5596 | 109.449 | 94.8806 | 159.289 | 165.956 | 61.6593 | 60.1778 |

| Fm | | | | | | | | | | | | |
|---|---|---|---|---|---|---|---|---|---|---|---|---|
| WT | soq1 | lcnp | lhcb1 | soq1 lhcb1 | soq1 lhcb1 lcnp | koLHCII | soq1 koLHCII | soq1 koLHCII lcnp | soq1 npq4 | npq4 | soq1 npq4 cpsrp43 | soq1 npq4 cpsrp43 lcnp |
| 658.403 | 682.847 | 664.181 | 452.872 | 476.131 | 466.008 | 352.059 | 353.366 | 267.045 | 686.149 | 718.692 | 411.976 | 397.063 |
| 573.309 | 678.107 | 649.317 | 461.81 | 485.81 | 396.576 | 323.884 | 333.563 | 341.563 | 785.853 | 725.112 | 413.26 | 398.445 |
| 691.532 | 782.198 | 713.408 | 445.613 | 457.86 | 438.354 | 316.378 | 388.872 | 361.465 | 781.507 | 709.458 | 387.877 | 436.519 |

| Fv/Fm | | | | | | | | | | | | |
|---|---|---|---|---|---|---|---|---|---|---|---|---|
| WT | soq1 | lcnp | lhcb1 | soq1 lhcb1 | soq1 lhcb1 lcnp | koLHCII | soq1 koLHCII | soq1 koLHCII lcnp | soq1 npq4 | npq4 | soq1 npq4 cpsrp43 | soq1 npq4 cpsrp43 lcnp |
| 0.80795 | 0.80001 | 0.8019 | 0.81045 | 0.8139 | 0.81229 | 0.72403 | 0.72898 | 0.7564 | 0.81139 | 0.81705 | 0.84422 | 0.86623 |
| 0.80362 | 0.80437 | 0.81121 | 0.80599 | 0.81679 | 0.81629 | 0.73709 | 0.73732 | 0.73277 | 0.79938 | 0.80586 | 0.85032 | 0.86297 |
| 0.82561 | 0.81776 | 0.79963 | 0.80869 | 0.81985 | 0.81138 | 0.73273 | 0.71855 | 0.73751 | 0.79618 | 0.76608 | 0.84103 | 0.86214 |

Rep3

| Fo | | | | | | | | | | | | |
|---|---|---|---|---|---|---|---|---|---|---|---|---|
| WT | soq1 | lcnp | lhcb1 | soq1 lhcb1 | soq1 lhcb1 lcnp | koLHCII | soq1 koLHCII | soq1 koLHCII lcnp | soq1 npq4 | npq4 | soq1 npq4 cpsrp43 | soq1 npq4 cpsrp43 lcnp |
| 119.6617 | 126.7361 | 149.2485 | 100.984 | 98.53777 | 95.8511 | 91.63054 | 80.53484 | 75.3609 | 146.147 | 136.428 | 60.34909 | 58.27787 |
| 122.9675 | 131.5295 | 142.0419 | 91.36421 | 95.1328 | 41.67506 | 84.16569 | 79.66682 | 78.00553 | 152.8578 | 136.7256 | 62.25973 | 62.84023 |
| 121.2155 | 112.3229 | 133.513 | 113.7113 | 99.16586 | 94.55405 | 98.65095 | 84.33803 | 94.56751 | 129.8165 | 124.1636 | 60.58173 | 70.28694 |

| Fm | | | | | | | | | | | | |
|---|---|---|---|---|---|---|---|---|---|---|---|---|
| WT | soq1 | lcnp | lhcb1 | soq1 lhcb1 | soq1 lhcb1 lcnp | koLHCII | soq1 koLHCII | soq1 koLHCII lcnp | soq1 npq4 | npq4 | soq1 npq4 cpsrp43 | soq1 npq4 cpsrp43 lcnp |
| 640.0319 | 678.412 | 771.8666 | 519.3046 | 516.0649 | 455.726 | 334.3276 | 289.8188 | 296.1395 | 736.3403 | 691.4478 | 364.7477 | 355.6696 |
| 686.5443 | 671.3377 | 757.0567 | 459.9327 | 489.6848 | 195.4725 | 313.4115 | 290.1825 | 313.8254 | 749.0015 | 713.4313 | 386.9439 | 364.2591 |
| 663.4368 | 571.6352 | 711.4368 | 555.4699 | 507.4699 | 471.3087 | 355.5975 | 302.2006 | 349.3957 | 648.5718 | 687.1503 | 353.2387 | 404.631 |

| Fv/Fm | | | | | | | | | | | | |
|---|---|---|---|---|---|---|---|---|---|---|---|---|
| WT | soq1 | lcnp | lhcb1 | soq1 lhcb1 | soq1 lhcb1 lcnp | koLHCII | soq1 koLHCII | soq1 koLHCII lcnp | soq1 npq4 | npq4 | soq1 npq4 cpsrp43 | soq1 npq4 cpsrp43 lcnp |
| 0.81303791 | 0.81318712 | 0.80663952 | 0.80553995 | 0.80905935 | 0.789673839 | 0.72592589 | 0.72212003 | 0.745522296 | 0.8026923 | 0.80152248 | 0.83454566 | 0.836146047 |
| 0.8208892 | 0.80407848 | 0.81237614 | 0.80135309 | 0.80572646 | 0.786798348 | 0.73145309 | 0.7254596 | 0.751436531 | 0.8083549 | 0.7959179 | 0.83909882 | 0.827484804 |
| 0.81729156 | 0.80350598 | 0.8123333 | 0.7952881 | 0.8045877 | 0.799379791 | 0.72257693 | 0.72092038 | 0.7293398 | 0.8193065 | 0.79984252 | 0.82849634 | 0.826293734 |

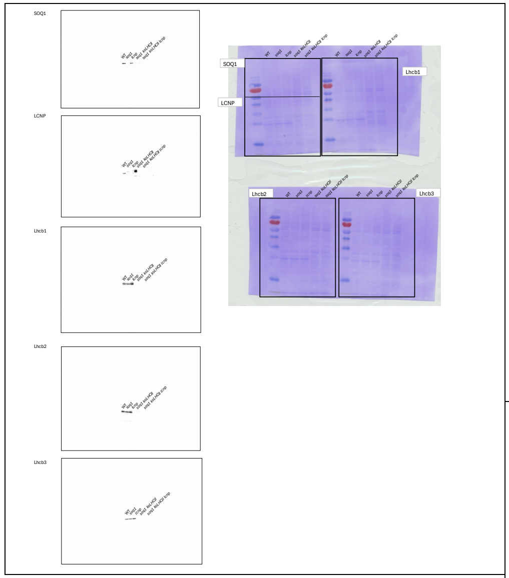


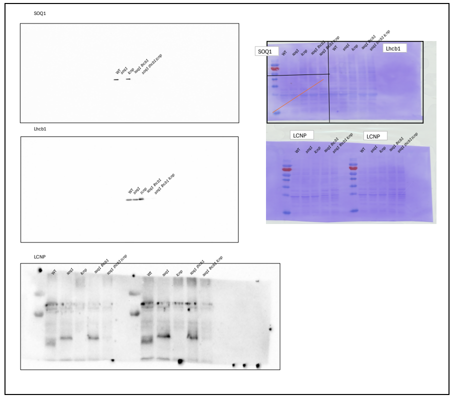


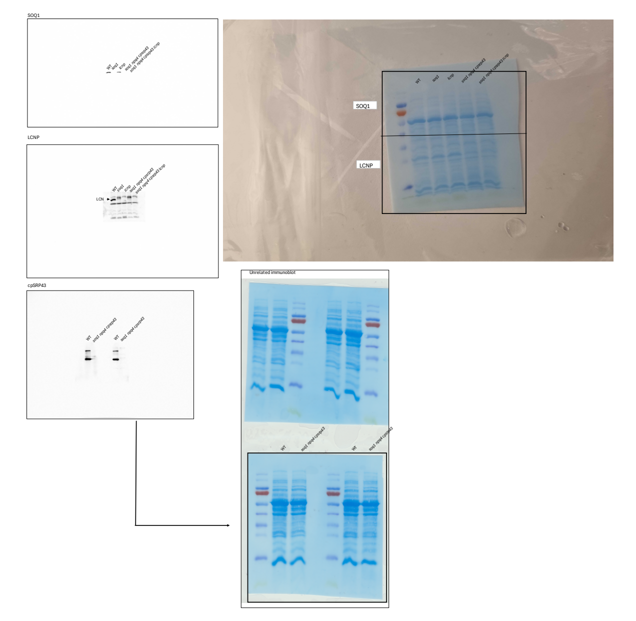


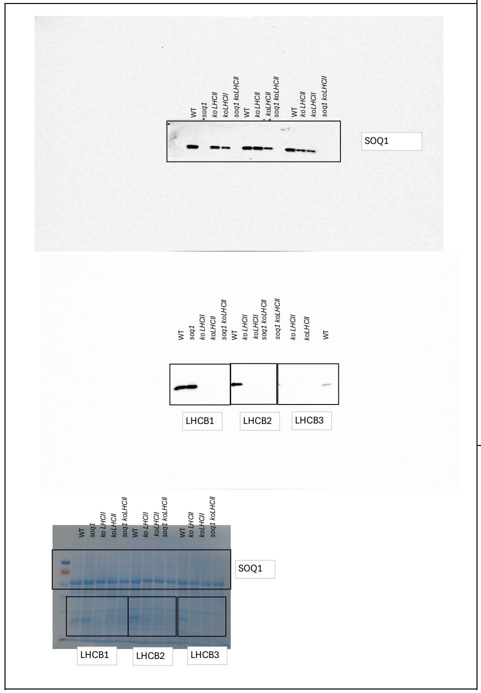

S2B See Figure 2 data for gels

| | Band | Rep 1 | Rep 2 | Rep 3 |
|---|---|---|---|---|
| | | Green band intensity | | |
| WT | Trimers | 3851 | 2786 | |
| WT | Monomers | 630 | 408 | |
| E40 | Trimers | 1980 | 1456 | 1800 |
| E40 | Monomers | 570 | 526 | 453 |

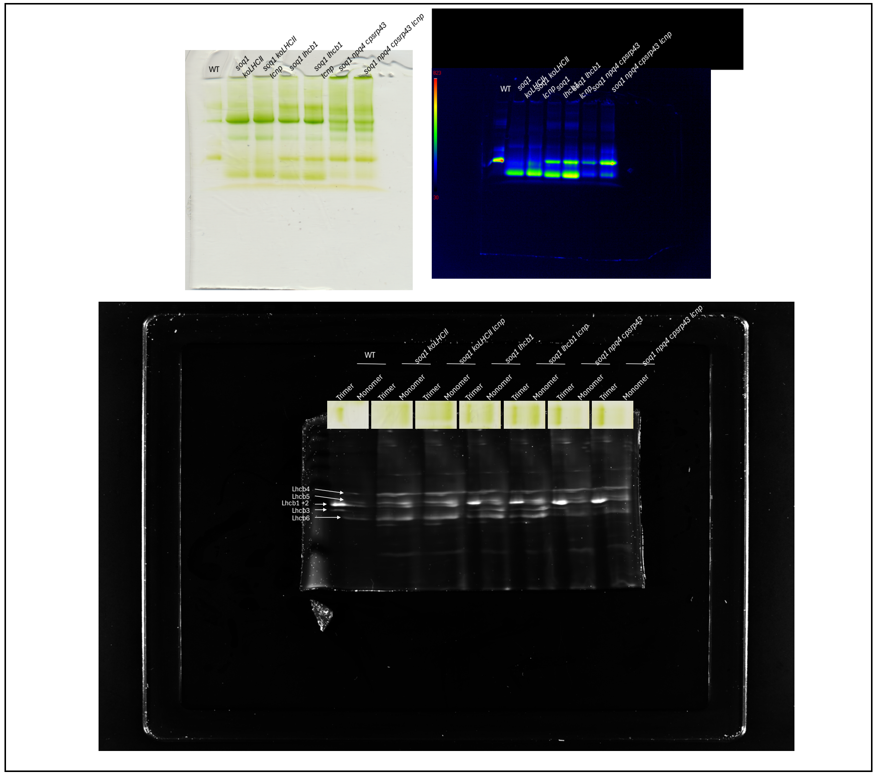


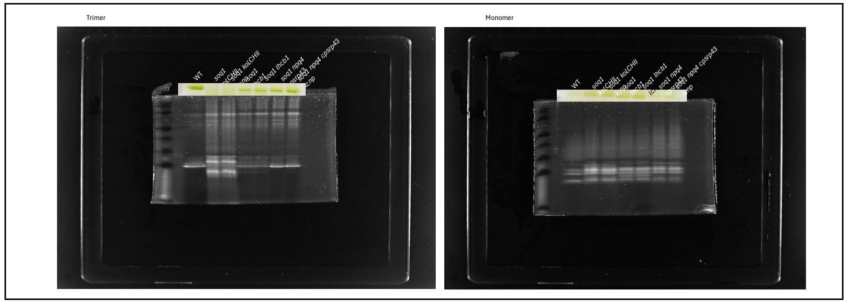


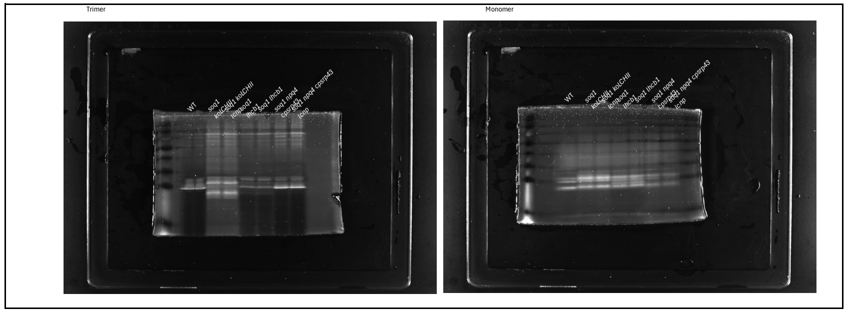


Figure S3 C

Trimer

| | | soq1 lhcb1 | oq1 lhcb1 lcn | E40 | E40 lcnp |
|---|---|---|---|---|---|
| Biological rep 1 | Band | 1160 | 957 | 1959 | 1770 |
| | fluo | 375.05 | 442.37 | 283.45 | 368.06 |
| | Fluorence/green band intensity | 0.32 | 0.46 | 0.14 | 0.21 |
| Biological rep 2 | Band | 1157 | 944 | 2322 | 2184 |
| | fluo | 382.9078 | 455.2534 | 377.9633 | 509.2353 |
| | Fluorence/green band intensity | 0.33 | 0.48 | 0.16 | 0.23 |
| Biological rep 3 | Band | 1245 | 1066 | 2147 | 2158 |
| | fluo | 366.0556 | 444.2 | 379.3 | 439.5909 |
| | Fluorence/green band intensity | 0.29 | 0.42 | 0.18 | 0.20 |

Monomer

| | | soq1 ko | soq1 ko lcnp | soq1 lhcb1 | oq1 lhcb1 lcn | E40 | E40 lcnp |
|---|---|---|---|---|---|---|---|
| Biological rep 1 | Band | 945 | 966 | 1403 | 1090 | 364 | 331 |
| | fluo | 329.18 | 397.31 | 427.33 | 456.67 | 223.93 | 288.49 |
| | Fluorence/green band intensity | 0.35 | 0.41 | 0.30 | 0.42 | 0.62 | 0.87 |
| Biological rep 2 | Band | 1251 | 1035 | 1601 | 1180 | 505 | 375 |
| | fluo | 389.3275 | 439.5497 | 436.1424 | 497.772 | 259.8954 | 325.13 |
| | Fluorence/green band intensity | 0.31 | 0.42 | 0.27 | 0.42 | 0.51 | 0.87 |
| Biological rep 3 | Band | 1093 | 894 | 1216 | 1077 | 494 | 365 |
| | fluo | 406.5494 | 444.6728 | 450.4506 | 501.0679 | 261.3642 | 285.7592 |
| | Fluorence/green band intensity | 0.37 | 0.50 | 0.37 | 0.47 | 0.53 | 0.78 |

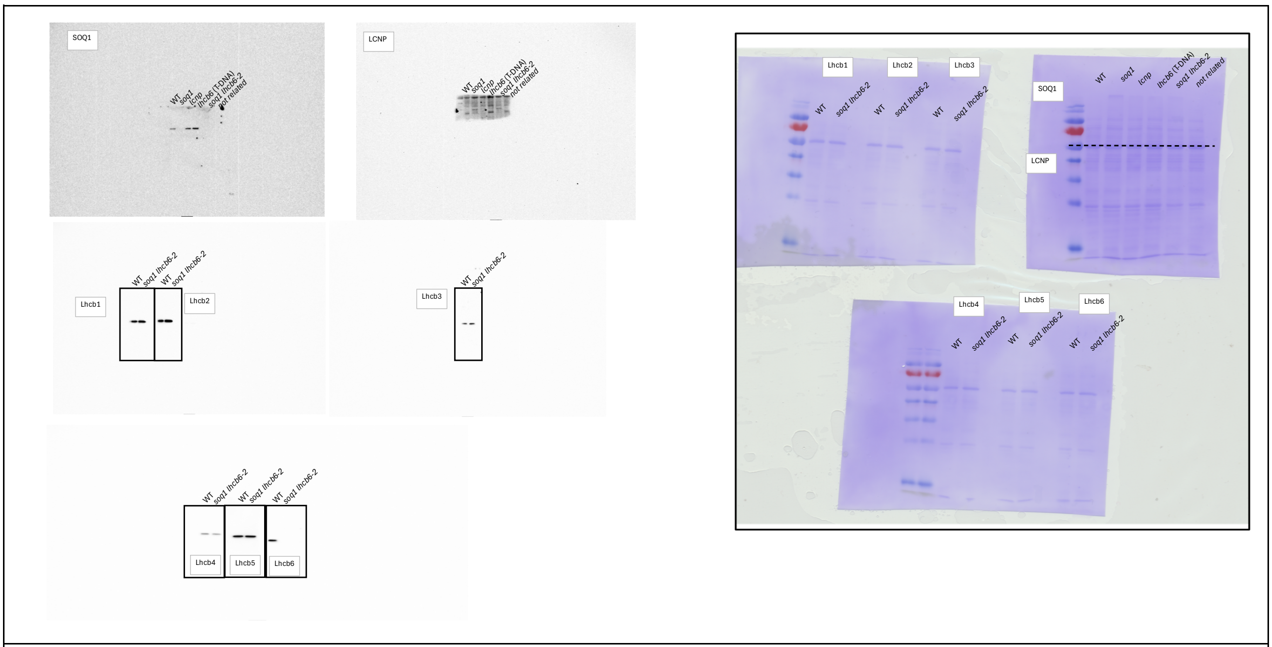
SOQ1
WT
soq1
lcnp
lhcb6 (T-DNA)
soq1 lhcb6-2
not related
LCNP
Lhcb1
Lhcb2
Lhcb3
Lhcb4
Lhcb5
Lhcb6
WT
soq1 lhcb6-2

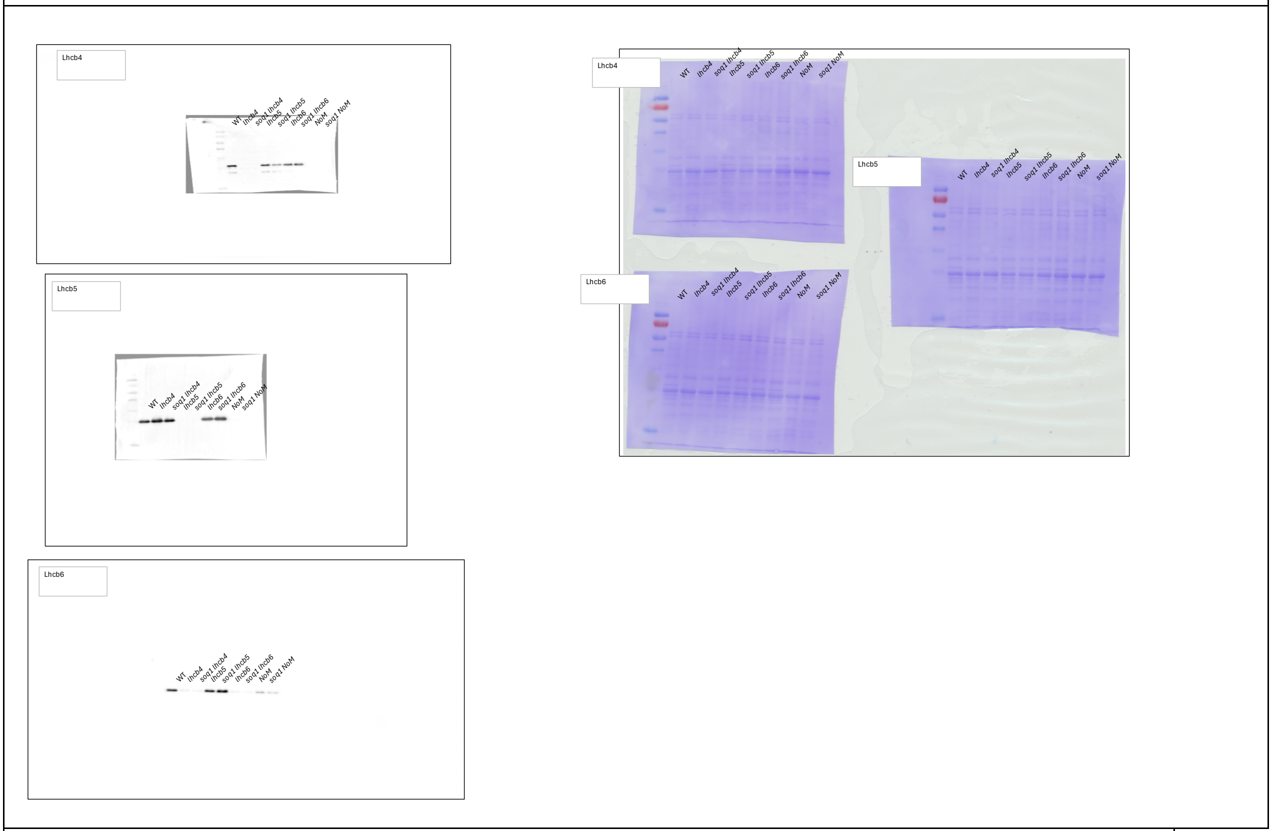
Lhcb4
Lhcb5
Lhcb6
WT
lhcb4
soq1 lhcb4
lhcb5
soq1 lhcb5
lhcb6
soq1 lhcb6
NoM
soq1 NoM

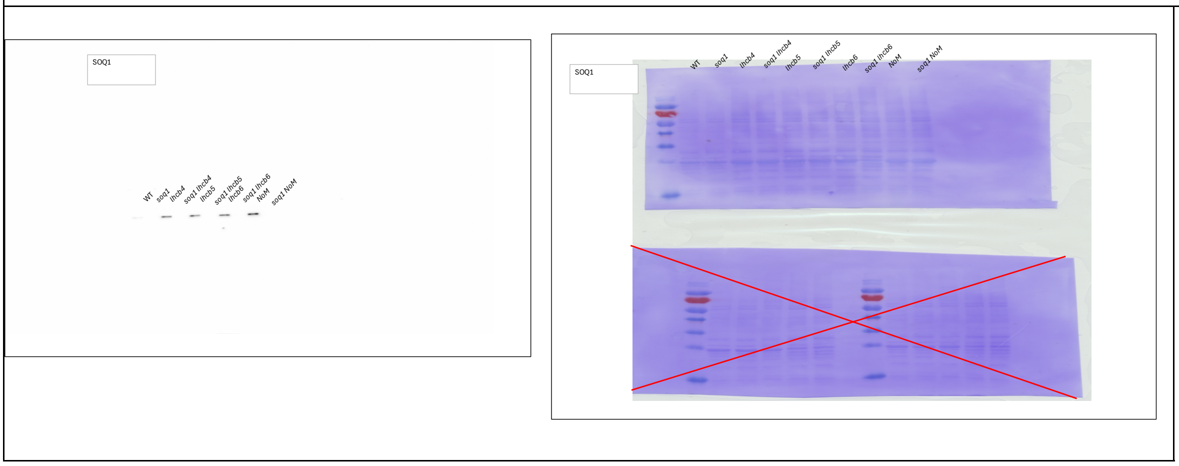
SOQ1
WT
soq1
lhcb4
soq1 lhcb4
lhcb5
soq1 lhcb5
lhcb6
soq1 lhcb6
NoM
soq1 NoM

Figure S5 A, B, C, D

**Fo**

| | WT | soq1 | NoM | soq1 NoM | lhcb4 | soq1 lhcb4 | lhcb5 | soq1 lhcb5 |
|---|---|---|---|---|---|---|---|---|
| | 101.603 | 128.862 | 334.844 | 359.19 | 115.338 | 151.486 | 136.523 | 119.388 |
| Rep1 | 112.072 | 132.911 | 316.523 | 345.067 | 143.388 | 161.561 | 126.696 | 123.931 |
| | 118.301 | 131.783 | 370.449 | 357.314 | 113.314 | 156.326 | 119.635 | 120.277 |
| | 118.091 | 136.56 | 298.943 | 369.511 | 97.9062 | 161.264 | 121.659 | 131.536 |
| Rep2 | 129.3 | 132.658 | 354.203 | 405.017 | 117.017 | 156.474 | 116.474 | 132.721 |
| | 99.9802 | 142.548 | 373.067 | 429.215 | 131.24 | 189.906 | 130.795 | 105.906 |
| | 119.6617 | 126.7361 | 369.7119 | 365.3581 | 156.5146 | 130.637 | 119.0633 | 134.4012 |
| Rep3 | 122.9675 | 131.5295 | 366.6098 | 394.0474 | 154.9635 | 151.8978 | 129.5395 | 133.8479 |
| | 121.2155 | 112.3229 | 329.6393 | 339.0996 | 126.8728 | 167.0179 | 126.3558 | 114.7277 |

**Fm**

| | WT | soq1 | NoM | soq1 NoM | lhcb4 | soq1 lhcb4 | lhcb5 | soq1 lhcb5 |
|---|---|---|---|---|---|---|---|---|
| | 554.636 | 646.586 | 649.718 | 561.175 | 468.187 | 605.125 | 648.73 | 571.298 |
| Rep1 | 600.611 | 695.426 | 588.533 | 583.101 | 597.767 | 616.73 | 607.249 | 562.459 |
| | 605.718 | 694.656 | 663.94 | 606.459 | 492.681 | 565.57 | 592.829 | 539.693 |
| | 658.403 | 682.847 | 600.272 | 672.717 | 446.445 | 626.149 | 617.211 | 625.507 |
| Rep2 | 573.309 | 678.107 | 658.742 | 682.198 | 522.495 | 620.569 | 559.137 | 635.038 |
| | 691.532 | 782.198 | 679.087 | 787.137 | 570.445 | 713.359 | 578.939 | 508.668 |
| | 640.0319 | 678.412 | 701.6813 | 629.8809 | 643.967 | 519.9307 | 588.357 | 609.7629 |
| Rep3 | 686.5443 | 671.3377 | 717.3547 | 622.933 | 622.0441 | 560.1666 | 616.6926 | 606.2074 |
| | 663.4368 | 571.6352 | 658.3888 | 569.2188 | 529.5543 | 604.6835 | 592.2028 | 540.4023 |

**Fv/Fm**

| | WT | soq1 | NoM | soq1 NoM | lhcb4 | soq1 lhcb4 | lhcb5 | soq1 lhcb5 |
|---|---|---|---|---|---|---|---|---|
| | 0.817 | 0.801 | 0.485 | 0.360 | 0.754 | 0.750 | 0.790 | 0.791 |
| Rep1 | 0.813 | 0.809 | 0.462 | 0.408 | 0.760 | 0.738 | 0.791 | 0.780 |
| | 0.805 | 0.810 | 0.442 | 0.411 | 0.770 | 0.724 | 0.798 | 0.777 |
| | 0.808 | 0.800 | 0.502 | 0.451 | 0.781 | 0.742 | 0.803 | 0.790 |
| Rep2 | 0.804 | 0.804 | 0.462 | 0.406 | 0.776 | 0.748 | 0.792 | 0.791 |
| | 0.826 | 0.818 | 0.451 | 0.455 | 0.770 | 0.734 | 0.774 | 0.792 |
| | 0.813 | 0.813 | 0.473 | 0.420 | 0.757 | 0.749 | 0.798 | 0.780 |
| Rep3 | 0.821 | 0.804 | 0.489 | 0.367 | 0.751 | 0.729 | 0.790 | 0.779 |
| | 0.817 | 0.804 | 0.499 | 0.404 | 0.760 | 0.724 | 0.787 | 0.788 |

| | Fo | | | | Fm | | | | Fv/Fm | | | |
|---|---|---|---|---|---|---|---|---|---|---|---|---|
| | WT | *soq1* | *lhcb6* | *soq1 lhcb6-2* | WT | *soq1* | *lhcb6* | *soq1 lhcb6-2* | WT | *soq1* | *lhcb6* | *soq1 lhcb6-2* |
| | 68.36108 | 78.90653 | 84.72472 | 89.58423 | 416.7286 | 440.398 | 378.7121 | 340.5302 | 0.84 | 0.82 | 0.78 | 0.74 |
| Rep1 | 72.19579 | 75.10488 | 83.83215 | 93.41894 | 442.5468 | 416.4311 | 375.7369 | 356.96 | 0.84 | 0.82 | 0.78 | 0.74 |
| | 63.4024 | 81.02224 | 85.81563 | 94.9396 | 392.8278 | 441.6542 | 377.6542 | 362.3815 | 0.84 | 0.82 | 0.77 | 0.74 |

Figure S5 E, F

Before cold HL

**Fo**

| | WT | soq1 | lcnp | NoM | soq1 NoM | lhcb4 | soq1 lhcb4 | lhcb5 | soq1 lhcb5 | lhcb6 | soq1 lhcb6 |
|---|---|---|---|---|---|---|---|---|---|---|---|
| | 116.9628 | 141.2043 | 152.8053 | 332.0329 | 449.5914 | 102.7747 | 147.8425 | 108.1327 | 102.1463 | 148.0796 | 172.8325 |
| Rep1 | 141.0592 | 153.5853 | 149.5127 | 315.9795 | 347.8398 | 130.3549 | 170.754 | 101.983 | 112.5771 | 154.6825 | 160.4969 |
| | 118.2732 | 126.5726 | 114.763 | 448.949 | 492.1054 | 134.0283 | 175.9512 | 98.30952 | 128.9762 | 164.4256 | 166.6465 |
| | 137.2628 | 162.8138 | 135.2492 | 313.55 | 337.3323 | 128.6611 | 148.2529 | 121.0821 | 118.0126 | 159.8841 | 170.2152 |
| Rep2 | 137.4805 | 187.9476 | 131.6573 | 350.9821 | 334.2938 | 143.8879 | 160.3981 | 120.8697 | 129.3414 | 159.0224 | 166.2514 |
| | 100.3709 | 120.21 | 121.8675 | 352.6611 | 307.1555 | 180.126 | 153.4956 | 107.0194 | 122.6838 | 197.3988 | 156.2923 |
| | 125.0173 | 133.235 | 110.8314 | 369.1043 | 402.2556 | 155.6894 | 117.4762 | 123.0998 | 124.2336 | 167.3663 | 148.8176 |
| Rep3 | 129.2713 | 126.3507 | 108.8473 | 388.5057 | 436.7872 | 153.0953 | 146.5737 | 125.1225 | 141.7483 | 154.9944 | 162.3323 |
| | 125.4164 | 121.8427 | 113.2804 | 414.8821 | 387.5483 | 131.4717 | 184.3969 | 135.6259 | 128.6145 | 162.8946 | 147.584 |

**Fm**

| | WT | soq1 | lcnp | NoM | soq1 NoM | lhcb4 | soq1 lhcb4 | lhcb5 | soq1 lhcb5 | lhcb6 | soq1 lhcb6 |
|---|---|---|---|---|---|---|---|---|---|---|---|
| | 553.6844 | 622.8627 | 669.8469 | 671.0597 | 601.4591 | 460.1968 | 586.0886 | 513.3 | 473.9803 | 486.9172 | 525.7471 |
| Rep1 | 596.1235 | 667.7153 | 665.5566 | 634.7099 | 543.0017 | 567.9213 | 589.4452 | 466.9236 | 502.9236 | 478.2476 | 508.5317 |
| | 603.6492 | 617.2456 | 611.0687 | 688.9735 | 629.4996 | 549.6357 | 658.6152 | 453.545 | 553.9713 | 527.1022 | 488.187 |
| | 551.0818 | 642.2655 | 624.0161 | 672.7684 | 594.92 | 572.6161 | 563.7908 | 559.6136 | 527.2012 | 500.5694 | 510.0569 |
| Rep2 | 615.4447 | 647.1725 | 614.7553 | 706.3486 | 561.2511 | 626.2055 | 604.4983 | 569.0788 | 539.8815 | 492.6964 | 508.6601 |
| | 529.496 | 595.9227 | 630.5935 | 697.756 | 517.4506 | 676.8067 | 602.4031 | 503.7545 | 516.8611 | 558.8188 | 441.7757 |
| | 676.7567 | 659.6047 | 596.8564 | 817.7099 | 739.6179 | 679.0522 | 455.6147 | 598.9344 | 567.2427 | 590.2484 | 536.008 |
| Rep3 | 676.6841 | 631.1421 | 587.0751 | 833.1203 | 742.2001 | 666.5625 | 555.0342 | 617.5103 | 635.0795 | 563.5001 | 569.3051 |
| | 669.192 | 599.4959 | 587.895 | 809.592 | 649.075 | 545.1656 | 696.2042 | 637.0478 | 559.3788 | 593.4683 | 502.8651 |

**Fv/Fm**

| | WT | soq1 | lcnp | NoM | soq1 NoM | lhcb4 | soq1 lhcb4 | lhcb5 | soq1 lhcb5 | lhcb6 | soq1 lhcb6 |
|---|---|---|---|---|---|---|---|---|---|---|---|
| | 0.78875547 | 0.77329787 | 0.77188026 | 0.50521109 | 0.2524988 | 0.77667229 | 0.74774718 | 0.7893382 | 0.78449252 | 0.69588341 | 0.67126305 |
| Rep1 | 0.76337252 | 0.76998385 | 0.7753569 | 0.50216705 | 0.35941306 | 0.77047013 | 0.71031404 | 0.78158525 | 0.77615467 | 0.67656398 | 0.68439155 |
| | 0.80406965 | 0.79493965 | 0.81219297 | 0.34837987 | 0.21825939 | 0.75615067 | 0.73284674 | 0.78324197 | 0.76717891 | 0.68805746 | 0.65864208 |
| | 0.75092119 | 0.74650079 | 0.78326008 | 0.53394066 | 0.43297872 | 0.77531002 | 0.73704271 | 0.78363267 | 0.77615263 | 0.68059554 | 0.66628194 |
| Rep2 | 0.776616 | 0.70958655 | 0.78583788 | 0.50310357 | 0.40437747 | 0.77022256 | 0.73465914 | 0.78760463 | 0.76042632 | 0.67724059 | 0.67315817 |
| | 0.81044068 | 0.79827921 | 0.80674159 | 0.49457819 | 0.40640614 | 0.73385902 | 0.74519454 | 0.78755644 | 0.76263681 | 0.64675705 | 0.64621798 |
| | 0.81526995 | 0.79800781 | 0.8143081 | 0.54861217 | 0.45613052 | 0.77072543 | 0.74215889 | 0.79446864 | 0.78098687 | 0.71644769 | 0.72235937 |
| Rep3 | 0.80896359 | 0.79980626 | 0.81459391 | 0.53367395 | 0.41149671 | 0.77032116 | 0.73591952 | 0.79737585 | 0.77680227 | 0.72494344 | 0.71485887 |
| | 0.81258533 | 0.79675808 | 0.80731185 | 0.48754175 | 0.40292216 | 0.7588408 | 0.73513963 | 0.78710248 | 0.7700762 | 0.72552098 | 0.70651374 |

After cold HL

**Fo**

| | WT | soq1 | lcnp | NoM | soq1 NoM | lhcb4 | soq1 lhcb4 | lhcb5 | soq1 lhcb5 | lhcb6 | soq1 lhcb6 |
|---|---|---|---|---|---|---|---|---|---|---|---|
| | 58.87123 | 41.99948 | 135.5686 | 77.90666 | 50.90115 | 87.96336 | 39.3389 | 86.7047 | 36.44909 | 71.95866 | 45.54273 |
| Rep1 | 72.38818 | 41.15594 | 126.8067 | 106.9956 | 41.80666 | 68.56098 | 43.67256 | 88.01778 | 35.21247 | 106.1265 | 47.34188 |
| | 67.35564 | 40.86131 | 125.9951 | 109.6246 | 58.87333 | 102.8205 | 43.56989 | 83.53705 | 42.30451 | 92.57919 | 40.35568 |
| | 76.13 | 43.97478 | 113.9579 | 93.05025 | 42.69727 | 108.9303 | 41.82417 | 105.1843 | 34.3342 | 99.89893 | 56.74818 |
| Rep2 | 82.92135 | 42.40311 | 115.137 | 134.0379 | 40.46083 | 122.2183 | 40.80779 | 105.9462 | 39.37909 | 127.1915 | 45.79164 |
| | 63.12432 | 46.59329 | 127.0528 | 104.4576 | 34.33154 | 127.0528 | 40.86668 | 109.8192 | 36.29979 | 122.5837 | 47.78503 |
| | 69.29482 | 38.40287 | 127.6591 | 123.3011 | 74.65778 | 91.66458 | 43.69179 | 82.99337 | 36.94378 | 82.51721 | 49.67367 |
| Rep3 | 93.46413 | 47.30857 | 121.7725 | 122.7072 | 102.8553 | 110.4109 | 51.07501 | 78.44916 | 46.15273 | 78.45372 | 48.82257 |
| | 97.78865 | 35.78159 | 121.2464 | 140.4185 | 79.36101 | 86.36753 | 54.39474 | 90.041 | 41.14046 | 93.49226 | 49.92779 |

**Fm**

| | WT | soq1 | lcnp | NoM | soq1 NoM | lhcb4 | soq1 lhcb4 | lhcb5 | soq1 lhcb5 | lhcb6 | soq1 lhcb6 |
|---|---|---|---|---|---|---|---|---|---|---|---|
| | 69.34815 | 50.96871 | 146.624 | 82.60333 | 54.21628 | 121.0398 | 49.73703 | 131.688 | 43.98818 | 79.13197 | 48.72728 |
| Rep1 | 84.20909 | 50.19774 | 139.785 | 121.6367 | 45.50333 | 91.67123 | 51.7886 | 151.9967 | 45.51311 | 123.2045 | 53.20908 |
| | 82.14641 | 47.19857 | 138.609 | 125.1854 | 61.40333 | 120.5319 | 47.94526 | 121.0036 | 53.68705 | 104.9425 | 47.35243 |
| | 135.1 | 61.58747 | 173.7193 | 104.4299 | 47.89273 | 175.8867 | 52.38667 | 183.7869 | 42.15827 | 132.6671 | 76.34545 |
| Rep2 | 113.163 | 54.0451 | 159.3156 | 148.3311 | 47.63285 | 186.4082 | 51.10629 | 195.3515 | 52.22 | 183.5787 | 55.11324 |
| | 81.46691 | 61.32959 | 148.8118 | 116.2817 | 39.81487 | 167.542 | 53.55254 | 213.7098 | 43.44341 | 138.1184 | 61.43185 |
| | 87.0963 | 48.51901 | 160.6957 | 123.0379 | 73.26271 | 105.1408 | 47.13623 | 115.6895 | 40.96428 | 83.68497 | 50.6056 |
| Rep3 | 124.3873 | 52.44762 | 166.0381 | 123.5837 | 101.7565 | 133.8059 | 53.57614 | 96.90495 | 49.11636 | 79.53077 | 49.96931 |
| | 125.3679 | 42.08043 | 157.7297 | 140.8031 | 78.91797 | 87.58975 | 57.91174 | 121.9117 | 45.88 | 96.87317 | 51.17323 |

**Fv/Fm**

| | WT | soq1 | lcnp | NoM | soq1 NoM | lhcb4 | soq1 lhcb4 | lhcb5 | soq1 lhcb5 | lhcb6 | soq1 lhcb6 |
|---|---|---|---|---|---|---|---|---|---|---|---|
| | 0.15107714 | 0.17597522 | 0.07539966 | 0.05685812 | 0.06114639 | 0.27326912 | 0.20906214 | 0.34158997 | 0.171389 | 0.09064996 | 0.06535456 |
| Rep1 | 0.1403757 | 0.18012365 | 0.09284473 | 0.12036746 | 0.08123955 | 0.25209927 | 0.1567148 | 0.42092309 | 0.22632248 | 0.13861507 | 0.11026689 |
| | 0.18005376 | 0.13426805 | 0.09100347 | 0.12430204 | 0.04120298 | 0.14694367 | 0.09125761 | 0.3096317 | 0.21201649 | 0.11781033 | 0.14775905 |
| | 0.43649149 | 0.28597846 | 0.34401129 | 0.10896927 | 0.10848118 | 0.38067915 | 0.20162572 | 0.42768337 | 0.18558802 | 0.24699545 | 0.25669205 |
| Rep2 | 0.26723973 | 0.2154125 | 0.27730241 | 0.0963601 | 0.15056878 | 0.34435127 | 0.2015114 | 0.45766375 | 0.24590023 | 0.30715546 | 0.1691354 |
| | 0.22515387 | 0.24028043 | 0.14621824 | 0.10168496 | 0.13772066 | 0.24166597 | 0.23688624 | 0.48612932 | 0.16443507 | 0.11247379 | 0.22214568 |
| | 0.20438848 | 0.20849848 | 0.20558484 | -0.0106601 | -0.047534 | 0.11769408 | 0.05643493 | 0.27646744 | 0.08121295 | 0.02070025 | 0.01433636 |
| Rep3 | 0.24860392 | 0.09798443 | 0.26659905 | -0.0149923 | -0.0230619 | 0.17215982 | 0.03475316 | 0.1793637 | 0.03594424 | 0.01220144 | 0.02677103 |
| | 0.21998654 | 0.14968573 | 0.23130267 | -0.0010049 | -0.0145513 | 0.01048799 | 0.03794637 | 0.24859263 | 0.07854664 | 0.03464189 | 0.02357424 |

Figure S6 A, B

Before cold HL

| | Fo | | | | | | | |
|---|---|---|---|---|---|---|---|---|
| | WT | soq1 | lcnp | lhcb6 | soq1 lhcb6 #10 | soq1 lhcb6 #3 | soq1 lhcb6 #7 | soq1 lhcb6-2 |
| | 123.4872 | 154.3988 | 161.7548 | 160.8024 | 142.9612 | 194.9158 | 155.0246 | |
| Rep1 | 155.7503 | 150.2446 | 136.875 | 187.9408 | 147.4419 | 171.333 | 168.4487 | |
| | 150.7979 | 147.1788 | 128.9022 | 179.7049 | 169.7367 | 156.4124 | 169.8092 | |
| | 116.9628 | 141.2043 | 152.8053 | 148.0796 | 172.8325 | 169.2437 | 175.1576 | |
| Rep2 | 141.0592 | 153.5853 | 149.5127 | 154.6825 | 160.4969 | 160.2188 | 166.0691 | |
| | 118.2732 | 126.5726 | 114.763 | 164.4256 | 166.6465 | 167.339 | 161.8968 | |
| | 125.0173 | 133.235 | 110.8314 | 167.3663 | 148.8176 | 176.5419 | 176.3164 | |
| Rep3 | 129.2713 | 126.3507 | 108.8473 | 154.9944 | 162.3323 | 173.0861 | 174.6553 | |
| | 125.4164 | 121.8427 | 113.2804 | 162.8946 | 147.584 | 167.0453 | 175.1995 | |
| | | | | | | | | 161.2516 |
| Rep4 | | | | | | | | 168.1541 |
| | | | | | | | | 170.8913 |

| | Fm | | | | | | | |
|---|---|---|---|---|---|---|---|---|
| | WT | soq1 | lcnp | lhcb6 | soq1 lhcb6 #1 | soq1 lhcb6 #3 | soq1 lhcb6 #7 | soq1 lhcb6-2 |
| | 647.0292 | 726.8297 | 790.095 | 598.3852 | 541.7956 | 655.2469 | 543.274 | |
| Rep1 | 786.5121 | 711.4192 | 692.6709 | 672.4623 | 535.392 | 598.8387 | 588.943 | |
| | 771.5371 | 716.426 | 659.6097 | 658.104 | 604.5258 | 541.2605 | 584.5983 | |
| | 553.6844 | 622.8627 | 669.8469 | 486.9172 | 525.7471 | 577.6996 | 588.4842 | |
| Rep2 | 596.1235 | 667.7153 | 665.5566 | 478.2476 | 508.5317 | 551.1872 | 563.7223 | |
| | 603.6492 | 617.2456 | 611.0687 | 527.1022 | 488.187 | 559.2325 | 567.4683 | |
| | 676.7567 | 659.6047 | 596.8564 | 590.2484 | 536.008 | 598.4214 | 577.0179 | |
| Rep3 | 676.6841 | 631.1421 | 587.0751 | 563.5001 | 569.3051 | 593.2061 | 586.5847 | |
| | 669.192 | 599.4959 | 587.895 | 593.4683 | 502.8651 | 576.4169 | 580.0178 | |
| | | | | | | | | 612.9544 |
| Rep4 | | | | | | | | 642.528 |
| | | | | | | | | 652.2867 |

| | Fv/Fm | | | | | | | |
|---|---|---|---|---|---|---|---|---|
| | WT | soq1 | lcnp | lhcb6 | soq1 lhcb6 #1 | soq1 lhcb6 #3 | soq1 lhcb6 #7 | soq1 lhcb6-2 |
| | 0.80914741 | 0.78757225 | 0.79527171 | 0.73127277 | 0.73613444 | 0.70253076 | 0.71464749 | |
| Rep1 | 0.80197342 | 0.78881003 | 0.80239534 | 0.72051846 | 0.72460945 | 0.71389124 | 0.71398132 | |
| | 0.80454874 | 0.79456524 | 0.80457807 | 0.72693541 | 0.7192234 | 0.71102196 | 0.70952841 | |
| | 0.78875547 | 0.77329787 | 0.77188026 | 0.69588341 | 0.67126305 | 0.70703857 | 0.70235802 | |
| Rep2 | 0.76337252 | 0.76998385 | 0.7753569 | 0.67656398 | 0.68439155 | 0.70932054 | 0.70540619 | |
| | 0.80406965 | 0.79493965 | 0.81219297 | 0.68805746 | 0.65864208 | 0.70077025 | 0.71470336 | |
| | 0.81526995 | 0.79800781 | 0.8143081 | 0.71644769 | 0.72235937 | 0.70498732 | 0.69443513 | |
| Rep3 | 0.80896359 | 0.79980626 | 0.81459391 | 0.72494344 | 0.71485887 | 0.70821929 | 0.7022505 | |
| | 0.81258533 | 0.79675808 | 0.80731185 | 0.72552098 | 0.70651374 | 0.71020055 | 0.69794117 | |
| | | | | | | | | 0.736927 |
| Rep4 | | | | | | | | 0.738293 |
| | | | | | | | | 0.738012 |

After cold HL

| | Fo | | | | | | | |
|---|---|---|---|---|---|---|---|---|
| | WT | soq1 | lcnp | lhcb6 | soq1 lhcb6 #10 | soq1 lhcb6 #3 | soq1 lhcb6 #7 | soq1 lhcb6-2 |
| | 65.15546 | 36.42364 | 130.9242 | 67.92191 | 37.91061 | 41.87854 | 45.20082 | |
| Rep1 | 54.88789 | 34.0242 | 125.7541 | 95.81306 | 37.13733 | 51.51698 | 42.07819 | |
| | 84.20309 | 36.95909 | 119.6135 | 78.40717 | 39.94522 | 40.89272 | 50.36 | |
| | 58.87123 | 41.99948 | 135.5686 | 71.95866 | 45.54273 | 55.05637 | 42.3719 | |
| Rep2 | 72.38818 | 41.15594 | 126.8067 | 106.1265 | 47.34188 | 56.69394 | 58.53963 | |
| | 67.35564 | 40.86131 | 125.9951 | 92.57919 | 40.35568 | 49.60952 | 56.98329 | |
| | 69.29482 | 38.40287 | 127.6591 | 82.51721 | 49.67367 | 50.97556 | 53.12001 | |
| Rep3 | 93.46413 | 47.30857 | 121.7725 | 78.45372 | 48.82257 | 50.43235 | 51.66692 | |
| | 97.78865 | 35.78159 | 121.2464 | 93.49226 | 49.92779 | 56.35828 | 56.25951 | |
| | | | | | | | | 51.39472 |
| Rep4 | | | | | | | | 55.00017 |
| | | | | | | | | 48.15817 |

| | Fm | | | | | | | |
|---|---|---|---|---|---|---|---|---|
| | WT | soq1 | lcnp | lhcb6 | soq1 lhcb6 #1 | soq1 lhcb6 #3 | soq1 lhcb6 #7 | soq1 lhcb6-2 |
| | 80.66553 | 42.73637 | 171.2778 | 70.62472 | 41.79056 | 48.31922 | 46.7517 | |
| Rep1 | 61.50907 | 40.15679 | 194.9785 | 101.2642 | 40.66553 | 54.90961 | 46.36364 | |
| | 109.89 | 44.69635 | 175.5952 | 81.85374 | 44.74249 | 47.56451 | 52.80909 | |
| | 69.34815 | 50.96871 | 146.624 | 79.13197 | 48.72728 | 55.81909 | 44.46317 | |
| Rep2 | 84.20909 | 50.19774 | 139.785 | 123.2045 | 53.20908 | 60.46592 | 60.20087 | |
| | 82.14641 | 47.19857 | 138.609 | 104.9425 | 47.35243 | 55.38995 | 59.36942 | |
| | 87.0963 | 48.51901 | 160.6957 | 83.68497 | 50.6056 | 51.99556 | 56.48546 | |
| Rep3 | 124.3873 | 52.44762 | 166.0381 | 79.53077 | 49.96931 | 54.66222 | 55.64988 | |
| | 125.3679 | 42.08043 | 157.7297 | 96.87317 | 51.17323 | 58.81037 | 57.32889 | |
| | | | | | | | | 57.44771 |
| Rep4 | | | | | | | | 57.7568 |
| | | | | | | | | 55.40075 |

| | Fv/Fm | | | | | | | |
|---|---|---|---|---|---|---|---|---|
| | WT | soq1 | lcnp | lhcb6 | soq1 lhcb6 #1 | soq1 lhcb6 #3 | soq1 lhcb6 #7 | soq1 lhcb6-2 |
| | 0.1922763 | 0.14771329 | 0.23560321 | 0.03827003 | 0.09284274 | 0.13329437 | 0.0331727 | |
| Rep1 | 0.10764559 | 0.15271614 | 0.35503607 | 0.05383087 | 0.08676144 | 0.06178572 | 0.09243127 | |
| | 0.23375112 | 0.1731072 | 0.31881111 | 0.04210645 | 0.10721956 | 0.14026824 | 0.0463763 | |
| | 0.15107714 | 0.17597522 | 0.07539966 | 0.09064996 | 0.06535456 | 0.01366414 | 0.04703376 | |
| Rep2 | 0.1403757 | 0.18012365 | 0.09284473 | 0.13861507 | 0.11026689 | 0.06238192 | 0.02759495 | |
| | 0.18005376 | 0.13426805 | 0.09100347 | 0.11781033 | 0.14775905 | 0.10435882 | 0.04019123 | |
| | 0.20438848 | 0.20849848 | 0.20558484 | 0.02070025 | 0.01433636 | 0.01961706 | 0.05958082 | |
| Rep3 | 0.24860392 | 0.09798443 | 0.26659905 | 0.01220144 | 0.02677103 | 0.07738197 | 0.07157176 | |
| | 0.21998654 | 0.14968573 | 0.23130267 | 0.03464189 | 0.02357424 | 0.04169486 | 0.01865342 | |
| | | | | | | | | 0.105365 |
| Rep4 | | | | | | | | 0.047728 |
| | | | | | | | | 0.130731 |

Figure S6 D

| | NPQ after cold HL | | | | | | | |
|---|---|---|---|---|---|---|---|---|
| | WT | soq1 | lcnp | lhcb6 | soq1 lhcb6 #10 | soq1 lhcb6 #3 | soq1 lhcb6 #7 | soq1 lhcb6-2 |
| | 7.02113617 | 16.0072868 | 3.61294458 | 7.47274439 | 11.9645451 | 12.56079217 | 10.62041166 | |
| Rep1 | 8.7869288 | 16.7160376 | 2.55255015 | 5.64067163 | 12.16574504 | 9.905899714 | 11.70269116 | |
| | 6.02099463 | 15.0287361 | 2.75642216 | 7.03999915 | 12.51122389 | 10.37950333 | 10.07003169 | |
| | 6.98412647 | 11.2204917 | 3.56846696 | 5.15322985 | 9.789584397 | 9.72073 | 11.97744 | |
| Rep2 | 6.07908731 | 12.3017004 | 3.76128769 | 2.88173809 | 8.557235344 | 8.810586 | 8.743791 | |
| | 6.34845503 | 12.0776335 | 3.40857881 | 4.02277152 | 9.309650423 | 9.406525 | 8.769639 | |
| | 6.77021182 | 12.5947683 | 2.71420268 | 6.05321876 | 9.591871255 | 10.11056 | 9.418331 | |
| Rep3 | 4.44013818 | 11.0337605 | 2.53578546 | 6.08530925 | 10.39309508 | 9.083513 | 9.129803 | |
| | 4.33782571 | 13.24643 | 2.72723083 | 5.12624011 | 8.826721901 | 8.50908 | 8.89847 | |
| | | | | | | | | 9.66978 |
| Rep4 | | | | | | | | 10.12472 |
| | | | | | | | | 10.77397 |

Figure S6 C

| | | Rep1 | | | Rep 2 | | | Rep3 | | | Rep1 | | | Rep 2 | |
|---|---|---|---|---|---|---|---|---|---|---|---|---|---|---|---|
| Time (min) | | | | | soq1 | | | | | | | | | lhcb6 | |
| 0 | 0 | 0 | 0 | 0 | 0 | 0 | 0 | 0 | 0 | 0 | 0 | 0 | 0 | 0 | 0 |
| 0.5 | 1.063186 | 1.191234 | 1.158744 | 1.224839 | 1.301047 | 1.265131 | 1.124963 | 1.206055 | 0.6988617 | 1.105121 | 1.079824 | 1.070994 | 1.17715 | 1.203738 | 1.203738 |
| 1.5 | 1.786346 | 1.929185 | 1.800421 | 2.037209 | 2.016932 | 1.987418 | 1.929976 | 1.933771 | 1.511127 | 1.455565 | 1.519212 | 1.520384 | 1.572069 | 1.57909 | 1.57909 |
| 2.5 | 2.163043 | 2.270444 | 2.135829 | 2.333359 | 2.334841 | 2.33917 | 2.256117 | 2.22831 | 1.964716 | 1.491337 | 1.675958 | 1.591368 | 1.713963 | 1.656151 | 1.656151 |
| 3.5 | 2.413918 | 2.513002 | 2.386838 | 2.518786 | 2.520277 | 2.549635 | 2.448503 | 2.42059 | 2.213537 | 1.60356 | 1.769586 | 1.646952 | 1.742474 | 1.624583 | 1.624583 |
| 4.5 | 2.598278 | 2.648236 | 2.545621 | 2.660915 | 2.696492 | 2.696065 | 2.555525 | 2.487034 | 2.412762 | 1.717639 | 1.808964 | 1.691118 | 1.737183 | 1.726743 | 1.726743 |
| 5.5 | 2.799942 | 2.803612 | 2.721513 | 2.787983 | 2.827759 | 2.83123 | 2.680255 | 2.572055 | 2.524489 | 1.837154 | 1.88153 | 1.762632 | 1.827932 | 1.891824 | 1.891824 |
| 6.5 | 2.946712 | 2.938934 | 2.875929 | 2.924256 | 2.960212 | 2.958821 | 2.786379 | 2.665073 | 2.672933 | 1.897642 | 1.961817 | 1.838752 | 1.905055 | 1.98528 | 1.98528 |
| 7.5 | 3.103159 | 3.072721 | 3.025082 | 3.058185 | 3.089149 | 3.081649 | 2.852053 | 2.744411 | 2.777568 | 1.955403 | 1.997757 | 1.888346 | 2.004008 | 2.047874 | 2.047874 |
| 8.5 | 3.219649 | 3.150911 | 3.122336 | 3.144793 | 3.237193 | 3.166776 | 2.954358 | 2.776323 | 2.862653 | 2.005472 | 2.02187 | 1.912949 | 2.065706 | 2.09327 | 2.09327 |
| 9.5 | 3.405527 | 3.266645 | 3.269215 | 3.222458 | 3.330669 | 3.293508 | 3.014436 | 2.854937 | 3.00923 | 2.055338 | 2.098358 | 1.977751 | 2.103851 | 2.171118 | 2.171118 |
| 10.6 | 3.477389 | 3.349118 | 3.347637 | 3.381299 | 3.40467 | 3.360742 | 3.077111 | 2.96295 | 3.123558 | 2.077587 | 2.103687 | 1.993457 | 2.158344 | 2.240741 | 2.240741 |
| 10.8 | 2.564016 | 2.433672 | 2.589018 | 2.379384 | 2.509716 | 2.388426 | 2.117651 | 1.96878 | 2.257405 | 0.9233678 | 0.9468036 | 0.7947398 | 1.22437 | 1.26487 | 1.26487 |
| 11.1 | 2.247926 | 2.056932 | 2.237863 | 1.986191 | 2.10394 | 1.957702 | 1.60671 | 1.516869 | 1.917616 | 0.6997752 | 0.7337279 | 0.6349944 | 0.8518769 | 0.934023 | 0.934023 |
| 12.2 | 1.922405 | 1.650913 | 1.825928 | 1.57405 | 1.691512 | 1.543688 | 1.182261 | 1.102058 | 1.639193 | 0.5423703 | 0.5882708 | 0.5339211 | 0.5358641 | 0.5738794 | 0.5738794 |
| 13.2 | 1.819382 | 1.533584 | 1.719346 | 1.492039 | 1.629279 | 1.481727 | 1.11563 | 1.026911 | 1.586229 | 0.4916948 | 0.5320387 | 0.4964748 | 0.5049316 | 0.5382055 | 0.5382055 |
| 14.2 | 1.767193 | 1.457647 | 1.653982 | 1.447914 | 1.595793 | 1.410528 | 1.053343 | 1.0009 | 1.530871 | 0.4554378 | 0.4912202 | 0.4672378 | 0.4724644 | 0.513692 | 0.513692 |
| 15.2 | 1.723704 | 1.376089 | 1.598304 | 1.417412 | 1.556446 | 1.391772 | 1.010123 | 0.9744635 | 1.51728 | 0.4254562 | 0.4658274 | 0.4401656 | 0.4419422 | 0.4954433 | 0.4954433 |
| 16.2 | 1.706335 | 1.352527 | 1.567056 | 1.391204 | 1.550996 | 1.372732 | 0.9648067 | 0.9497319 | 1.499642 | 0.3949189 | 0.4540262 | 0.4270556 | 0.4157187 | 0.4743215 | 0.4743215 |
| 17.2 | 1.660604 | 1.328819 | 1.545678 | 1.359806 | 1.513002 | 1.310423 | 0.9247051 | 0.9338358 | 1.48827 | 0.354177 | 0.4362632 | 0.4197216 | 0.3983363 | 0.4595424 | 0.4595424 |
| 18.2 | 1.604056 | 1.293709 | 1.510563 | 1.344191 | 1.486824 | 1.281881 | 0.9028957 | 0.9056707 | 1.477147 | 0.3129643 | 0.4211427 | 0.4071154 | 0.3750342 | 0.4464597 | 0.4464597 |
| 19.2 | 1.599033 | 1.274783 | 1.500767 | 1.309569 | 1.437678 | 1.23367 | 0.887763 | 0.8829742 | 1.446843 | 0.2897761 | 0.3910247 | 0.3899979 | 0.3466754 | 0.4097908 | 0.4097908 |
| 20.2 | 1.591102 | 1.277943 | 1.493261 | 1.262029 | 1.398531 | 1.20078 | 0.8675089 | 0.8228216 | 1.455215 | 0.2915463 | 0.3705188 | 0.3815178 | 0.3332153 | 0.4064571 | 0.4064571 |
| 21.2 | 1.575054 | 1.252931 | 1.479038 | 1.240986 | 1.368624 | 1.165192 | 0.8542453 | 0.7891608 | 1.442521 | 0.2898108 | 0.3502428 | 0.3667488 | 0.3103442 | 0.3845089 | 0.3845089 |

| | | Rep3 | | | | | | Rep1 | | | | | | | |
|---|---|---|---|---|---|---|---|---|---|---|---|---|---|---|---|
| Time (min) | | | | | soq1 lhcb6 #3 | | | soq1 lhcb6 #7 | | | soq1 lhcb6 #10 | | | soq1 lhcb6 #3 | |
| 0 | 0 | 0 | 0 | 0 | 0 | 0 | 0 | 0 | 0 | 0 | 0 | 0 | 0 | 0 | 0 |
| 0.5 | 1.03023 | 1.13287 | 1.055329 | 1.130456 | 1.107622 | 1.110163 | 1.145975 | 1.129651 | 1.144499 | 1.07899 | 1.134804 | 1.129794 | 1.135881 | 1.254734 | 1.122385 |
| 1.5 | 1.448526 | 1.634596 | 1.522169 | 1.54145 | 1.578493 | 1.589499 | 1.612606 | 1.559787 | 1.608536 | 1.435823 | 1.517833 | 1.512542 | 1.498869 | 1.676877 | 1.523395 |
| 2.5 | 1.633357 | 1.820499 | 1.734674 | 1.72779 | 1.757773 | 1.77655 | 1.819588 | 1.755161 | 1.821547 | 1.550559 | 1.677298 | 1.693336 | 1.669103 | 1.859016 | 1.681423 |
| 3.5 | 1.781101 | 1.941058 | 1.86334 | 1.885215 | 1.941801 | 1.953024 | 2.012461 | 1.945036 | 2.00423 | 1.585558 | 1.710911 | 1.734926 | 1.744277 | 1.91227 | 1.767104 |
| 4.5 | 1.833746 | 1.999237 | 1.930458 | 1.991027 | 2.069232 | 2.069777 | 2.133769 | 2.076376 | 2.121778 | 1.617193 | 1.726405 | 1.724304 | 1.775503 | 1.969401 | 1.787449 |
| 5.5 | 1.929106 | 2.092353 | 1.940206 | 2.141536 | 2.209402 | 2.21367 | 2.313283 | 2.223544 | 2.278696 | 1.678969 | 1.829077 | 1.823996 | 1.879819 | 2.104193 | 1.871905 |
| 6.5 | 1.952541 | 2.167742 | 1.998291 | 2.278182 | 2.339773 | 2.335654 | 2.439013 | 2.343739 | 2.384816 | 1.736844 | 1.905608 | 1.948844 | 2.015697 | 2.276218 | 2.026553 |
| 7.5 | 2.045233 | 2.270705 | 2.071043 | 2.375228 | 2.4549 | 2.441935 | 2.564123 | 2.460754 | 2.519933 | 1.783576 | 2.039041 | 2.077098 | 2.143239 | 2.392316 | 2.133405 |
| 8.5 | 2.10161 | 2.299614 | 2.18132 | 2.445427 | 2.484548 | 2.514722 | 2.628233 | 2.548491 | 2.585798 | 1.825484 | 2.104627 | 2.129067 | 2.213948 | 2.433942 | 2.219635 |
| 9.5 | 2.185414 | 2.391709 | 2.269455 | 2.555237 | 2.604964 | 2.631229 | 2.770106 | 2.678207 | 2.716536 | 1.86933 | 2.112905 | 2.211997 | 2.284644 | 2.556216 | 2.299205 |
| 10.6 | 2.241234 | 2.42429 | 2.3436 | 2.612006 | 2.674541 | 2.681251 | 2.855308 | 2.704414 | 2.795709 | 1.91592 | 2.219188 | 2.279274 | 2.373855 | 2.623475 | 2.390335 |
| 10.8 | 1.408226 | 1.462292 | 1.521134 | 1.577714 | 1.662385 | 1.769382 | 1.968184 | 1.700608 | 1.937794 | 0.8605296 | 1.219214 | 1.365229 | 1.472092 | 1.669532 | 1.477896 |
| 11.1 | 1.003027 | 1.064249 | 1.059169 | 1.406355 | 1.445204 | 1.523667 | 1.723611 | 1.510208 | 1.684987 | 0.6359209 | 0.9886379 | 1.097723 | 1.24675 | 1.389862 | 1.249731 |
| 12.2 | 0.6495774 | 0.6964722 | 0.6941339 | 1.304376 | 1.285029 | 1.386099 | 1.496894 | 1.398498 | 1.417903 | 0.4794698 | 0.788946 | 0.9019937 | 1.096367 | 1.113472 | 1.081045 |
| 13.2 | 0.5672405 | 0.6242517 | 0.6186139 | 1.23968 | 1.212747 | 1.338099 | 1.42621 | 1.345671 | 1.330006 | 0.4491006 | 0.7527227 | 0.8653218 | 1.060124 | 1.057873 | 1.047033 |
| 14.2 | 0.5351492 | 0.5889768 | 0.5631541 | 1.190407 | 1.159103 | 1.285763 | 1.371332 | 1.300162 | 1.2698 | 0.4188424 | 0.696239 | 0.8155784 | 1.028651 | 0.9993374 | 1.0026 |
| 15.2 | 0.5205435 | 0.5574208 | 0.537792 | 1.14096 | 1.099485 | 1.240926 | 1.315016 | 1.247952 | 1.203782 | 0.3887161 | 0.6654484 | 0.7716618 | 0.9851768 | 0.9644058 | 0.9581981 |
| 16.2 | 0.5040697 | 0.5255495 | 0.5117595 | 1.114012 | 1.070183 | 1.222603 | 1.282234 | 1.237734 | 1.193905 | 0.3637584 | 0.6417589 | 0.7513332 | 0.9716057 | 0.9352993 | 0.9476616 |
| 17.2 | 0.4750784 | 0.4905591 | 0.4890141 | 1.091338 | 1.026692 | 1.203924 | 1.250512 | 1.229776 | 1.172245 | 0.3529474 | 0.6342018 | 0.730862 | 0.9594356 | 0.9142656 | 0.9229431 |
| 18.2 | 0.4673035 | 0.4730054 | 0.473807 | 1.045496 | 0.9718561 | 1.173045 | 1.223239 | 1.202275 | 1.14561 | 0.3456587 | 0.6148348 | 0.7170514 | 0.9388666 | 0.8901792 | 0.9218431 |
| 19.2 | 0.4688929 | 0.4645061 | 0.4686438 | 1.01402 | 0.9378964 | 1.138516 | 1.198723 | 1.179858 | 1.123195 | 0.3241644 | 0.584254 | 0.6832479 | 0.9235604 | 0.8380162 | 0.888053 |
| 20.2 | 0.4649682 | 0.4624561 | 0.464747 | 1.012462 | 0.9227352 | 1.12191 | 1.181973 | 1.171509 | 1.124641 | 0.3230753 | 0.5615965 | 0.6670401 | 0.8985527 | 0.8256932 | 0.8886297 |
| 21.2 | 0.46817 | 0.4665601 | 0.4728767 | 0.995186 | 0.9107713 | 1.102622 | 1.169164 | 1.151851 | 1.114056 | 0.307736 | 0.537238 | 0.6406916 | 0.8898328 | 0.7984088 | 0.8669997 |

| | | Rep2 | | | | | | | | | Rep3 | | | | |
|---|---|---|---|---|---|---|---|---|---|---|---|---|---|---|---|
| Time (min) | | soq1 lhcb6 #7 | | | soq1 lhcb6 #10 | | | soq1 lhcb6 #3 | | | soq1 lhcb6 #7 | | | soq1 lhcb6 #10 | |
| 0 | 0 | 0 | 0 | 0 | 0 | 0 | 0 | 0 | 0 | 0 | 0 | 0 | 0 | 0 | 0 |
| 0.5 | 1.171091 | 1.171091 | 1.240394 | 1.044148 | 1.070311 | 0.993634 | 1.14625 | 1.085958 | 0.9100392 | 1.001182 | 1.078229 | 1.082899 | 1.023275 | 1.093111 | 0.987899 |
| 1.5 | 1.505973 | 1.505973 | 1.69582 | 1.548109 | 1.578605 | 1.46291 | 1.597685 | 1.412129 | 1.049212 | 1.396206 | 1.365092 | 1.337288 | 1.352914 | 1.319864 | 1.264603 |
| 2.5 | 1.6178 | 1.6178 | 1.892009 | 1.74049 | 1.770805 | 1.557303 | 1.747324 | 1.529986 | 1.272732 | 1.556404 | 1.354648 | 1.430333 | 1.331584 | 1.277929 | 1.246946 |
| 3.5 | 1.647757 | 1.647757 | 1.973748 | 1.889055 | 1.88271 | 1.530104 | 1.870441 | 1.722079 | 1.462716 | 1.739329 | 1.516039 | 1.608338 | 1.49657 | 1.432122 | 1.345757 |
| 4.5 | 1.609907 | 1.609907 | 1.966243 | 1.970526 | 1.967356 | 1.520282 | 2.047408 | 1.878993 | 1.594758 | 1.859035 | 1.715404 | 1.738281 | 1.723874 | 1.661791 | 1.513009 |
| 5.5 | 1.727957 | 1.727957 | 2.052898 | 2.002532 | 2.009118 | 1.658467 | 2.178365 | 1.993336 | 1.701086 | 1.955996 | 1.819558 | 1.826162 | 1.934031 | 1.799082 | 1.604178 |
| 6.5 | 1.858249 | 1.858249 | 2.21321 | 1.991942 | 2.070822 | 1.807302 | 2.256456 | 2.094841 | 1.79987 | 2.015976 | 1.880463 | 1.888224 | 2.069013 | 1.890017 | 1.720651 |
| 7.5 | 1.958712 | 1.958712 | 2.356184 | 2.099051 | 2.14054 | 1.898834 | 2.347577 | 2.160186 | 1.889841 | 2.083393 | 1.957175 | 1.980774 | 2.099277 | 1.966269 | 1.808814 |
| 8.5 | 2.019161 | 2.019161 | 2.474462 | 2.173495 | 2.20464 | 1.956121 | 2.447003 | 2.243419 | 2.000836 | 2.216375 | 2.063074 | 2.104727 | 2.24815 | 2.055052 | 1.867914 |
| 9.5 | 2.109671 | 2.109671 | 2.523516 | 2.23068 | 2.253233 | 2.016986 | 2.5009 | 2.301762 | 2.067428 | 2.24773 | 2.101234 | 2.154575 | 2.296921 | 2.24918 | 2.001918 |
| 10.6 | 2.219528 | 2.219528 | 2.570685 | 2.329309 | 2.349583 | 2.038165 | 2.58047 | 2.365761 | 2.126379 | 2.320749 | 2.165546 | 2.229421 | 2.443382 | 2.360485 | 2.118852 |
| 10.8 | 1.333399 | 1.333399 | 1.715792 | 1.41653 | 1.44575 | 1.032672 | 1.473578 | 1.301149 | 1.129433 | 1.129361 | 0.9816718 | 1.16707 | 1.45271 | 1.222604 | 1.046951 |
| 11.1 | 1.08681 | 1.08681 | 1.39791 | 0.9592122 | 1.018022 | 0.7693173 | 1.131924 | 1.085391 | 0.98975 | 0.9219914 | 0.7175688 | 0.9605054 | 1.280557 | 0.981057 | 0.868472 |
| 12.2 | 0.9022833 | 0.9022833 | 1.107066 | 0.6637331 | 0.6679443 | 0.6005699 | 0.876062 | 0.9501874 | 0.8868043 | 0.7826514 | 0.5051513 | 0.7684445 | 1.156821 | 0.835491 | 0.7278119 |
| 13.2 | 0.8689292 | 0.8689292 | 1.066571 | 0.5642675 | 0.569892 | 0.5576511 | 0.7645914 | 0.8426856 | 0.8161576 | 0.7191051 | 0.4213142 | 0.6527264 | 1.1208 | 0.8053256 | 0.6653565 |
| 14.2 | 0.8297527 | 0.8297527 | 1.004821 | 0.5117385 | 0.5297587 | 0.5181706 | 0.6563457 | 0.7440909 | 0.7139289 | 0.6653856 | 0.3680604 | 0.5560597 | 1.071422 | 0.7672749 | 0.6187866 |
| 15.2 | 0.8002713 | 0.8002713 | 0.9694261 | 0.4871825 | 0.504633 | 0.4944287 | 0.6115463 | 0.7131395 | 0.6492357 | 0.6317204 | 0.3440974 | 0.5112641 | 1.031726 | 0.7437063 | 0.5951264 |
| 16.2 | 0.7827294 | 0.7827294 | 0.9335774 | 0.4634939 | 0.4841888 | 0.4885107 | 0.5896118 | 0.7049024 | 0.6234184 | 0.6035876 | 0.3290714 | 0.4899235 | 1.018774 | 0.7174884 | 0.5675073 |
| 17.2 | 0.7674704 | 0.7674704 | 0.9170849 | 0.4436414 | 0.4547637 | 0.4740174 | 0.5674372 | 0.6895515 | 0.6060153 | 0.5691135 | 0.3146091 | 0.4730159 | 0.97512 | 0.6894713 | 0.5133461 |
| 18.2 | 0.7411156 | 0.7411156 | 0.8759612 | 0.4356408 | 0.4530922 | 0.4671387 | 0.5518435 | 0.6806712 | 0.5971417 | 0.5359193 | 0.3090837 | 0.4534455 | 0.97195 | 0.6924344 | 0.5221265 |
| 19.2 | 0.7226551 | 0.7226551 | 0.8259247 | 0.4328583 | 0.4433134 | 0.4599631 | 0.5497561 | 0.682241 | 0.5958406 | 0.512384 | 0.3128052 | 0.4496209 | 0.937774 | 0.675668 | 0.5004642 |
| 20.2 | 0.7132171 | 0.7132171 | 0.803005 | 0.4351777 | 0.4359263 | 0.4565065 | 0.5480962 | 0.6856412 | 0.592185 | 0.4866837 | 0.3159371 | 0.444095 | 0.892662 | 0.6336532 | 0.4657488 |
| 21.2 | 0.7033311 | 0.7033311 | 0.8005363 | 0.4355516 | 0.4311733 | 0.4587948 | 0.5542892 | 0.6867381 | 0.5970509 | 0.4687299 | 0.3278216 | 0.4414768 | 0.86129 | 0.6293389 | 0.4429923 |
